# Broadly applicable and accurate protein design by integrating structure prediction networks and diffusion generative models

**DOI:** 10.1101/2022.12.09.519842

**Authors:** Joseph L. Watson, David Juergens, Nathaniel R. Bennett, Brian L. Trippe, Jason Yim, Helen E. Eisenach, Woody Ahern, Andrew J. Borst, Robert J. Ragotte, Lukas F. Milles, Basile I. M. Wicky, Nikita Hanikel, Samuel J. Pellock, Alexis Courbet, William Sheffler, Jue Wang, Preetham Venkatesh, Isaac Sappington, Susana Vázquez Torres, Anna Lauko, Valentin De Bortoli, Emile Mathieu, Regina Barzilay, Tommi S. Jaakkola, Frank DiMaio, Minkyung Baek, David Baker

**Author notes:** Equal contribution (Joseph L. Watson, David Juergens, Nathaniel R. Bennett, Brian L. Trippe, Jason Yim, Helen E. Eisenach, Woody Ahern).

## Abstract

There has been considerable recent progress in designing new proteins using deep learning methods^1–9^. Despite this progress, a general deep learning framework for protein design that enables solution of a wide range of design challenges, including *de novo* binder design and design of higher order symmetric architectures, has yet to be described. Diffusion models^10,11^ have had considerable success in image and language generative modeling but limited success when applied to protein modeling, likely due to the complexity of protein backbone geometry and sequence-structure relationships. Here we show that by fine tuning the RoseTTAFold structure prediction network on protein structure denoising tasks, we obtain a generative model of protein backbones that achieves outstanding performance on unconditional and topology-constrained protein monomer design, protein binder design, symmetric oligomer design, enzyme active site scaffolding, and symmetric motif scaffolding for therapeutic and metal-binding protein design. We demonstrate the power and generality of the method, called RoseTTAFold Diffusion (RF*diffusion*), by experimentally characterizing the structures and functions of hundreds of new designs. In a manner analogous to networks which produce images from user-specified inputs, RF*diffusion* enables the design of diverse, complex, functional proteins from simple molecular specifications.

## Main

De novo protein design seeks to generate proteins with specified structural and/or functional properties, for example making a binding interaction with a given target^12^, folding into a particular topology^13^, or stabilizing a desired functional “motif” (geometries and amino acid identities that produce a desired activity)^4^. Denoising diffusion probabilistic models (DDPMs), a powerful class of machine learning models recently demonstrated to generate novel photorealistic images in response to text prompts^14,15^, have several properties well-suited to protein design. First, DDPMs generate highly diverse outputs – DDPMs are trained to denoise data (for instance images or text) that have been corrupted with Gaussian noise; by learning to stochastically reverse this corruption, diverse outputs closely resembling the training data are generated. Second, DDPMs can be guided at each step of the iterative generation process towards specific design objectives through provision of conditioning information. Third, for almost all protein design applications it is necessary to explicitly model 3D structure; SE(3)-equivariant DDPMs are able to do this in a representation-frame independent manner. Recent work has adapted DDPMs for protein monomer design by conditioning on small protein “motifs”^5,9^ or on secondary structure and block-adjacency (“fold”) information^8^. While promising, these attempts have shown limited success in generating sequences that fold to the intended structures *in silico*^5,16^, likely due to the limited ability of the denoising networks to generate realistic protein backbones, and have not been tested experimentally.

We reasoned that improved diffusion models for protein design could be developed by taking advantage of the deep understanding of protein structure implicit in powerful structure prediction methods like AlphaFold2 (AF2) and RoseTTAFold (RF). RF has properties particularly well suited for use in a protein design DDPM (Fig. 1A). First, RF can generate protein structures with very high precision, and in our previous work we demonstrated considerable success in accurately scaffolding motifs following fine tuning of RF for protein design (“RF_joint_ Inpainting”)^4^. Second, RF operates on a rigid-frame representation of residues with rotational and translational equivariance. Third, the RF architecture enables conditioning on design specifications at three different levels: individual residue properties, pairwise distances and orientations between residues, and 3D coordinates. In RF_joint_ Inpainting, we fine-tuned RF to design protein scaffolds in a *single* step. Experimental characterization showed that the method can scaffold a wide range of protein functional motifs with atomic accuracy^17^, but the approach fails on minimalist site descriptions that do not sufficiently constrain the overall fold, and because it is deterministic, can produce only a limited diversity of designs for a given problem. We reasoned that by instead fine-tuning RoseTTAFold as the denoising network in a generative diffusion model, we could overcome both problems: because the starting point is random noise, each denoising trajectory yields a different solution, and because structure is built up progressively through many denoising iterations, little to no starting structural information should be required.

**Figure 1:**
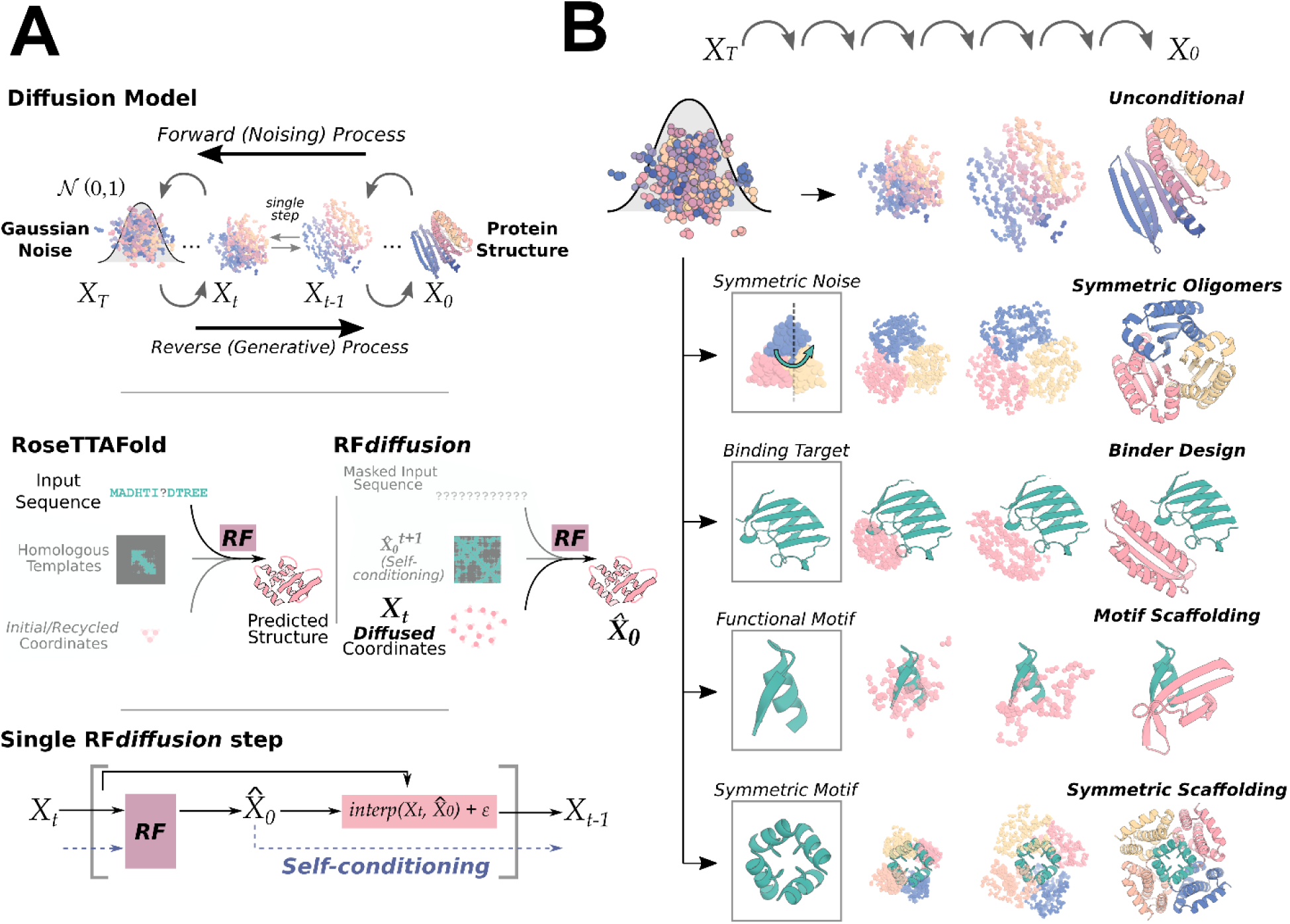
RF*diffusion* is a denoising diffusion probabilistic model with RoseTTAFold fined-tuned as the denoising network. **A)** Top panel: Diffusion models for proteins are trained to recover structures of proteins corrupted with noise, and generate new structures by reversing the corruption process through iterative denoising of initially random noise *X*_*T*_ into a realistic structure *X* _0_. Middle panel: RoseTTAFold (RF, left) can be fine-tuned as the denoising network in a DDPM. RF*diffusion* (right) is trained from a *pre-trained* RF network with minimal architectural changes. While in RF, the primary input to the model is sequence, in RF*diffusion*, the primary input is diffused residue frames. In both cases, the model predicts final 3D coordinates directly (denoted 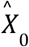 in RF*diffusion*). In RF*diffusion*, the model receives its previous prediction as a template input (“self-conditioning”, see Methods 2.4). Bottom panel: At each timestep “t” of a design trajectory (typically 200 steps), RF*diffusion* takes *X*_*t*_ and 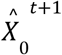 from the previous step and then predicts an updated *X*_0_ structure 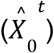. The coordinate input to the model at the next time step (*X*_*t*−1_) is generated by a noisy interpolation toward 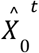. B) RF*diffusion* is of broad applicability to protein design. RF*diffusion* generates protein structures either without additional input (top row), or by conditioning on: symmetric inputs to design symmetric oligomers (second row); a binding target (third row); protein functional motifs (fourth row); symmetric functional motifs to design symmetric oligomers scaffolds (bottom row). In each case random noise, along with conditioning information, is input to RF*diffusion*, which iteratively refines that noise until a final protein structure is designed.

We construct a RoseTTAFold-based diffusion model, RF*diffusion*, using the RF frame representation which comprises a C_ɑ_ coordinate and N-C_ɑ_-C rigid orientation for each residue. We generate training inputs by simulating the noising process for a random number of steps (up to 200) on structures sampled from the Protein Data Bank (PDB)^18^. For translations, we perturb C_ɑ_ coordinates with 3D Gaussian noise. For residue orientations, we use Brownian motion on the manifold of rotation matrices (building on refs [^19,20^]). To enable RF*diffusion* to learn to reverse each step of the noising process, we train the model by minimizing a mean squared error (MSE) loss between frame predictions and the *true* protein structure (without alignment), averaged across all residues (Methods 2.5). This loss drives denoising trajectories to match the data distribution at each timestep and hence to converge on structures of designable protein backbones (Fig. S1A). MSE contrasts to the loss used in RF structure prediction training (“frame aligned point error”, FAPE) in that unlike FAPE, MSE loss is not invariant to the global reference frame and therefore promotes continuity of the global coordinate frame between timesteps (Methods 2.5). While in this study we use RoseTTAFold as the basis for the denoising network architecture, other SE(3)-equivariant structure prediction networks (AF2^21^, OmegaFold^22^, ESMFold^23^) could in principle be substituted into an analogous DDPM.

To generate a new protein backbone, we first initialize random residue frames and RF*diffusion* makes a denoised prediction. Each residue frame is updated by taking a step in the direction of this prediction with some noise added to generate the input to the next step. The nature of the noise added and the size of this reverse step is chosen such that the denoising process matches the distribution of the noising process (Methods 2.2-2.3, Figure S1A). RF*diffusion* initially seeks to match the full breadth of possible protein structures compatible with the purely random frames with which it is initialized, and hence the denoised structures do not initially appear protein-like (Fig. 2A left). However, through many such steps, the breadth of possible protein structures from which the input could have arisen narrows, and RF*diffusion* predictions come to closely resemble protein structures (Fig. 2A right). We use the ProteinMPNN network^1^ to subsequently design sequences encoding these structures. We also considered simultaneously designing structure and sequence within RF*diffusion*, but given the excellent performance of combining ProteinMPNN with the diffusion of structure alone, we did not extensively explore this possibility.

**Figure 2:**
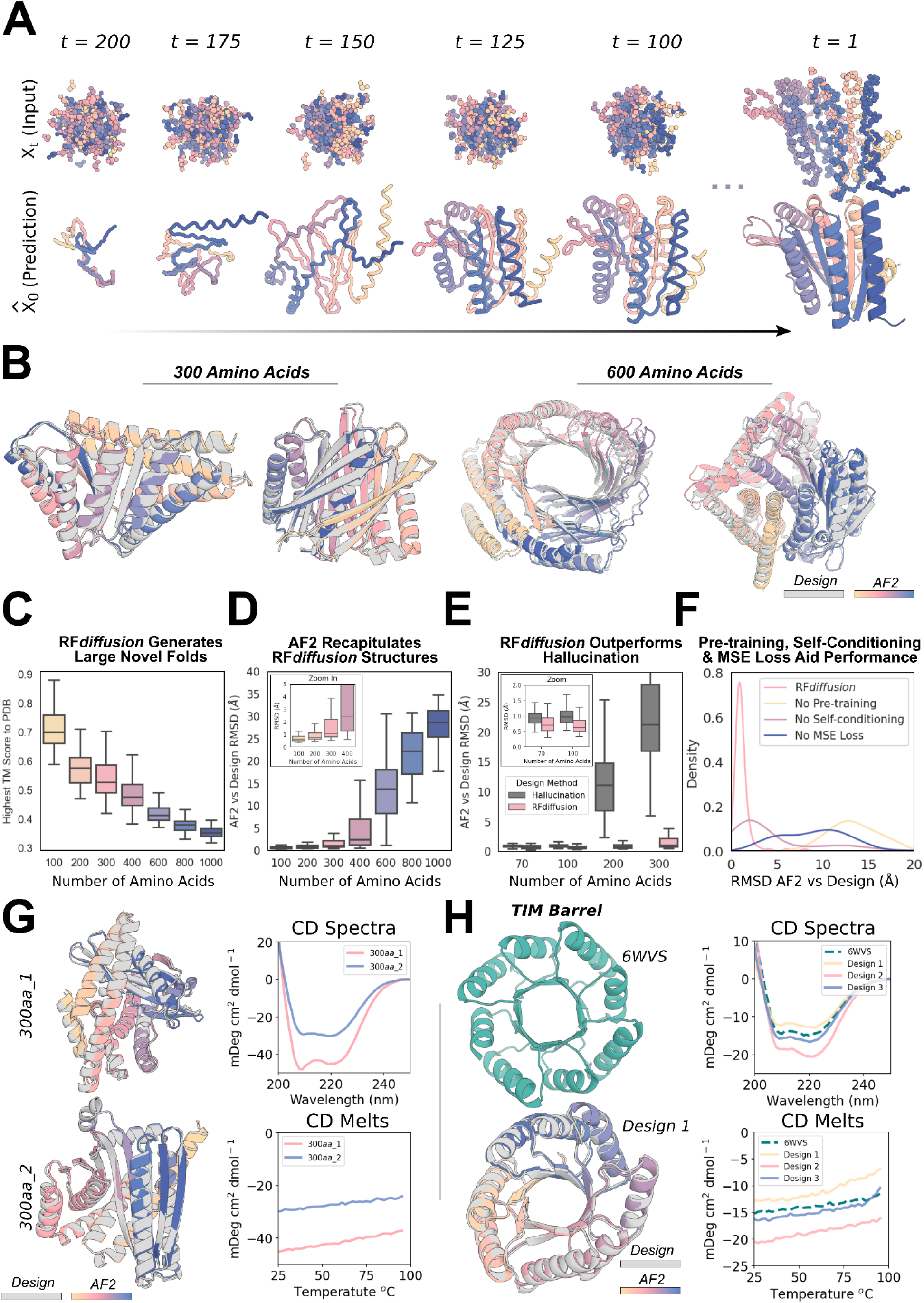
Outstanding performance of RF*diffusion* for monomer generation. **A)** An example trajectory of an unconditional 300 amino acid design, depicting the input to the model (X_t_) and the corresponding 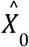 prediction. At early timesteps (high t), 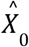 predictions bear little resemblance to a protein, but are gradually refined into a protein structure. **B)** RF*diffusion* can generate new monomeric proteins of different lengths (left: 300, right: 600) with no conditioning information. Gray=design model; colors= AlphaFold2 (AF2) prediction. RMSD AF2 vs design (Å), left to right: 0.90, 0.98, 1.15, 1.67. **C)** Unconditional designs from RF*diffusion* are novel and not present in the training set as quantified by highest TM score to the protein data bank (PDB). Designs are increasingly novel with increasing length. **D)** Unconditional samples are closely re-predicted by AF2. Beyond 400 amino acids, the recapitulation by AF2 deteriorates. **E)** RF*diffusion* significantly outperforms Hallucination (with RoseTTAFold) at unconditional monomer generation (two-way ANOVA & Tukey’s test, *p<0*.*001*). While Hallucination successfully generates designs up to 100 amino acids in length, success rates rapidly deteriorate beyond this length. **F)** Ablating pre-training (by starting from untrained RF), self-conditioning, or MSE losses (by training with FAPE) each dramatically decrease the performance of RF*diffusion*. RMSD between design and AF2 is shown, for the motif-scaffolding problem “5TPN” (see Methods 5.2). **G)** Two example 300 amino acid proteins that expressed as soluble monomers. Designs (gray) overlaid with AF2 predictions (colors) are shown on the left, alongside CD spectra (top) and melt curves (bottom) on the right. The designs are highly thermostable. **H)** RF*diffusion* can condition on fold information. An example TIM barrel is shown (bottom left), conditioned on the secondary structure and block-adjacency of a previously designed TIM barrel, PDB: 6WVS (top left). Designs have very similar CD spectra to 6WVS (top right), and are highly thermostable (bottom right).

Fig. 1A highlights the similarities between RoseTTAFold structure prediction and an RF*diffusion* denoising step: in both cases, the networks transform coordinates into a predicted structure, conditioned on inputs to the model. In RoseTTAFold, sequence is the primary input, with additional structural information provided as templates and initial coordinates to the model. In RF*diffusion*, the primary input is the noised coordinates from the previous step. For design tasks, we optionally provide a range of auxiliary conditioning information, including partial sequence, fold information, or fixed functional motif coordinates (see Methods 3).

We explored two different strategies for training RF*diffusion*: 1) in a manner akin to “canonical” diffusion models, with predictions at each timestep independent of predictions at previous timesteps (as in previous work^5,8,9,16^), and 2) with self-conditioning^24^, where the model can condition on previous predictions between timesteps (Fig. 1A bottom row, Methods 2.4). The latter strategy was inspired by the success of “recycling” in AF2, which is also central to the more recent RF model used here (Methods 1). Self-conditioning within RF*diffusion* dramatically improved performance on *in silico* benchmarks encompassing both conditional and unconditional protein design tasks (Fig. S2E). Increased coherence of predictions within self-conditioned trajectories may, at least in part, explain these performance increases (Fig. S2H). Fine-tuning RF*diffusion* from pre-trained RF weights was far more successful than training for an equivalent length of time from untrained weights (Fig. S2F) and the MSE loss was also crucial (Fig. S2D). For all *in silico* benchmarks in this paper, we use the AF2 structure prediction network^21^ for validation and define an *in silico* “success” as an RF*diffusion* output for which the AF2 structure predicted from a single sequence is (1) of high confidence (mean predicted aligned error, pAE, < 5), (2) globally within 2Å backbone-RMSD of the designed structure, and (3) within 1Å backbone-RMSD on any scaffolded functional-site. This definition of success is significantly more stringent than those described elsewhere (refs [^5,8,16,25^], Fig. S3A-B) but is a good predictor of experimental success^4,7,26^.

### Unconditional protein monomer generation

Physically-based protein design methodologies have struggled in unconstrained generation of diverse protein monomers due to the difficulty of sampling on the very large and rugged conformational landscape^27^, and overcoming this limitation has been a primary test of deep learning based protein design approaches^5,6,8,16,28,29^. As illustrated in Fig. 2B-D, Fig. S4B-C, starting from random noise, RF*diffusion* can readily generate elaborate protein structures with little overall structural similarity to any known protein structures, indicating considerable generalization beyond the PDB training set. The designs are diverse (Fig. S4A), spanning a wide range of alpha-, beta- and mixed alpha-beta-topologies, with AF2 and ESMFold (Fig. S2B-C, Fig. S3A) predictions very close to the design structure models for *de novo* designs with as many as 600 residues. RF*diffusion* generates plausible structures for even very large proteins, but these are difficult to validate *in silico* as they are likely beyond the single sequence prediction capabilities of AF2 and ESMFold. The quality and diversity of designs that are sampled is inherent to the model, and does not require *any* auxiliary conditioning input (for example secondary structure information^8^). Characterization of two of these 300 amino acid proteins is shown in Figure 2G, demonstrating circular dichroism (CD) spectra consistent with the mixed alpha-beta topologies of the two designs, and CD melts showing that designs are extremely thermostable. RF*diffusion* strongly outperforms Hallucination^4^ (Fig. 2E), the only previously described deep learning method for unconditional protein structure generation that has been experimentally validated^6^. Hallucination uses Monte Carlo search or gradient descent to identify sequences predicted to fold into stable structures; in contrast to RF*diffusion*, Hallucination success rates deteriorate beyond 100 amino acids. RF*diffusion* is also more compute efficient than unconstrained Hallucination, requiring ∼2.5 minutes on an NVIDIA RTX A4000 GPU to generate a 100 residue structure compared to ∼8.5 minutes for Hallucination. The computational efficiency of RF*diffusion* can be further improved by taking larger steps at inference time, and by truncating trajectories early - an advantage of predicting the *final* structure at each timestep (Fig. S3C-D). For design problems where a particular fold or architecture is desired (such as TIM barrels or cavity-containing NTF2s for small molecule binder and enzyme design^30,31^), we further fine-tuned RF*diffusion* to condition on secondary structure and/or fold information, enabling rapid and accurate generation of diverse designs with the desired topologies (Fig. 2H, Fig. S5). *In silico* success rates were 42.5% and 54.1% for TIM barrels and NTF2 folds respectively (Fig. S5G), and experimental characterization of 11 TIM barrel designs indicated that at least 9 designs were soluble, thermostable, and had circular dichroism (CD) spectra consistent with the design model (Fig. 2H, Fig. S5D-F).

### Design of higher order oligomers

There is considerable interest in designing symmetric oligomers, which can serve as vaccine platforms^32^, delivery vehicles^33^, and catalysts^34^. Cyclic oligomers have been designed using structure prediction networks with an adaptation of Hallucination that searches for sequences predicted to fold to the desired cyclic symmetry, but this approach fails for higher order dihedral, tetrahedral, octahedral, and icosahedral symmetries, likely in part because of the much lower representation of such structures in the PDB^7^.

We set out to generalize RF*diffusion* to create symmetric oligomeric structures with any specified point group symmetry. Given a specification of a point group symmetry for an oligomer with N chains, and the monomer chain length, we generate random starting residue frames for a single monomer subunit as in the unconditional generation case, and then generate N-1 copies of this starting point arranged with the specified point group symmetry. Because RF*diffusion* exhibits equivariance (inherited from RF) with respect to rotation and relabelings of chains, symmetry is largely maintained in the denoising predictions; although we explicitly re-symmetrize at each step, this changes the structures only slightly (Compare gray and colored chains in Fig. S6A, Methods Proposition 2). For octahedral and icosahedral architectures, we explicitly model only the smallest subset of monomers required to generate the full assembly (e.g. for icosahedra, the subunits at the five-fold, three-fold, and two-fold symmetry axes) to reduce the computational cost and memory footprint.

Despite not being trained on symmetric inputs, RF*diffusion* is able to generate symmetric oligomers with high *in silico* success rates (Fig. S6B), particularly when guided by an auxiliary inter- and intra-chain contact potential (Fig. S6C). As illustrated in Fig. 3 and Fig. S6E, RF*diffusion*-generated designs are nearly indistinguishable from AF2 predictions of the structures adopted by the designed sequences (predictions of the full assemblies for the cyclic and dihedral designs, and trimeric substructures of the octahedral and icosahedral designs), and show little resemblance to proteins in the PDB (Fig. S6D). These include a number of oligomeric topologies not seen in nature, including two-layer beta strand barrels (Fig. 3A, C10 symmetry) and complex mixed alpha/beta topologies (Fig. 3A, C8 symmetry) (closest TM align in PDB: 6BRP, 0.47; 6BRO, 0.43 respectively).

**Figure 3:**
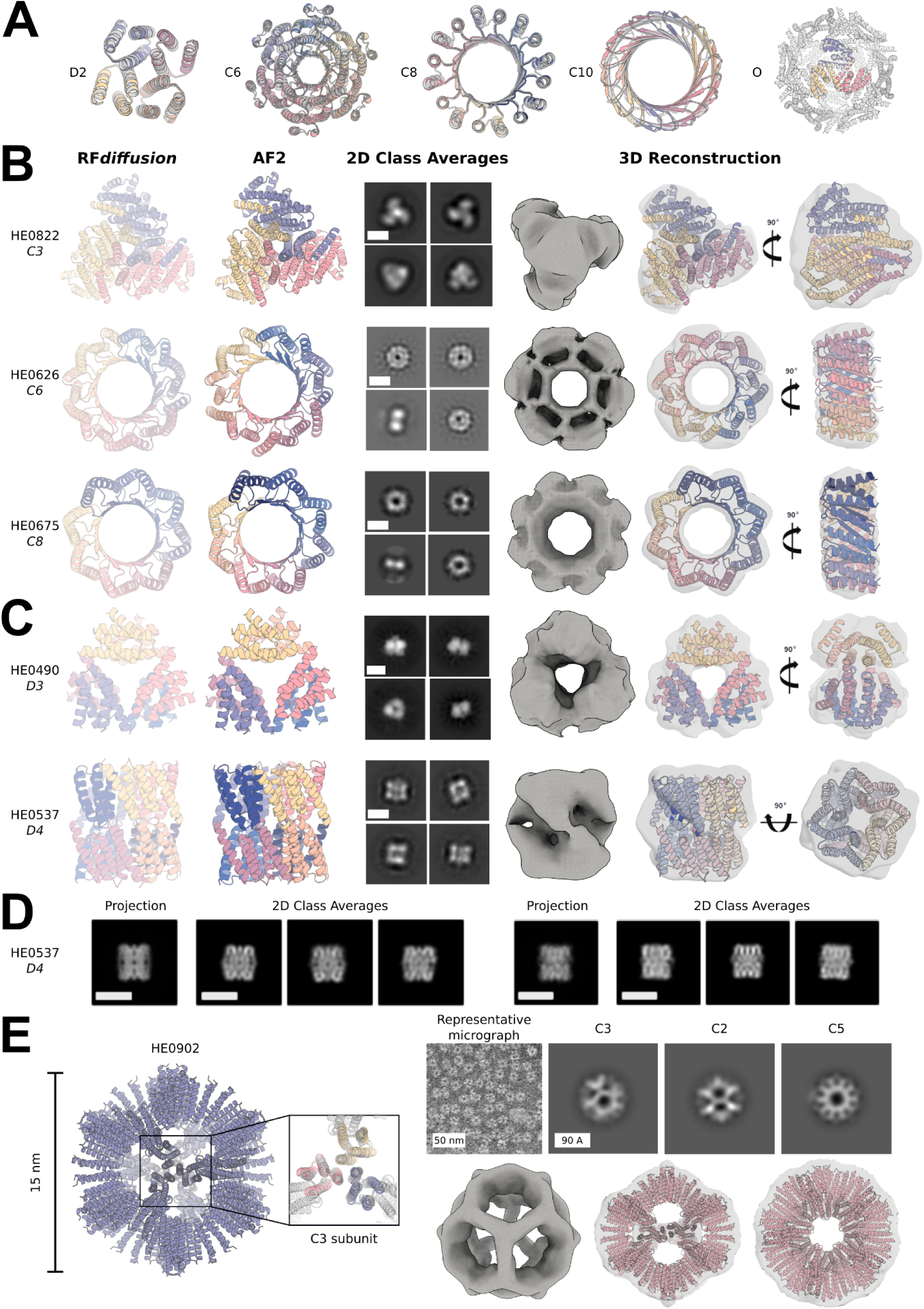
Design and experimental characterization of high-order symmetric oligomers. **A)** RF*diffusion*-generated assemblies overlaid with the AF2 structure predictions based on the designed sequences; in all 5 cases they are nearly indistinguishable. Symmetries are indicated to the left of the design models. The octahedral symmetries were validated by their C3 subunits only, as shown in panel A. **B-C)** Designed assemblies characterized by negative stain electron microscopy. Model symmetries: **B)** Cyclic: C3 (HE0822, 350 AA/chain); C6 (HE0626, 100 AA/ chain); C8 (HE0675, 60 AA/ chain) **C)** Dihedral: D3 (HE0490, 80 AA/ chain); and D4 (HE0537, 100 AA/ chain). From left to right: 1) symmetric design model, 2) AF2 prediction of design following sequence design with ProteinMPNN, 3) 2D class averages showing a combination of (at minimum) top and side views (scale bar = 60 Å for all class averages), 4) 3D reconstructions from class averages with the design model fit into the density map. The overall shapes are closely consistent with the design models, and confirm the intended oligomeric state. As in **A**), the AF2 predictions of each design are nearly indistinguishable from the original diffusion model (backbone RMSDs (Å) for HE0822, HE0626, HE0490, HE0675, and HE0537, are 1.33, 1.03, 0.60, 0.74, and 0.75, respectively). **D)** Two orthogonal side views of HE0537 by cryo-EM. Representative 2D class averages from the cryo-EM data are shown to the right of the predicted 2D projection images of the computational design model (lowpass filtered to 8 Å), which appear nearly identical to the experimental data. Scale bar shown is 60 Å for all images. **E)** Characterized icosahedral particle (HE0902, 100 AA/ chain) by negative stain electron microscopy. The design model, including the AF2 prediction of the C3 subunit are shown on the left. nsEM data are shown on the right: on top, a representative micrograph is shown alongside 2D class averages representing each axis of symmetry (C3, C2, and C5, from left to right) with their corresponding 3D reconstruction map views shown directly below and demonstrating high agreement to the design model.

We selected 608 designs for experimental characterization, and found using size exclusion chromatography (SEC) that at least 70 had oligomerization states closely consistent with the design models (within the 95% CI, 109 designs within the 99% CI, as determined by SEC calibration curves) (Fig. S11, S12). We collected negative stain electron microscopy (nsEM) data on a subset of these designs across different symmetry groups and, for most, distinct particles were evident with shapes resembling the design models in both the raw micrographs and subsequent 2D classifications (Fig. 3, and Fig. S6F). We describe these designs in the following paragraphs; most have structures that are, to our knowledge, unprecedented in nature.

As described above, RF*diffusion* was able to unconditionally generate a wide range of monomer structures, and while AF2 predictions and circular dichroism measurements were consistent with the design models, the structures were too small for electron microscopy-based structure validation. We took advantage of the increase in size upon oligomerization to evaluate, using electron microscopy, unconstrained structure generation for oligomers with subunits over 175 amino acids in length (Fig. 3B, top row). Electron microscopy characterization of a C3 design (HE0822) with 350 residue subunits (1050 residues in total) suggests that the actual structure is very close to the design, both over the 350 residue subunits and the overall C3 architecture. 2D class averages are clearly consistent with both top- and side-views of the design model, and a 3D reconstruction of the density had key features consistent with the design, including the distinctive pinwheel shape. Electron microscopy 2D class averages of C5 and C6 designs with greater than 750 residues (HE0795, HE0789, HE0841) were also consistent with the respective design models (Fig. S6F).

RF*diffusion* also generated cyclic oligomers with alpha/beta barrel structures that resemble expanded TIM barrels and provide an interesting comparison between innovation during natural evolution and innovation through deep learning. The TIM barrel fold, with 8 strands and 8 helices, is one of the most abundant folds in nature^35^. Electron microscopy characterization validated two RF*diffusion* cyclic oligomers which considerably extend beyond this fold (Fig. 3B, bottom rows). HE0626 is a C6 alpha/beta barrel composed of 18 strands and 18 helices, and HE0675 is a C8 octamer composed of an inner ring of 16 strands and an outer ring of 16 helices arranged locally in a very similar repeating pattern to the TIM barrel (1:1 helix:strand). By nsEM, we observed 2D class averages for HE0626 that resemble this two ring organization, and for both HE0626 and HE0675 we were able to obtain 3D reconstructions that are in agreement with the computational design models. The HE0600 design is also an alpha-beta barrel (Fig. S6F), but has two strands for every helix (24 strands and 12 helices in total) and is hence locally quite different from a TIM barrel. Whereas natural evolution has extensively explored structural variations of the classic 8-strand/8-helix TIM barrel fold, RF*diffusion* can more readily explore global changes in barrel curvature, enabling discovery of TIM barrel-like structures with many more helices and strands.

RF*diffusion* readily generated structures with dihedral and tetrahedral symmetries (Fig. 3C, Fig. S6E,F). SEC characterization indicated that 38 D2, 7 D3, and 3 D4 designs had the expected molecular weights (these have 4, 6, and 8 chains, respectively) (Fig. S12). While the D2 dihedrals are too small for nsEM, 2D class averages–and for some, 3D reconstructions– of D3 and D4 designs were congruent with the overall topologies of the design models (Fig. 3C, Fig. S6F). The reconstruction for the D3 HE0490 shows the characteristic triangular shape of the design. Similarly, the 3D reconstruction of the D4 HE0537 closely matches the design model, recapitulating the approximate 45° offset between tetramic subunits. We were also able to obtain cryogenic electron microscopy (cryo-EM) data for HE0537; however, a preferred orientation precluded generation of a reliable 3D reconstruction for in-depth structural analysis. Nonetheless, the resulting 2D class averages bear a striking level of secondary-structure similarity to generated 2D projections of the corresponding design model (Fig. 3D). 2D class averages for a 12 chain tetrahedron (HE0964) were consistent with the design, but we were unable to generate a 3D reconstruction of high confidence due to a lack of clear discernable design features visible at the resolution range provided by nsEM (Fig. S6F).

Icosahedra have 60 subunits arrayed around 2-fold, 3-fold and 5-fold symmetry axes. Of the 48 icosahedra selected for experimental validation, one was confirmed by nsEM to form the intended assembly. As shown in Fig. 3E on the left, HE0902 is a 15nm (diameter) highly-porous icosahedron composed of alpha helical subunits. The nsEM micrographs reveal highly homogeneous particles, and the corresponding 2D class averages and 3D reconstruction nearly perfectly match the design model (Fig. 3E), with triangular hubs arrayed around the empty C5 axes. Designs such as HE0902 (and future similar large assemblies) should be useful as new nanomaterials and vaccine scaffolds, with robust assembly and (in the case of HE0902) the outward facing N- and C-termini offering multiple possibilities for antigen display.

### Functional motif scaffolding

We next investigated the use of RF*diffusion* for scaffolding protein structural motifs that carry out binding and catalytic functions, where the role of the scaffold is to hold the motif in precisely the 3D geometry needed for optimal function. In RF*diffusion*, we input motifs as 3D coordinates (including sequence and sidechains) both during conditional training and inference, and RF*diffusion* builds scaffolds that hold the motif atomic coordinates in place. A number of deep learning methods have been developed recently to address this problem, including RF_joint_ Inpainting^4^, constrained Hallucination^4^, and other DDPMs^5,8,25^. To rigorously evaluate the performance of these methods in comparison to RF*diffusion* across a broad set of design challenges, we established an *in silico* benchmark test comprising 25 motif-scaffolding design problems addressed in six recent publications encompassing several design methodologies^4,5,25,36–38^. The benchmark includes 25 challenges that span a broad range of motifs, including simple “inpainting” problems, viral epitopes, receptor traps, small molecule binding sites, binding interfaces and enzyme active sites. Full details of this benchmark set are described in Supplementary Table 9.

RF*diffusion* solves all but two of the 25 benchmark problems, with greater success (23/25) than both Hallucination (15/25) and RF_joint_ Inpainting (12/25) (Fig. 4A-B). For 22/23 of the problems solved by RF*diffusion*, it also has a higher fraction of successful designs than either Hallucination or RF_joint_ Inpainting. The excellent performance of RF*diffusion* required no hyperparameter tuning or external potentials; this contrasts with Hallucination, for which problem-specific optimization can be required. In 17/23 of the problems, RF*diffusion* generated successful solutions with higher success rates when noise was not added during the reverse diffusion trajectories (see Fig. S2I for further discussion of the effect of noise on design quality).

**Figure 4:**
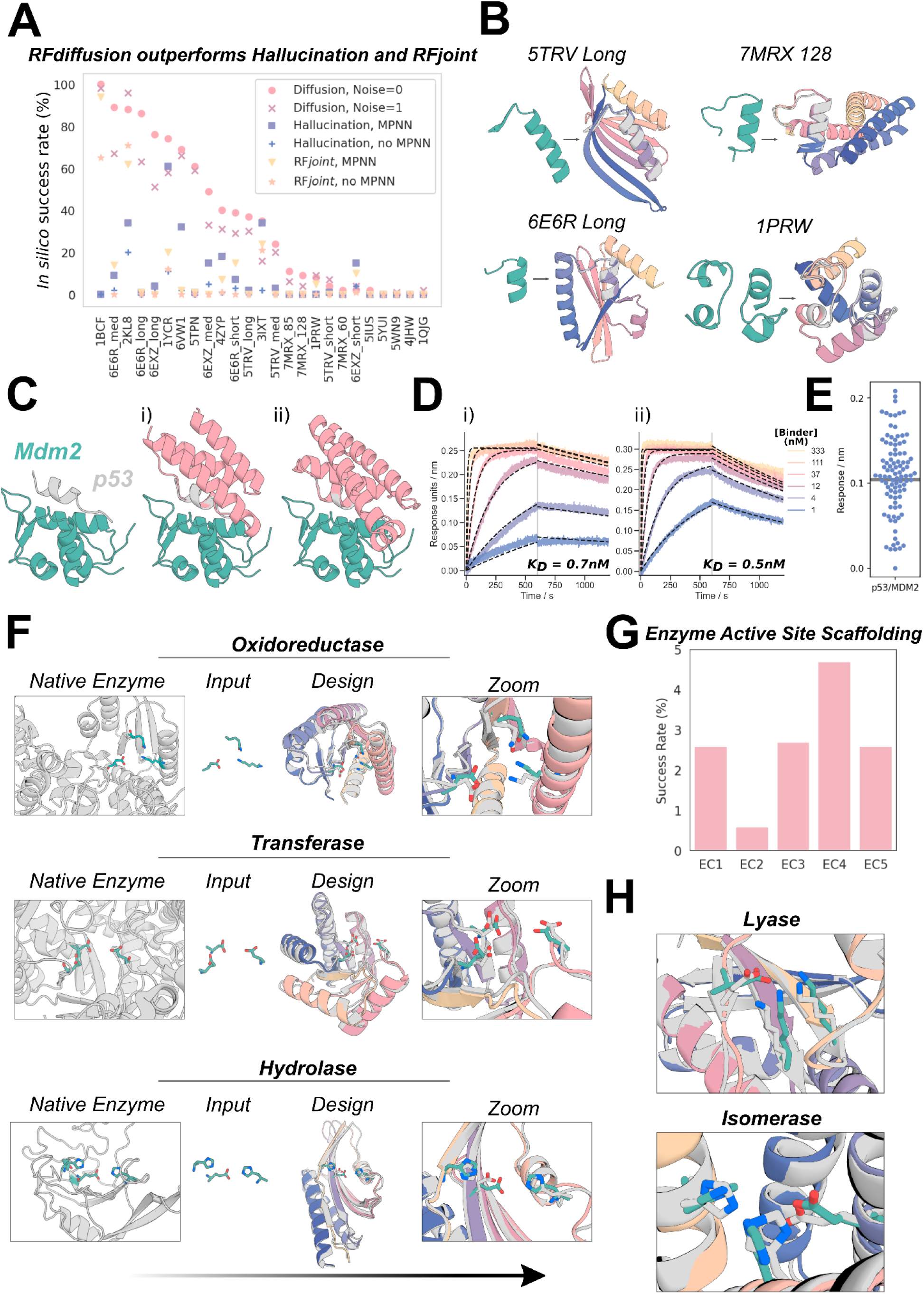
Scaffolding of diverse functional-sites with RF*diffusion*. **A)** RF*diffusion* has state of the art performance across 25 benchmark motif scaffolding problems collected from six recent publications, encompassing a broad range of motifs (Supplementary Table 9). Success was defined as AF2 RMSD to design model < 2Å, AF2 RMSD to the native functional site (the “motif”) < 1Å, and AF2 predicted alignment error (pAE) < 5, and the examples are ordered by success rate with RF*diffusion* (with noise scale = 0). 100 designs were generated per problem, with no prior optimization on the benchmark set (some optimization was necessary for the Hallucination results). Supplementary Table 10 presents full results. **B)** Four examples of designs for benchmarking problems where RF*diffusion* significantly outperforms existing methods. Teal: native motif; colors: AF2 prediction of an RF*diffusion* design. Metrics (RMSD AF2 vs design / vs native motif (Å), AF2 pAE): 5TRV Long: 1.17/0.57, 4.73; 6E6R Long: 0.89/0.27, 4.56; 7MRX Long: 0.84/0.82 4.32; 1PRW: 0.77/0.89, 4.49. **C)** RF*diffusion* can scaffold the native p53 helix that binds to MDM2 (left) and makes additional contacts with the target (right, average 31% increased surface area). **D)** Biolayer interferometry (BLI) measurements demonstrate high affinity (0.7nM and 0.5nM) binding to MDM2 for the two designs shown in **C**; the native p53 helix affinity is 600nM^**40**^. **E)** Experimental success rates were high, with 55/95 designs showing significant binding to MDM2 (> 50% of maximum response). **F)** After fine-tuning on a task that mimics active-site scaffolding (Methods 4.2), RF*diffusion* can scaffold a broad range of enzyme active sites. Three examples are shown (Enzyme Classes, EC, 1-3; ref [^**50**^]). Left to right: native enzyme (PDB: 1A4I, 1CWY, 1DE3); catalytic site (teal); RF*diffusion* output (gray: model, colors: AF2 prediction); zoom of active site. **G)** *In silico* success rates on active sites derived from EC1-5 (AF2 Motif RMSD vs native: backbone < 1Å, backbone and sidechain atoms < 1.5Å, RMSD AF2 vs design < 2, AF2 pAE < 5). H) Zoom in views of two further successful designs, for EC4 and EC5 (active sites from PDB: 1P1X, 1SNZ). Metrics for examples in (**F**) and (**H**) (AF2 vs design backbone RMSD, AF2 vs design motif backbone RMSD, AF2 vs design motif full-atom RMSD, AF2 pAE): EC2: 0.93Å, 0.50Å, 1.29Å, 3.51; EC3: 0.92Å, 0.60Å, 1.07Å, 4.59; EC4: 0.93Å, 0.80Å, 1.03Å, 4.41; EC5: 0.78Å, 0.44Å, 1.14Å, 3.32.

One of the benchmark problems is the scaffolding of the p53 helix that binds MDM2. Inhibiting this interaction through high-affinity competitive inhibition by scaffolding the p53 helix and making additional interactions with MDM2 is a promising avenue for therapeutics^39^. *In silico* success has been described elsewhere^4^, but experimental success has not been reported. We tested 96 designs scaffolding this helix, which were predicted to make additional interactions with MDM2, and identified 0.5nM and 0.7nM binders (Fig. 4C-D), three orders of magnitude higher affinity than the reported 600nM affinity of the p53 peptide alone^40^. The success rate for this problem was particularly striking with 55/95 designs showing some detectable binding at 10μM (Fig. 4E) and multiple designs with affinities in the low-to sub-nanomolar range (Fig. 4D).

### Scaffolding enzyme active sites

A grand challenge in protein design is to scaffold minimal descriptions of enzyme active sites comprising a few single amino acids. While some *in silico* success has been reported previously^4^, a general solution that can readily produce high-quality, orthogonally-validated outputs remains elusive. Following fine-tuning on a task mimicking this problem (Methods 4.2), RF*diffusion* was able to scaffold enzyme active sites comprising multiple sidechain and backbone functional groups with high accuracy and *in silico* success rates across a range of enzyme classes (Fig. 4F-H). While RF*diffusion* is currently unable to *explicitly* model bound small molecules (see conclusion), the substrate can be *implicitly* modeled using an external potential to guide the generation of “pockets” around the active site. As a demonstration, we scaffold a retroaldolase active site triad while implicitly modeling its substrate (Fig. S7).

### Symmetric functional-motif scaffolding for metal coordinating assemblies and antiviral therapeutics and vaccines

A number of important design challenges involve the scaffolding of multiple copies of a functional motif in symmetric arrangements. For example, many viral glycoproteins are trimeric, and symmetry matched arrangements of inhibitory domains can be extremely potent^41–44^. Conversely, symmetric presentation of viral epitopes in an arrangement that mimics the virus could induce new classes of neutralizing antibodies^45,46^. To explore this general direction, we sought to design trimeric multivalent binders to the SARS-CoV-2 spike protein. In previous work, flexible linkage of a binder to the ACE2 binding site (on the spike protein receptor binding domain) to a trimerization domain yielded a high-affinity inhibitor that had potent and broadly neutralizing antiviral activity in animal models^41^. Ideally, however, symmetric fusions to binders would be rigid, so as to reduce the entropic cost of binding while maintaining the avidity benefits from multivalency. We used RF*diffusion* to design C3 symmetric trimers which rigidly hold three binding domains (the functional motif in this case) such that they exactly match the ACE2 binding sites on the SARS-CoV-2 spike protein trimer. Design models were confidently predicted by AF2 to both assemble as C3-symmetric oligomers, and to scaffold the AHB2 SARS-CoV-2 binder interface with high accuracy (Fig. 5A).

**Figure 5:**
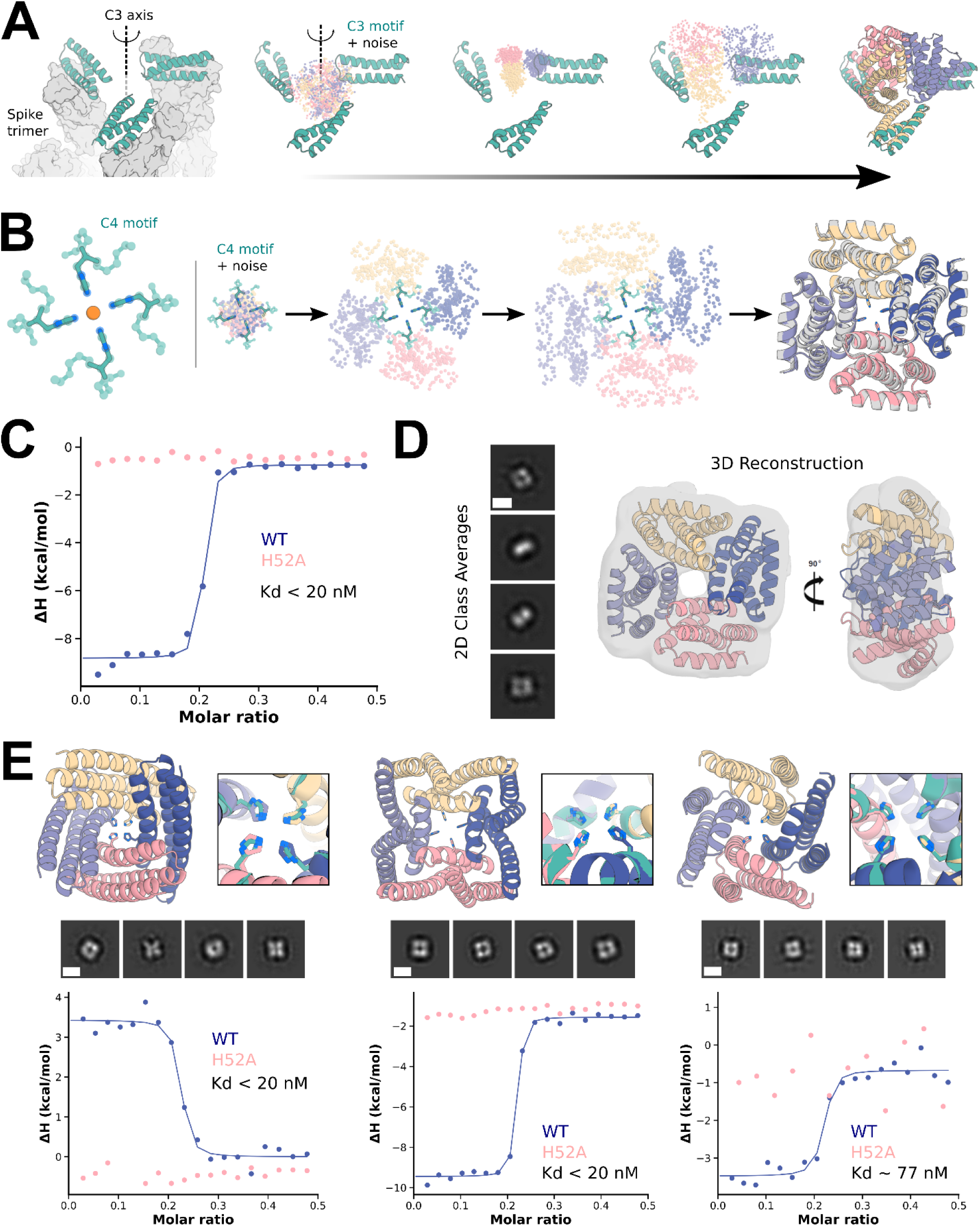
Symmetric motif scaffolding with RF*diffusion*. **A)** Design of C3-symmetric oligomers to scaffold the binding interface of the designed ACE2 mimic, AHB2 (left, teal), against the SARS-CoV-2 spike trimer (left, gray). Starting from AHB2 bound to each of the three ACE2 binding sites on the spike trimer, RF*diffusion* was used to generate C3-symmetric oligomers that hold the three AHB2 exactly in place to simultaneously engage the binding sites on all three spike subunits.The first 55 amino-acids of each minibinder copy are used as the symmetric motif input to RF*diffusion* (middle). The method produces designs whose AF2 predictions (right) recapitulate the mini-binder motif with high accuracy on the asymmetric unit (0.6 Å RMSD) and good accuracy the symmetric motif (2.9 Å RMSD). **B)** Design of C4-symmetric oligomers to scaffold a theoretical Ni^2+^ binding motif (left). Starting from a square-planar set of histidine rotamers within three-residue helical fragments (Methods 5.9) and C4-symmetric noise, an RF*diffusion* trajectory iteratively builds a symmetric oligomer scaffolding the theoretical Ni^2+^ binding domain (middle). AF2 predictions (color) overlaid with the RF*diffusion* design model (gray) agree closely, with backbone RMSD for the particular example < 1.0 Å (right). **C)** Isothermal titration calorimetry (ITC) binding isotherm of design (“E1”) and corresponding H52? mutant. The inflection point of the wild-type isotherm (blue) displays an estimated dissociation constant of less than 20 nM at the designed metal:monomer stoichiometry of 1:4. Importantly, the H52A mutant isotherm (pink) displays complete ablation of binding, indicating the scaffolded histidine at position 52 of each protomer is critical for metal binding. **D)** 2D class averages (left) and corresponding 3D reconstruction with the model of design E1 docked into the 3D reconstructed density (right). The four-fold symmetry and general shape of the designed oligomer can be readily identified in the 2D class averages, with both top-down views and side views captured (scale bar = 60 Å). **E)** Additional experimentally characterized Ni^2+^ binding oligomers G3 (left), C10 (middle), and A5 (right) from RF*diffusion* show structural diversity in successful designs. Design models and binding-site zoom (top, AF2 in colors and ideal motif in teal) show close recapitulation of the motif sidechains by AF2. 2D nsEM class averages (middle, scale bar = 60 Å), and binding isotherms for wild-type and H52A mutant (bottom) indicate tight Ni^2+^ binding mediated directly by the scaffolded histidines at the designed 1:4 stoichiometry.

The ability to scaffold functional sites with any desired symmetry opens up new approaches to designing metal-coordinating protein assemblies. Divalent transition metal ions exhibit distinct preferences for specific coordination geometries (e.g., square planar, tetrahedral, and octahedral) with ion-specific optimal sidechain–metal bond lengths. RF*diffusion* provides a general route to building up symmetric protein assemblies around such sites, with the symmetry of the assembly matching the symmetry of the coordination geometry. As a first test, we sought to design square planar Ni^2+^ binding sites. We designed C4 protein assemblies with four central histidine imidazoles arranged in an ideal Ni^2+^-binding site with square planar coordination geometry. Diverse designs starting from various different C4-symmetric histidine square planar sites (Fig. 5B, Fig. S8A,B,C) had good *in silico* success (Fig. S8D), with the histidine residues in near ideal geometries for coordinating metal in the AF2 predicted structures (Fig. 5E Fig. S8B,C,E,F).

We expressed and purified 44 designs in *E. coli*., and found that 37 had SEC chromatograms consistent with the intended oligomeric state (Fig. S9B). 36 of these designs were tested for Ni^2+^ coordination by isothermal titration calorimetry. 18 designs bound Ni^2+^ with dissociation constants ranging from low nanomolar to low micromolar (Fig. 5C,E and Fig. S9A). The inflection points in the wild-type isotherms indicate binding with the designed stoichiometry, a 1:4 ratio of ion:monomer. While most of the designed proteins displayed exothermic metal coordination, in a few cases binding was endothermic (Fig. 5E, left, Fig. S9A), suggesting that Ni^2+^ coordination is entropically driven in these assemblies. To confirm that Ni^2+^ binding was indeed mediated by the scaffolded histidine 52, we mutated this residue to alanine, which abolished or dramatically reduced binding in all cases (Fig. S9A,C and Fig. 5C,E). We structurally characterized by nsEM a subset of the designs – E1, C10, G3, and A5 – that displayed histidine-dependent binding. All four designs exhibited clear 4-fold symmetry both in the raw micrographs and in 2D class averages (Fig. 5D,E), with design E1 also clearly displaying 2-fold axis “side-views” with a measured diameter approximating the design model. A 3D reconstruction of E1 was in close agreement to the design model (Fig. 5D).

### Design of de novo protein-binding proteins

The design of high-affinity binders to target proteins is a grand challenge in protein design, with numerous therapeutic applications^47^. A general method to de novo design binders to protein binders from target structure information alone using the physically-based Rosetta method was recently described^12^. Subsequently, utilizing ProteinMPNN for sequence design and AF2 for design filtering was found to improve design success rates^26^. However, experimental success rates were low, requiring many thousands of designs to be screened for each design campaign^12^, and the approach relied on pre-specifying a particular set of protein scaffolds as the basis for the designs, inherently limiting the diversity and shape complementarity of possible solutions^12^. To our knowledge, no deep-learning method has yet demonstrated experimental general success in designing completely *de novo* binders.

We reasoned that RF*diffusion* might be able to address this challenge by directly generating binding proteins in the context of the target. For many therapeutic applications, for example blocking a protein-protein interaction, it is desirable to bind to a particular site on a target protein. To enable this, we fine-tuned RF*diffusion* on protein complex structures, providing as input a subset of the residues on the target chain (called “interface hotspots”) to which the diffused chain binds (Fig. 6A, Fig. S10A,B). To enable control over binder scaffold topology, we fine-tuned an additional model to condition binder diffusion on secondary structure and block-adjacency information, in addition to conditioning on interface hotspots (Fig. S10C-D, Methods 4.3).

**Figure 6:**
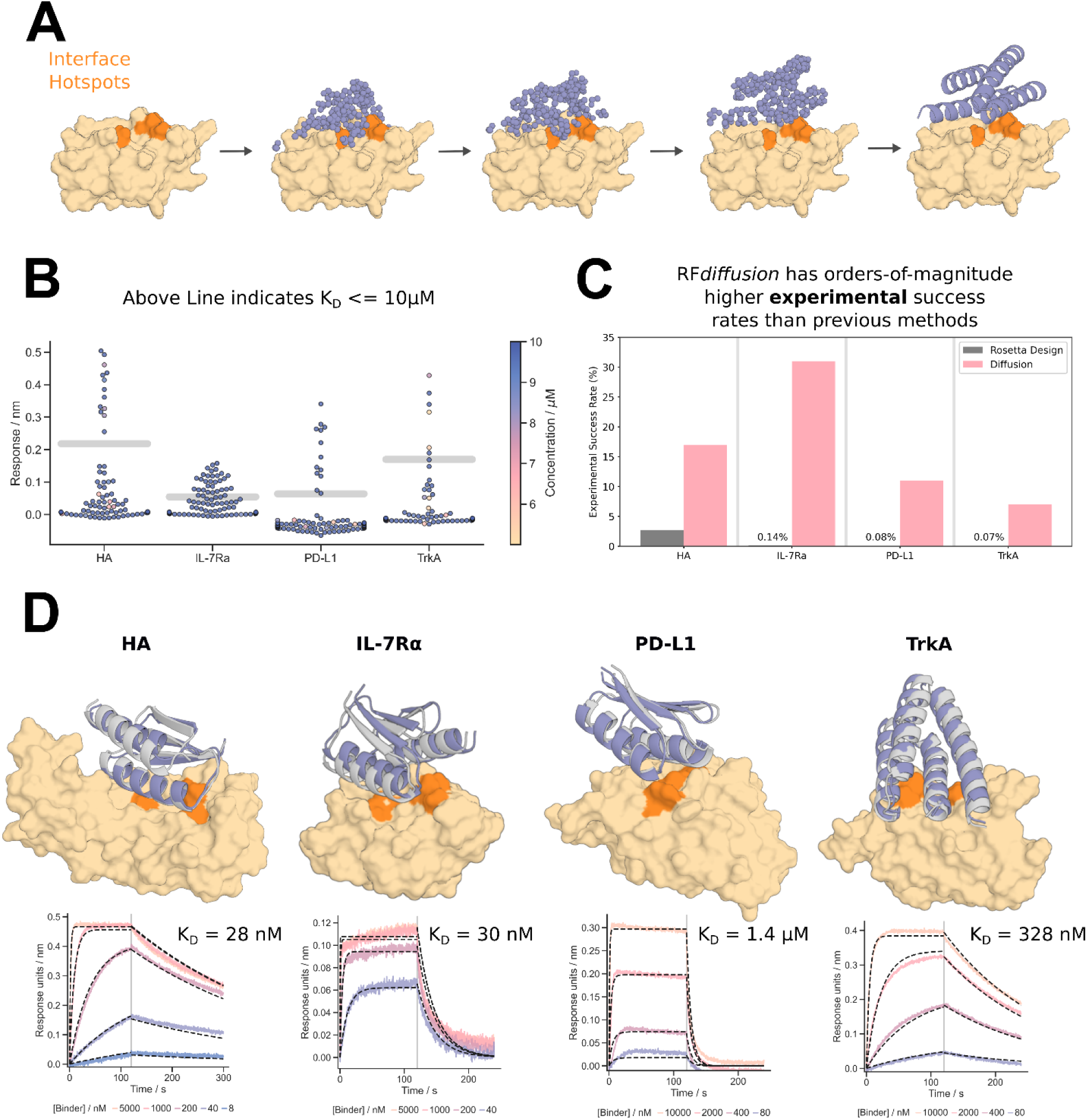
Design of *de novo* protein-binding proteins. **A-B)** *De novo* binders were designed to four protein targets; Influenza Hemagglutinin A, IL-7 Receptor-a, PD-L1, and TrkA receptor. RF*diffusion* generates protein binders by conditioning on interface hotspot residues. Additionally, the general topology of the binders generated by RF*diffusion* can be controlled using fold-conditioning. **B)** *De novo* protein binders were identified for all four of the targets for which we could obtain suitable target protein (see Fig. S15 for expression data on Insulin receptor binders). Designs that bound at 10μM during single point BLI screening with a response equal to or greater than 50% of the positive control were considered binders. Concentration is denoted by hue for designs that were screened at concentrations less than 10μM and thus may be false negatives. **C)** RF*diffusion* designed binders have very high experimental success rates compared to the previous design campaigns against the same targets. For IL-7Ra, PD-L1, and TrkA, RF*diffusion* has success rates ∼2 orders-of-magnitude higher than the original design campaigns. **D)** For each target, the highest affinity binder is shown alongside a BLI titration series. Reported K_D_s are based on global kinetic fitting with fixed global R_max_. Yellow/orange: target/hotspot residues; gray: design model; purple: AF2 prediction (RMSD AF2 vs design, left to right: 0.7Å, 0.9Å, 1.2Å, 0.7Å).

To compare RF*diffusion* to previous binder design methods, we performed binder design campaigns against 4 targets: Influenza A H1 Hemagglutinin (HA) ^48^ (HA), Interleukin-7 Receptor-a (IL-7Ra)^12^, Programmed Death-Ligand 1 (PD-L1)^12^, and Tropomyosin Receptor Kinase A (TrkA) ^12^ (we also designed against the insulin receptor, and although these binder designs expressed solubly and monomerically (Fig. S15), we were not able to obtain suitable target receptor for experimental testing). We designed putative binders to each target, both with and without conditioning on compatible fold information, with high success rates (Fig. S10E,F). Designs were filtered by AF2 confidence in the interface and monomer structure^26^, and 95 were selected for each target for experimental characterization.

The designed binders were expressed in *E. coli* and purified, and binding was assessed through single point biolayer interferometry (BLI) screening at 10μM binder (Fig. 6B). In each case a positive control was included that binds to the site targeted by the designs on the target protein^12^. The overall success rate, as defined by binding at 10μM at or above 50% of the maximal response for the positive control, was 18% (this is a conservative estimate as some designs which showed binding had insufficient material to permit screening at 10μM (Fig. 6B)). This is a success rate increase of approximately 2 orders-of-magnitude over our previous Rosetta-based method on the same targets (Fig. 6C). Binders were identified for all 4 targets, with fewer than 100 designs tested per target compared to thousands in previous studies. Full BLI titrations for a subset of the designs showed moderate to high affinities with no further experimental optimization, including HA and IL-7Ra binders with affinities of approximately 30nM (Fig. 6D). To assess binder specificity, 6 of the highest affinity IL-7Ra binders were assessed via competition BLI, and all 6 competed for binding with the structurally validated positive control (Fig. S14).

## Discussion

RF*diffusion* is a major improvement over current physically-based and deep learning protein design methods over a wide range of design challenges. Substantial progress was recently made using Rosetta in designing binding proteins from target structural information alone^12^, but this required testing tens of thousands of designs. RF*diffusion* achieves experimental success rates that are two orders of magnitude higher. Consequently, high affinity binders (at least to the targets experimentally characterized here) can be identified through testing only dozens of designs. In the accompanying paper (Vázquez Torres *et al*.), we demonstrate the ability of RF*diffusion* to design picomolar affinity binders to flexible helical peptides, further highlighting the utility of RF*diffusion* for *de novo* binder design. Vázquez Torres *et al*. also show that RF*diffusion* can be used to improve upon starting designs by partial noising and denoising, which enables tunable sampling around a given input structure. For peptide binder design, this enabled increases in affinity of nearly three orders of magnitude, without high-throughput screening of designs.

There has been recent progress in scaffolding protein functional motifs using deep learning methods (Hallucination, RF_joint_ Inpainting, and diffusion), but Hallucination becomes very slow for large systems, inpainting fails when insufficient starting information is provided, and previous diffusion methods had quite low accuracy. Our benchmark tests show that RF*diffusion* considerably outperforms all previous methods in the complexity of the motifs that can be scaffolded, the ability to precisely position sidechains (for catalysis and other functions), and the accuracy of motif recapitulation by AF2. The robust design of MDM2 binding proteins with three orders of magnitude higher binding affinities than the scaffolded P53 motif experimentally demonstrates the power of RF*diffusion* for motif scaffolding.

For the classic unconstrained protein structure generation problem, RF*diffusion* readily generates novel protein structures with as many as 600 residues that are accurately predicted by AF2 (and ESMFold), far exceeding the complexity and accuracy achieved by previously described diffusion and other methods. Experimental data demonstrate that designs express solubly, with CD spectra consistent with the design models. That the designs are also extremely thermostable also shows that RF*diffusion* designs retain the desirable ideality and stability of previous *de novo* design methods, while achieving considerably increased complexity. The versatility and control provided by diffusion models enabled extension of RF*diffusion* unconditional generation to higher order architectures with any desired symmetry (Hallucination methods are primarily limited to cyclic symmetries); experimental characterization of a subset of these designs using electron microscopy revealed structures very similar to the design models and largely without precedent in nature. Combining the accurate motif scaffolding with the ability to design symmetric assemblies, we were able to scaffold functional motifs spanning multiple symmetrically arranged chains.

Overall, the complexity of the problems solvable with RF*diffusion* and the robustness and accuracy of the solutions (extensively validated both *in silico* and experimentally) far exceeds what has been achieved previously. In a manner reminiscent of the generation of images from text prompts, RF*diffusion* makes possible, with minimal specialist knowledge, the generation of proteins from very simple molecular specifications (for example, from a specification of a target protein, high affinity binders to that protein, and from specification of a desired symmetry, diverse protein assemblies with that symmetry).

The power and scope of RF*diffusion* can be extended in several directions. RF has recently been extended to nucleic acids and protein-nucleic acid complexes^49^, which should enable RF*diffusion* to design nucleic acid binding proteins, and perhaps folded RNA structures. Extension of RF to incorporate ligands should similarly enable extension of RF*diffusion* to explicitly model ligand atoms, allowing the design of protein-ligand interactions. The ability to customize RF*diffusion* to specific design challenges by addition of external potentials and by fine-tuning (as illustrated here for catalytic site scaffolding, binder-targeting and fold-specification), along with continued improvements to the underlying methodology, should enable protein design to achieve still higher levels of complexity, to approach and – in some cases – surpass what natural evolution has achieved.

## Acknowledgements

We thank Namrata Anand and Doug Tischer for helpful discussions, and Indrek Kalvet and Yakov Kipnis for providing helpful Rosetta scripts. We thank Rachel Wu, Jody Mou, Kristy Choi, Luhuan Wu, and David Blei for valuable feedback during writing. We thank Ian Haydon for help with graphics. We also thank Luki Goldschmidt and Kandise VanWormer, respectively, for maintaining the computational and wet lab resources at the Institute for Protein Design.

This work was supported by gifts from Microsoft (D.J., M.B., D.B.), Amgen (J.L.W.), the Audacious Project at the Institute for Protein Design (B.L.T., I.S., J.Y., H.E., D.B.), the Washington State General Operating Fund supporting the Institute for Protein Design (P.V., I.S.), grant INV-010680 from the Bill and Melinda Gates Foundation Grant (W.B.A., D.J., J.W., D.B.), grant DE-SC0018940 MOD03 from the U.S. Department of Energy Office of Science (A.J.B., D.B.), grant 5U19AG065156-02 from the National Institute for Aging (S.V.T., D.B.), an EMBO long-term fellowship ALTF 139-2018 (B.I.M.W.), the Open Philanthropy Project Improving Protein Design Fund (R.J.R., D.B.), The Donald and Jo Anne Petersen Endowment for Accelerating Advancements in Alzheimer’s Disease Research (N.R.B.), a Washington Research Foundation Fellowship (S.J.P.), a Human Frontier Science Program Cross Disciplinary Fellowship (LT000395/2020-C, L.F.M.), an EMBO Non-Stipendiary Fellowship (ALTF 1047-2019, L.F.M.), the Defense Threat Reduction Agency grants HDTRA1-19-1-0003 (N.H., D.B.) and HDTRA12210012 (F.D.), the Institute for Protein Design Breakthrough Fund (A.C., D.B.), an EMBO Postdoctoral Fellowship (ALTF 292-2022, J.L.W.) and the Howard Hughes Medical Institute (A.C., W.S., R.R., D.B.), an NFS-GRFP (J.Y), an NSF Expeditions grant (1918839, J.Y, R.B., T.S.J.), the Machine Learning for Pharmaceutical Discovery and Synthesis consortium (J.Y, R.B., T.S.J.), the Abdul Latif Jameel Clinic for Machine Learning in Health (J.Y, R.B., T.S.J), the DTRA Discovery of Medical Countermeasures Against New and Emerging threats program (J.Y, R.B., T.S.J), and EPSRC Prosperity Partnership EP/T005386/1 (E.M.), the DARPA Accelerated Molecular Discovery program and the Sanofi Computational Antibody Design grant (J.Y, R.B., T.S.J.). We thank Microsoft and AWS for generous gifts of cloud computing resources.

## Author Contributions

Conceived the study: J.L.W., D.J., N.R.B, B.L.T., J.Y., D.B.; Trained RF*diffusion*: J.L.W., D.J., N.R.B, W.A., B.L.T., J.Y.; Extended diffusion to residue orientations: B.L.T., J.Y. with assistance from V.D.B., E.M.; Generated experimentally characterized designs: H.E.E., D.J., J.L.W., N.R.B., N.H., W.S., P.V., I.S.; Generated computational designs: W.A., B.L.T., J.Y., D.J., J.L.W., N.R.B.; Experimentally characterized designs: H.E.E., A.J.B., R.J.R., L.F.M., B.I.M.W., S.J.P., N.H., A.C., S.V.T., J.L.W., B.L.T.; Contributed additional code: J.W., A.L., W.S.; Trained RF: M.B., F.D.; Offered supervision throughout the project: D.B., T.S.J. and R.B.; Wrote the manuscript: J.L.W., D.J., B.L.T., J.Y., N.R.B., D.B. All authors read and contributed to the manuscript. J.L.W. and D.J. agree that the order of their respective names may be changed to best suit their own interests for personal pursuits.

## Data Availability

All datasets presented in this work will be available upon publication, including design structures, AlphaFold2 models and experimental measurements.

## Code Availability

Code for running RFdiffusion will be released upon publication.

## Supplementary Figures

**Figure S1:**
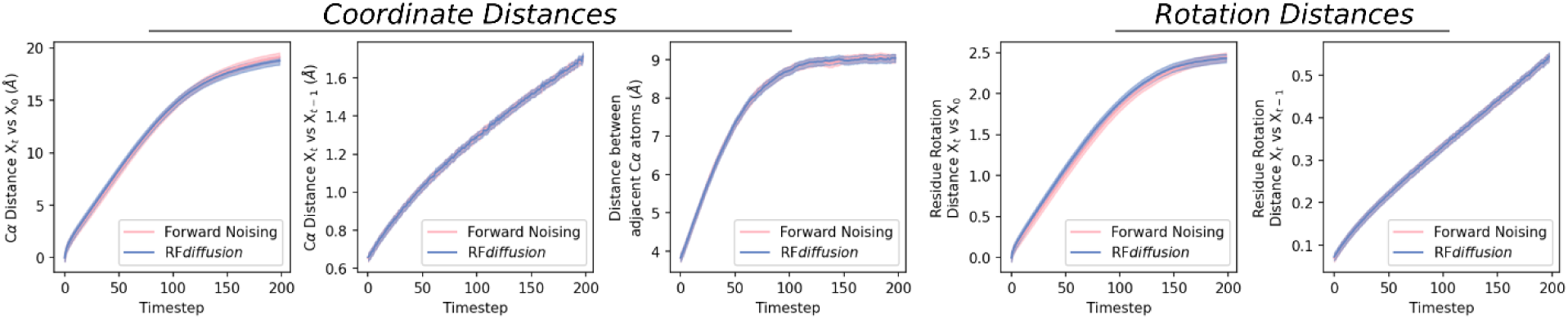
RF*diffusion* learns the distribution of the denoising process. Analysis of simulated forward (noising) and reverse (denoising) trajectories shows that the distribution of C_ɑ_ coordinates and residue orientations closely match, demonstrating that RF*diffusion* has learned the distribution of the denoising process as desired. Left to right: i) average distance between a C_ɑ_ coordinate at X_t_ and its position in X_0_; ii) average distance between a C_ɑ_ coordinate at X_t_ and X_t-1_; iii) average distance between adjacent C_ɑ_ coordinates at X_t_; iv) average rotation distance between a residue orientation at X_t_ and X_0_; v) average rotation distance between a residue orientation at X_t_ and X_t-1_.

**Figure S2:**
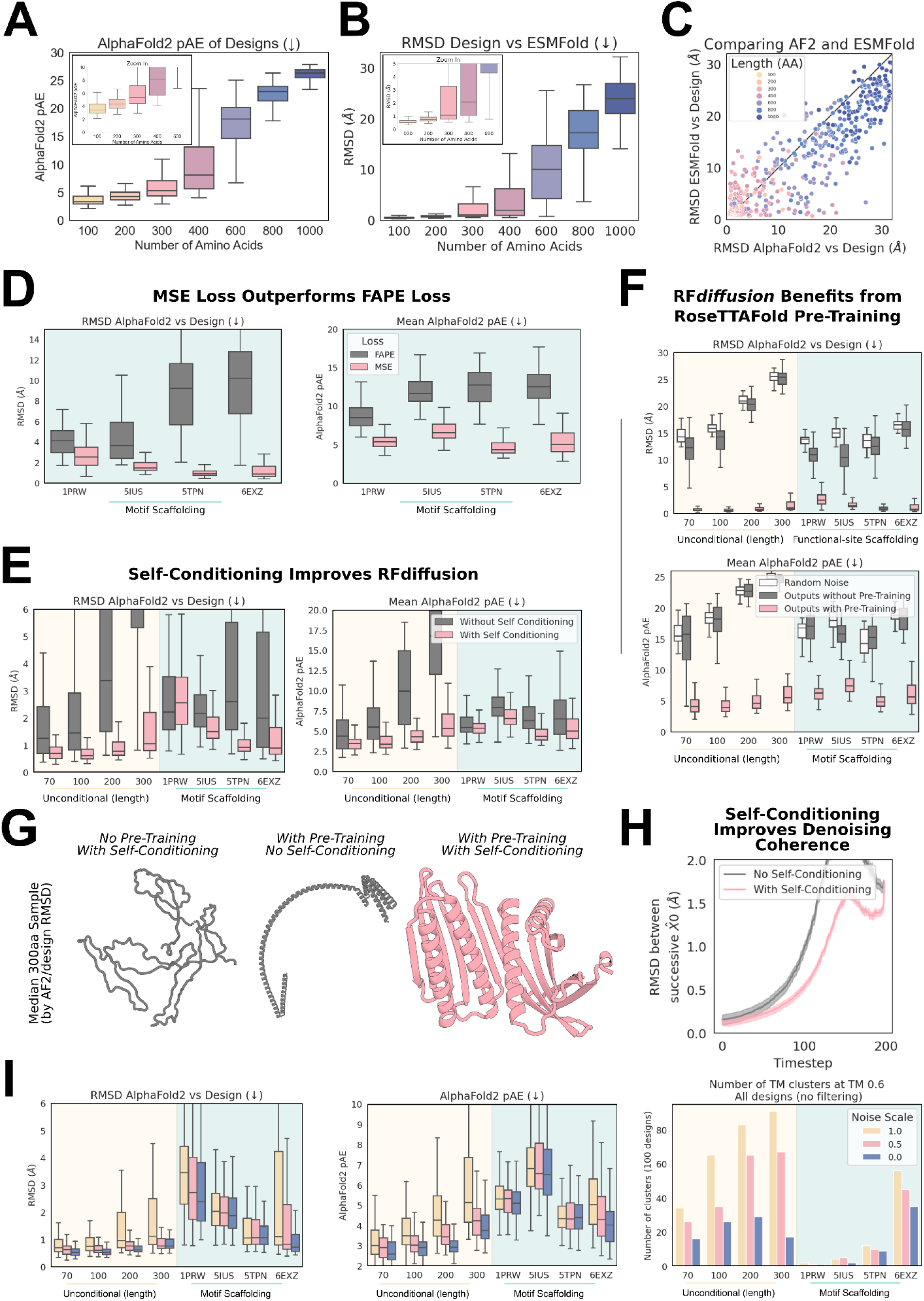
Training ablations reveal determinants of RF*diffusion* success. **A-C)** RF*diffusion* can generate high quality large unconditional monomers. Designs are routinely accurately recapitulated by AF2 (see also Fig. 2D), with high confidence (**A**) for proteins up to approximately 400 amino acids in length. **B)** Further orthogonal validation of designs by ESMFold. **C)** Recapitulation of the design structure is often better with ESMFold compared with AF2. For each backbone, the best of 8 ProteinMPNN sequences is plotted, with points therefore paired by backbone rather than sequence. **D)** Comparing RF*diffusion* trained with MSE loss on C_ɑ_ atoms and N-C_ɑ_-C backbone frames (Methods 2.5), rather than with FAPE loss^8,21^. The two models were benchmarked on motif scaffolding problems (see Methods 5.2 for justification of this decision), and across all cases, AF2 recapitulation of the structure (left) and AF2 confidence (right) was improved when RF*diffusion* was trained with MSE loss. Two-way ANOVA: Success rate *p<0*.*001*. **E)** Allowing the model to condition on its X_0_ prediction at the previous timestep (see Methods 2.4) improves designs. Designs with self-conditioning (pink) have improved recapitulation by AF2 (left) and better AF2 confidence in the prediction (right). Two-way ANOVA, *in silico* success rate: *p<0*.*001*. **F)** RF*diffusion* leverages the protein representations learned during RF pre-training. RF*diffusion* fine-tuned from pre-trained RF (pink) comprehensively outperforms a model trained for an equivalent amount of time, from untrained weights (gray). Training RF*diffusion* without pre-training (for 5 epochs) showed no significant improvement (in terms of *in silico* success rates) compared with generating ProteinMPNN sequences from random Gaussian-sampled coordinates (white, two-way ANOVA & Tukey’s test, *p<0*.*001; Random noise vs no pre-training, p=0*.*9 (n*.*s*.*); Random noise vs with pre-training, p<0*.*001; Pre-training vs not, p<0*.*001*). Note that the data in pink in **D-F** is the same data, reproduced in each plot for clarity. **G)** The median (by AF2 RMSD vs design) 300 amino acid unconditional sample highlighting the importance of self-conditioning and pre-training. Without pre-training, RF*diffusion* outputs bear little resemblance to proteins (gray, left). Without self-conditioning, outputs show characteristic protein secondary structures, but lack core-packing and ideality (gray, middle). With pre-training and self-conditioning, proteins are diverse and well-packed (pink, right). **H)** Greater coherence during unconditional denoising may partly explain the effect of self-conditioning. Successive X_0_ predictions are more similar when the model can self-condition (lower RMSD between X_0_ predictions, pink curve). Data are aggregated from unconditional design trajectories of 100, 200 and 300 residues. **I)** During the reverse (generation) process, the noise added at each step can be scaled (reduced). Reducing the noise scale improves the *in silico* design success rates (left, middle; two-way ANOVA & Tukey’s test: *p<0*.*001*, 0 vs 0.5: *p=0*.*13*, 0 vs 1: *p<0*.*001*; 0.5 vs 1: *p<0*.*001*). This comes at the expense of diversity, with the number of unique clusters at a TM score cutoff of 0.6 reduced when noise is reduced (right). Note throughout this figure the 6EXZ_long benchmarking problem is abbreviated to 6EXZ for brevity.

**Figure S3:**
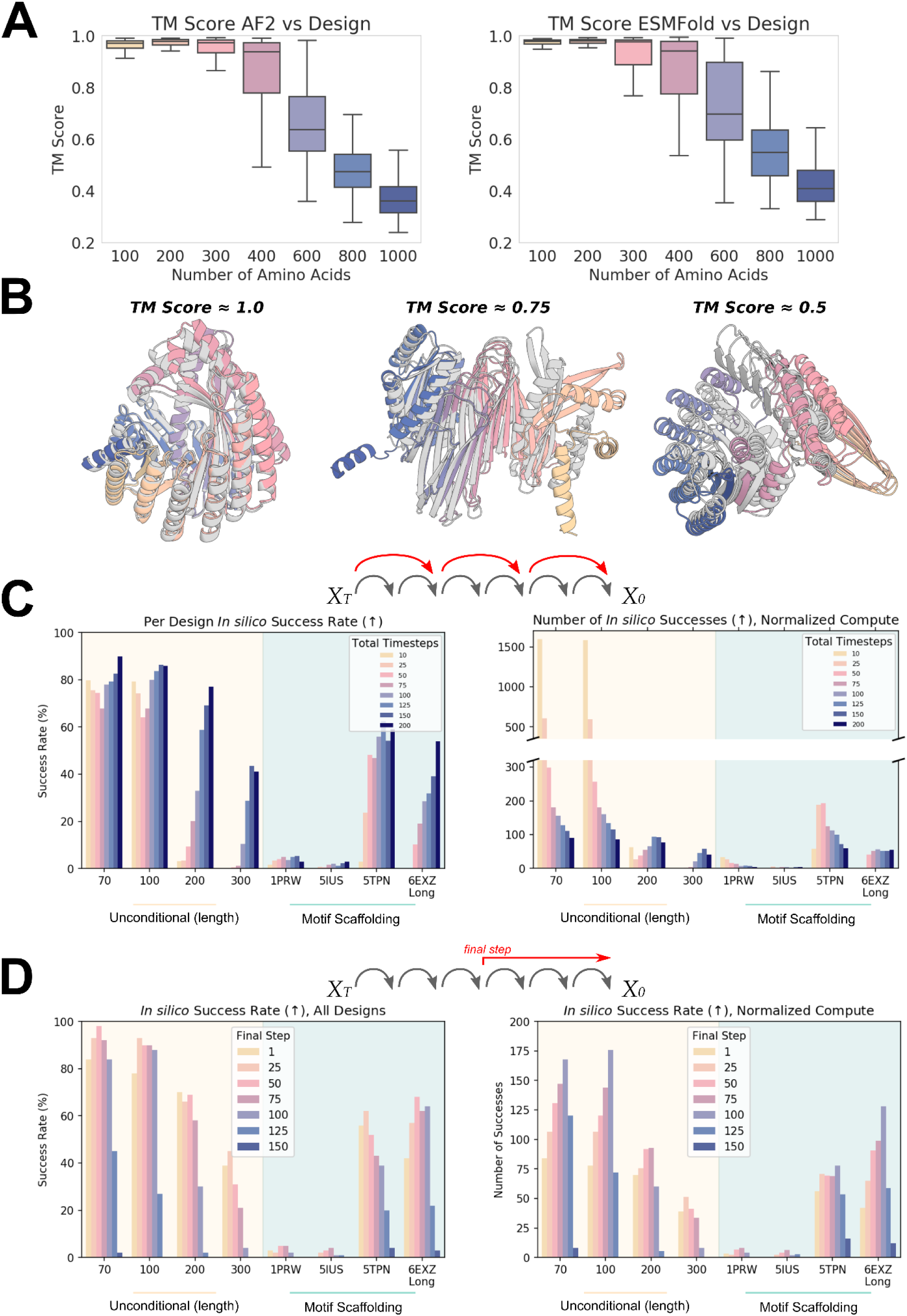
Optimizing inference and improving metrics for *in silico* success. **A-B)** TM score between a design and a subsequent orthogonal prediction (e.g. AF2), has been previously used, typically with a threshold of > 0.5, as a metric for design success. **A)** RF*diffusion* designs have high TM score agreement to both the AF2 (left) and ESMFold (right) predictions of the unconditional structures, with TM > 0.5 for a significant fraction of designs even up to 1000 amino acids in length. **B)** TM score is, however, much less stringent than RMSD alignment. Depicted here are three unconditional RF*diffusion* designs of 600 amino acids in length (gray), overlaid with the AF2 prediction (colors), with TM scores of 0.983, 0.757 and 0.506 respectively. While a TM score of 0.5 clearly shows some resemblance to the designed structure, it differs significantly and should not be classed as “successfully designed”. RMSD with a strict threshold (for example, 2Å) is significantly more stringent. RMSDs for the displayed designs are 1.15Å, 9.78Å and 21.4Å respectively. **C-D)** While RF*diffusion* is trained to generate samples over 200 timesteps, in many cases, trajectories can be shortened to improve computational efficiency. **C)** Bigger steps can be taken between timesteps at inference. While decreasing the number of timesteps typically reduces the per-design success rate (left), when normalized for compute budget (right), it is often more efficient to run more trajectories with fewer timesteps. For example, while generating 100 amino acid unconditional proteins, using a schedule with just 10 timesteps (as opposed to 200) allows the generation of 1584 *in silico* successful designs in the time taken to generate 86 successful designs with 200 timesteps. As problems get more challenging, however, this no longer remains the case (for example, fourth column, with generation of 300 amino acid designs). **D)** An alternative to taking larger steps is to stop trajectories early (possible because RF*diffusion* predicts X_0_ at every timestep). In many cases, trajectories can be stopped at timestep 50-75 with little effect on the final success rate of designs (left), and when normalized by compute budget (right), success rates per unit time are typically higher generating more designs with early-stopping. For example, in the *6EXZ_Long* benchmarking motif-scaffolding problem, stopping trajectories at t=100 allows the generation of 128 *in silico* successful designs in the time it takes to generate 42 successful designs running full trajectories.

**Figure S4:**
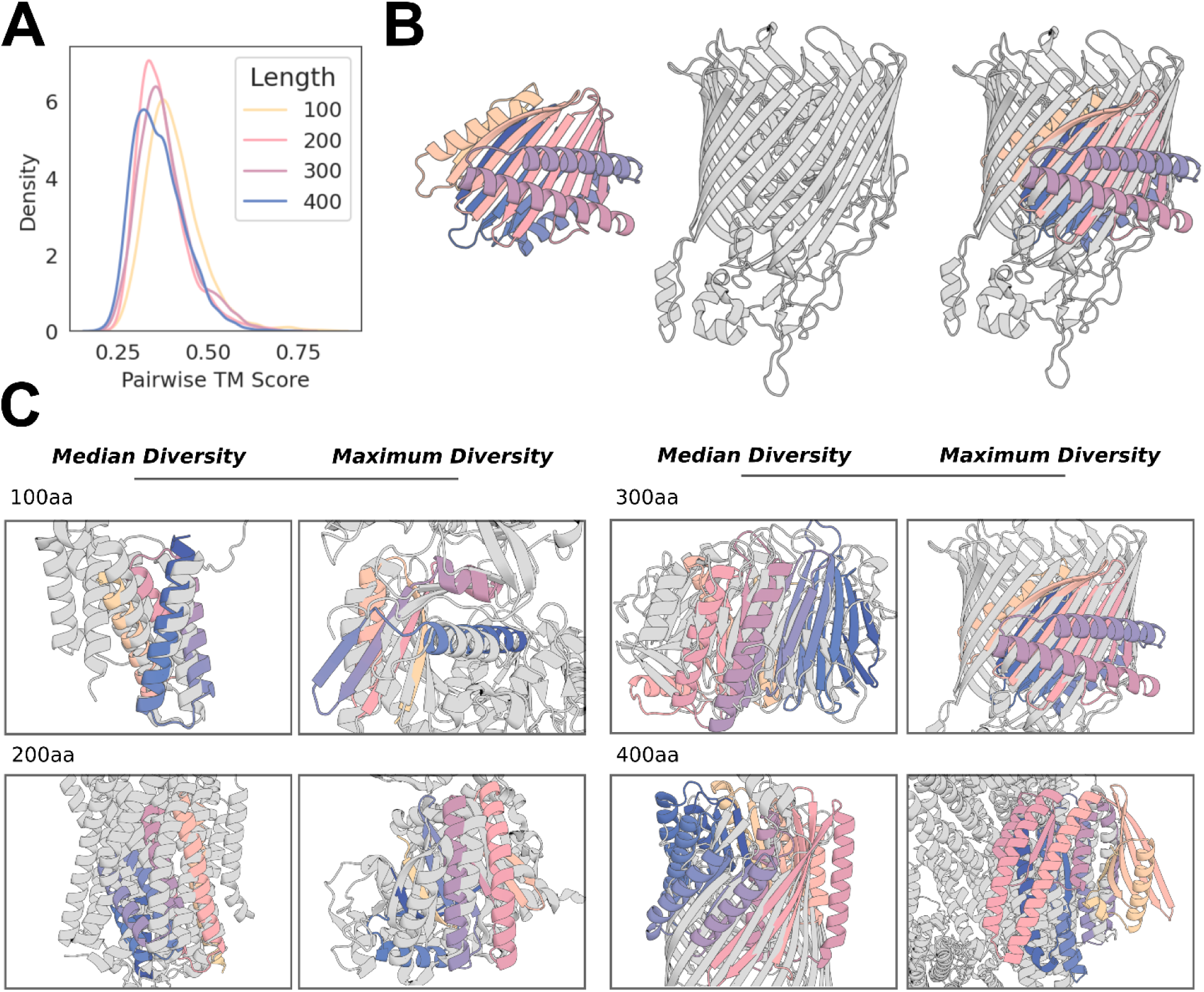
RF*diffusion* designs are diverse and dissimilar to proteins in the PDB. **A)** Comparing unconditional designs to one another (100 designs per length) demonstrates that, by TM score alignment, designs are diverse (medians 100-400aa: 0.39, 0.36, 0.37, 0.35). **B-C)** Designs also bear little resemblance to the training set (PDB). **B)** Example of the most diverse (lowest TM score hit) to the PDB for a set of 300 amino acid designs. The folds of the design (left) and native protein (middle) are highly dissimilar, aligning only across a portion of the -sheet. **C)** Example designs demonstrating extrapolation beyond the training set for generating novel folds. Gray: closest protein in the PDB by TM score, colors: RF*diffusion* design model, overlaid by TM alignment. For each protein length, the median and most diverse samples are shown (the 300aa design is the same as in **B**). While for short proteins, designs typically show some similarity to known protein folds, with increasing length, designs become increasingly dissimilar to the PDB. TM score (closest PDB, TM score; median, most diverse): 100aa: 5WVE_A, 0.71; 4W5T_A, 0.59; 200aa: 4AV3_A, 0.58; 4CLY_A, 0.47; 300aa: 4PEW_B, 0.53; 4RDR_A, 0.46; 400aa: 4AIP_A, 0.49; 6R9T_A, 0.42.

**Figure S5:**
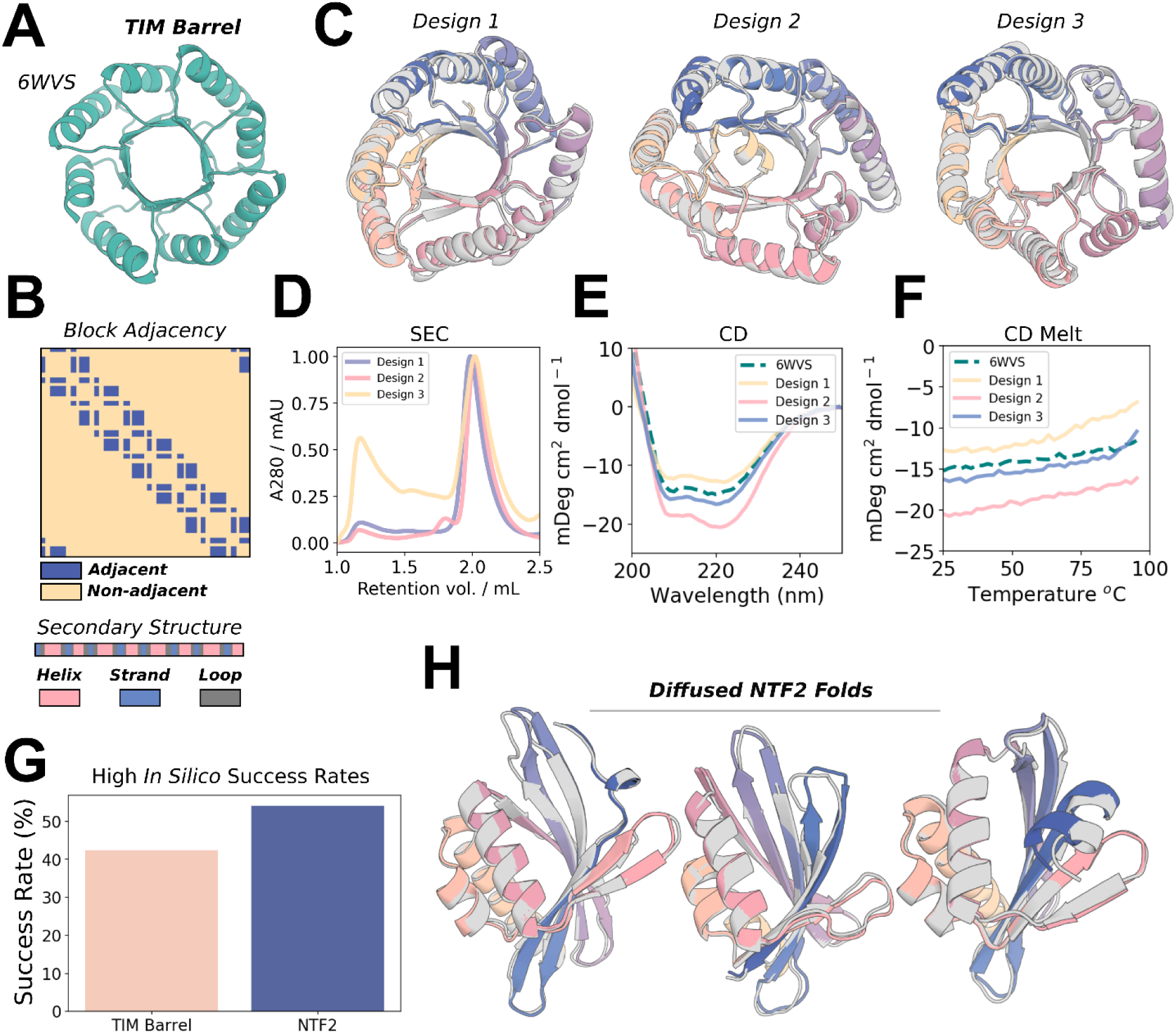
RF*diffusion* can condition on fold information to generate specific scaffold types. **A-B)** 6WVS is a previously-described *de novo* designed TIM barrel (left). A fine-tuned RF*diffusion* model can condition on 1D and 2D inputs representing this protein fold, specifically secondary structure (**B**, bottom) and block-adjacency information (**B**, top) (see Methods 4.3.2). **C)** RF*diffusion* readily conditions on fold information and generates a diverse set of TIM barrels. **D-F)** Purification of the three designs depicted in (**C**) show elution at the predicted volume (**D**), circular dichroism (CD) spectra very similar to 6VWS (**E**), and very high thermal stability (**F**). Note that **E)** and **F**) are reproduced from Fig. 2H, for clarity. **G)** TIM barrels are generated with an *in silico* success rate of 42.5% (left bar). Success incorporates AF2 metrics and a TM score vs 6WVS > 0.5. **G-H)** NTF2 folds are useful scaffolds for *de novo* enzyme design, and can also be readily generated with fold-conditioning in RF*diffusion*. Designs are diverse (**H**) and designed with an *in silico* success rate of 54.1% (**G**, right bar). NTF2 fold design success also included both AF2 metrics and a TM score vs PDB: 1GY6 > 0.5. Gray: RF*diffusion* design, colors: AF2 prediction.

**Figure S6:**
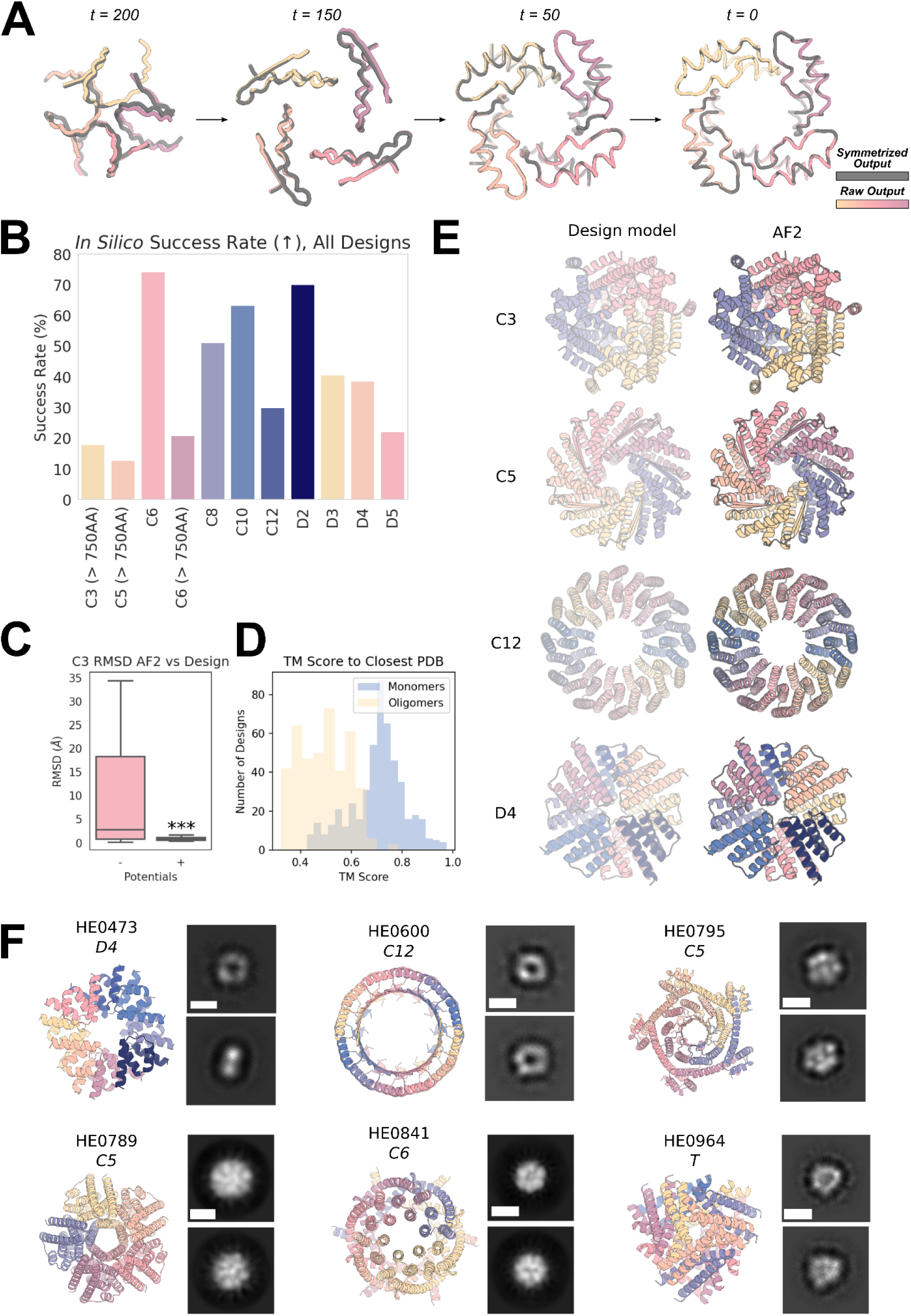
Symmetric oligomer design with RF*diffusion*. **A)** Due to the (near-perfect - see Methods 3.1) equivariance properties of RF*diffusion*, X_0_ predictions from symmetric inputs are also symmetric, even at very early timepoints (and becoming more symmetric through time; RMSD vs symmetrized: *t=200* 1.20Å; *t=150* 0.40Å; *t=50* 0.06Å; *t=0* 0.02Å). Gray: symmetrized (top left) subunit; colors: RF*diffusion* X_0_ prediction. **B)** *In silico* success rates for symmetric oligomer designs of various cyclic and dihedral symmetries. Success is defined here as the proportion of designs for which AF2 yields a prediction from a single sequence that has mean pLDDT > 80 and backbone RMSD over the oligomer between the design model and AF2 < 2 Å. Note that 16 sequences per RF*diffusion* design were sampled. **C)** Box plots of the distribution of backbone RMSDs between AF2 and the RF*diffusion* design model with and without the use of external potentials during the trajectory. The external potentials used are the “inter-chain” contact potential (pushing chains together), as well as the “intra-chain” contact potential (making chains more globular). Using these potentials dramatically improves *in silico* success (Student’s unpaired t-test, *p<0*.*001*). **D)** Designs are diverse with respect to the training dataset (the PDB). While the monomers (typically 60-100aa) show reasonable alignment to the PDB (median 0.72), the whole oligomeric assemblies showed little resemblance to the PDB (median 0.50). **E)** Additional examples of design models (left) against AF2 predictions (right) for C3, C5, C12, and D4 symmetric designs (the symmetries not displayed in Fig. 3) with backbone RMSDs against their AF2 predictions of 0.82, 0.63, 0.79, and 0.78 with total amino acids 750, 900, 960, 640. **F)** Additional nsEM data for symmetric designs. The model is shown on the left and the 2D class averages on the right for each design. Scale bars shown (white) are 60 Å.

**Figure S7:**
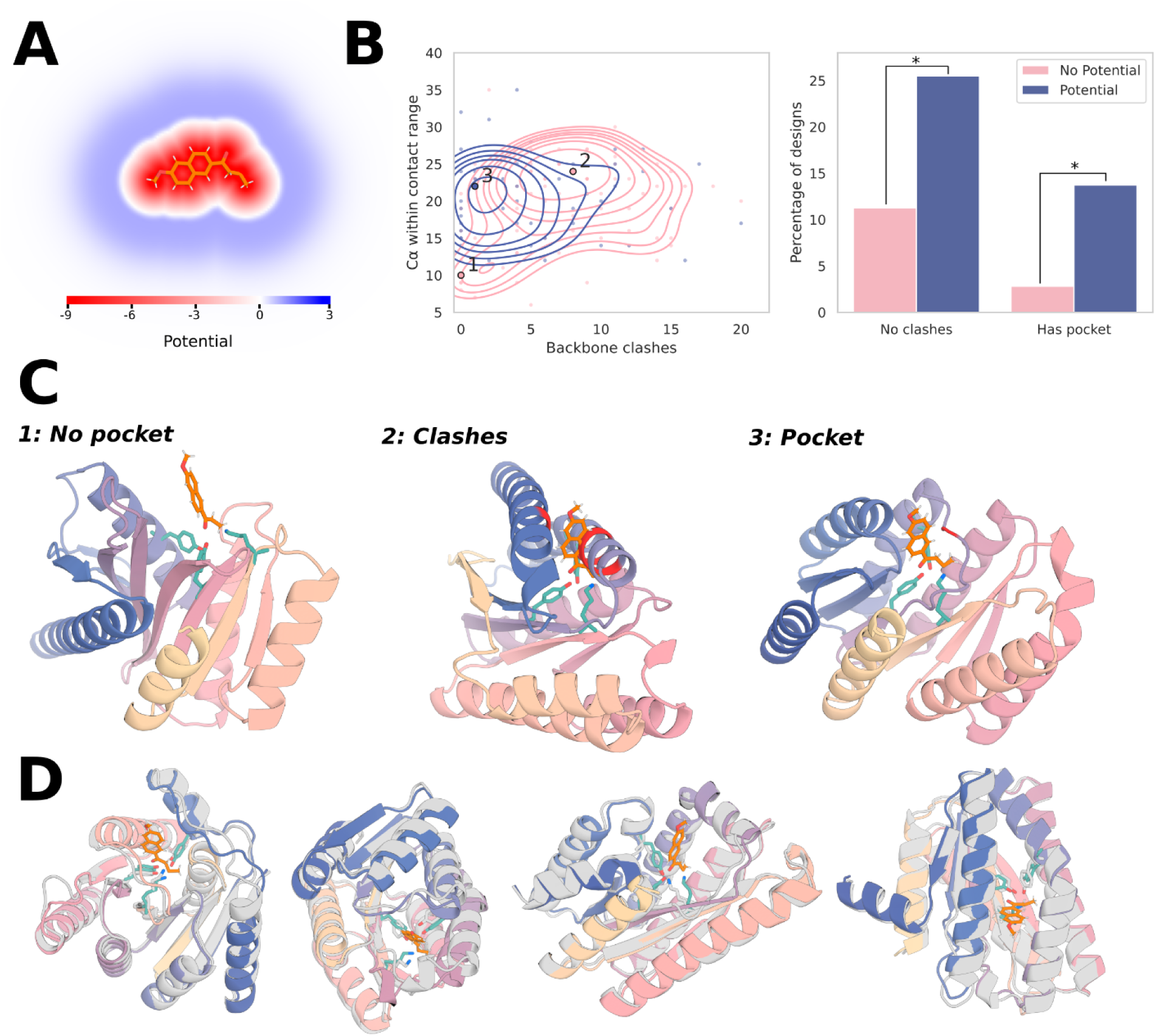
External potentials for generating pockets around substrate molecules. Enzymes generated from a retroaldolase active site triad [TYR1051-LYS1083-TYR1180] of a retro-aldolase: PDB: 5AN7. **A)** The potential used to implicitly model the substrate, which has both a repulsive and attractive field (see Methods 4.4). **B)** Left: Kernel densities demonstrate that without using the external potential (pink), designs often fall into two failure modes: (1) no pocket, and (2) clashes with the substrate. Right: clashes (substrate < 3A of the backbone) & pockets (no clash and > 16 Cα within 3-8A of substrate) with and without the potential. Two-proportion z-test: clashes *p<0*.*03*, pocket *p<0*.*02*. Each datapoint represents a design already passing the stringent success metrics (AF2 motif RMSD < 1Å, AF2 backbone RMSD < 2Å, AF2 pAE < 5). **C)** Designs close to the labeled local maxima of the kernel density estimate. Without the potential, the catalytic triad is predominantly (*1*) exposed on the surface with no residues available to provide substrate stabilization or (*2*) buried in the protein core, preventing substrate access. With the potential, the catalytic triad is predominantly (*3*), partially buried in a concave pocket with shape complementary to the substrate. Backbone atoms within 3Å of the substrate are shown in red. **D)** A variety of diverse designs with pockets made using the potential, with no clashes between the substrate and the AF2-predicted backbone. The functional form and parameters used for the pocket potential are discussed in Methods 4.4. In each case the substrate is superimposed on the AF2 prediction of the catalytic triad.

**Figure S8:**
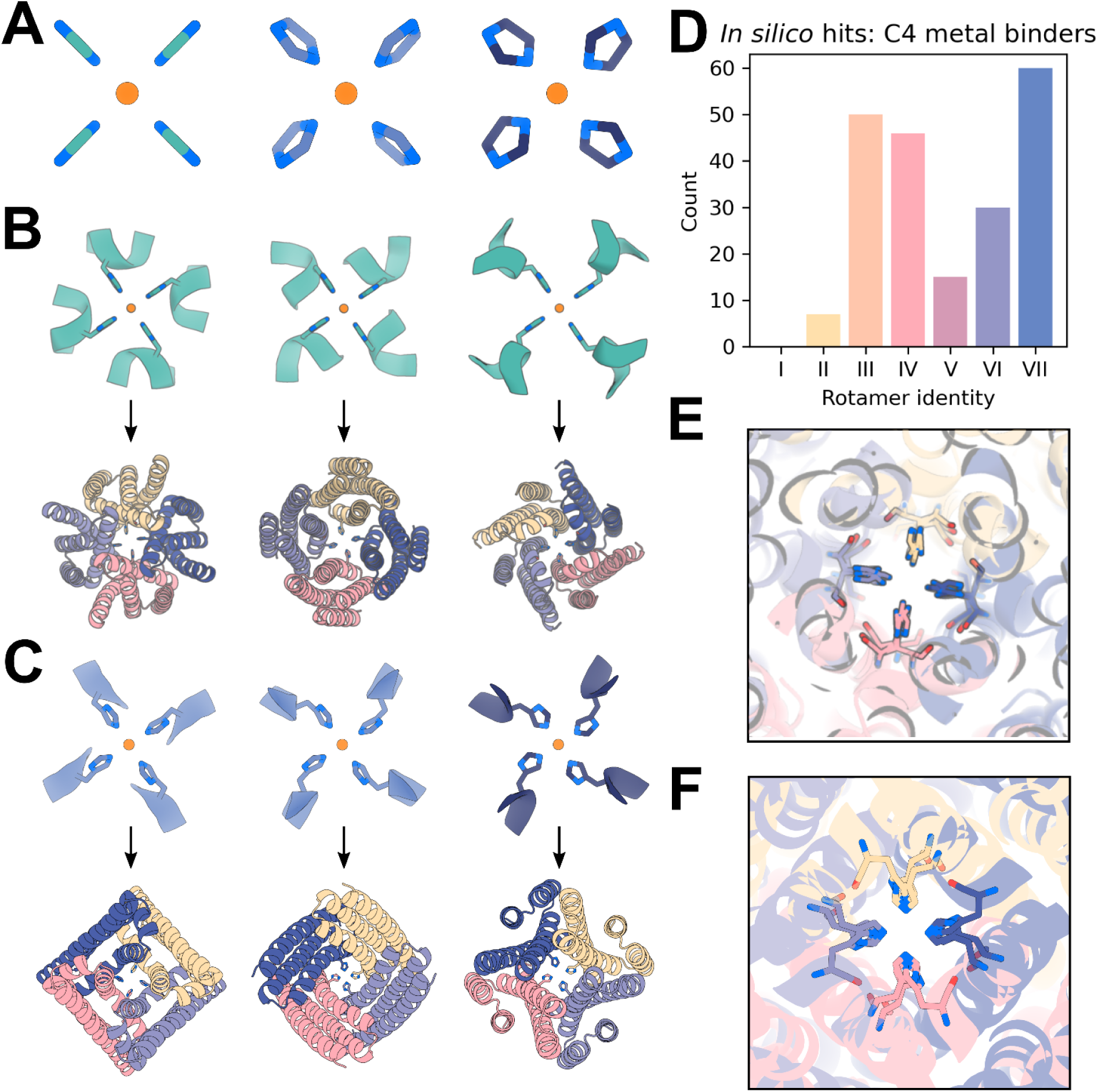
Symmetric motif scaffolding for square-planar Ni^2+^ binding. **A)** Symmetrized imidazole groups of varying amounts of shear used for constructing the square-planar motifs to scaffold, with 2.2 Å between the theoretically Ni^2+^ coordinating nitrogen and the symmetry axis. Depiction of a subset of the C4-symmetrized backbone-dependent (***φ*** = −40°, ***ψ*** = −60°) rotamers^51^ (“inverse rotamers”, Methods 5.9) used as motifs from set #1 input to RF*diffusion* for symmetrically scaffolding the theoretical Ni^2+^ binding site (teal, top). AF2 predictions of selected *in silico* successes scaffolding the C4 inverse rotamers show significant structural diversity in RF*diffusion* solutions (colors, bottom). All AF2 structures have full-atom RMSD < 1.0 Å between AF2 predictions and the input motif, AF2 PAE < 6, and AF2 pLDDT > 90. **C)** Depiction of a different subset of the C4-symmetrized backbone-dependent (***φ*** = −40°, ***ψ*** = −60°) inverse rotamers^51^ used as motifs from sets #2 and #3 (top), with AF2 predictions of selected *in silico* successes (bottom). All AF2 structures have full-atom RMSD < 1.0 Å between AF2 predictions and the input motif, AF2 PAE < 6, AF2 pLDDT > 90 **D)** *In silico* success count for the inverse rotamers from set #1 depicted in panel B. An *in silico* “success” here is defined as an AF2 prediction for a single sequence which has (1) full-atom RMSD over the four histidine residues between the AF2 prediction and the ideal C4 motif of < 1.0 Å and (2) an AF2 pAE < 10. **E)** Overlay of various AF2 predictions for designs scaffolding motifs derived from imidazole groups with no shear (panel A, left) shows a diverse array of RF*diffusion* solutions can all place the histidine imidazole groups at near-ideal distances from a theoretical nickel ion. **F)** Overlay of various AF2 predictions for motifs derived from imidazole groups with shear (panel A, middle and right) again displays diverse backbone solutions for placing the imidazole groups at near-ideal distances from the theoretical Ni^2+^ ion.

**Figure S9:**
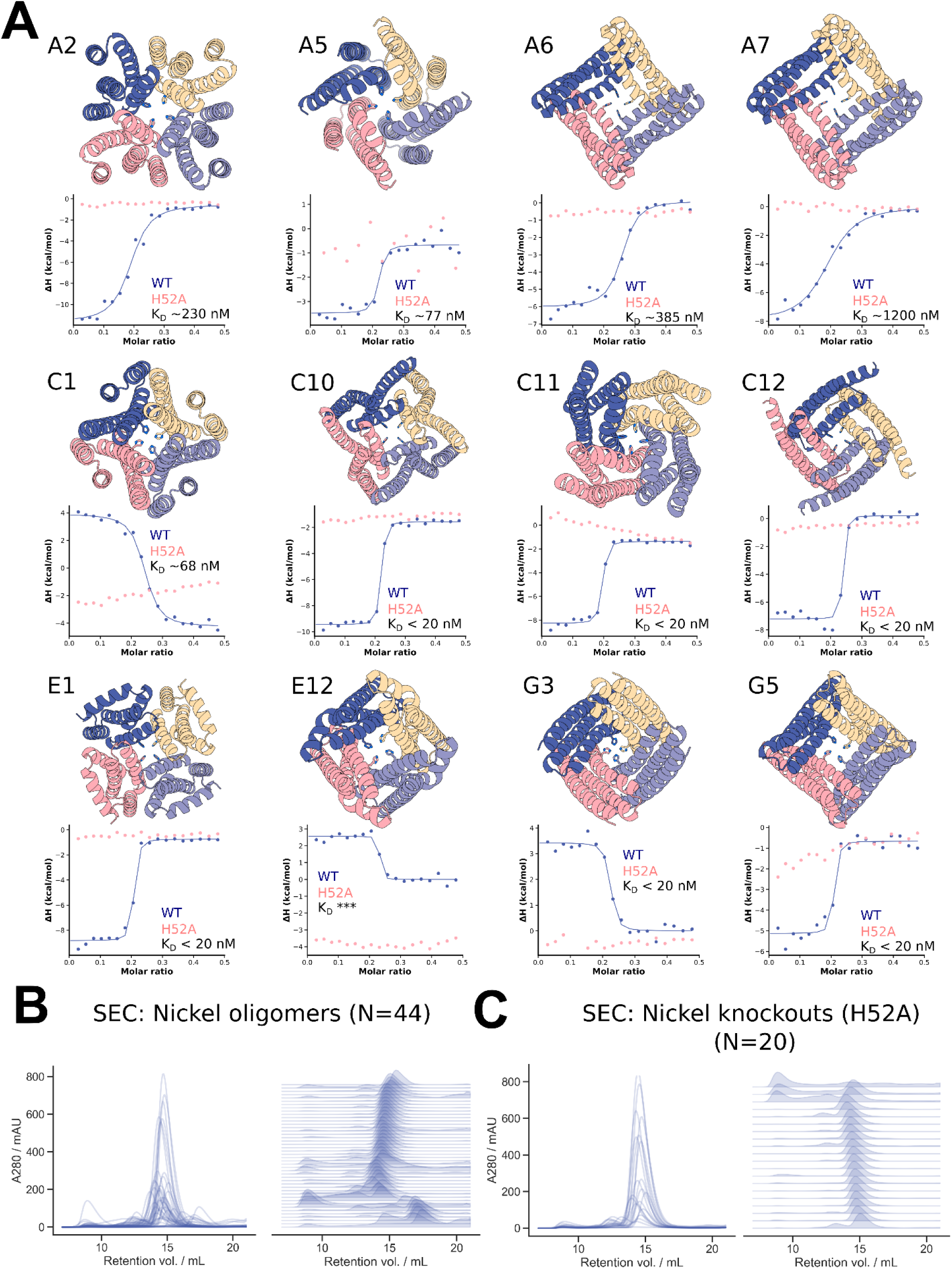
Additional Ni^2+^ binding C4 oligomers. **A)** AF2 predictions of a subset of the experimentally verified Ni^2+^ binding oligomers, with corresponding isothermal titration calorimetry (ITC) binding isotherms for the wild-type (blue) and H52A mutant (pink) below. Wild-type dissociation constants are displayed in each plot. We observe a mixture of endothermic (C1, E12, G3) and exothermic isotherms. For all cases displayed we observe no binding to the ion for H52A mutants, indicating the scaffolded histidine at position 52 is critical for ion binding. Kd values in the isotherms indicate binding of the ion with the designed stoichiometry (1:4 Ni^2+^:protein). Note that each backbone depicted is from a unique RF*diffusion* sampling trajectory, and that models and data for designs G3, C10, A5 and E1 from Figure 5 are duplicated here for ease of viewing. **B)** Size exclusion chromatograms for elutions from the 44 purifications indicate the vast majority of designs are soluble and have the correct oligomeric state. **C)** Size exclusion chromatograms for 20 H52A mutants show that the mutants remain soluble and retain the intended oligomeric state.

**Figure S10:**
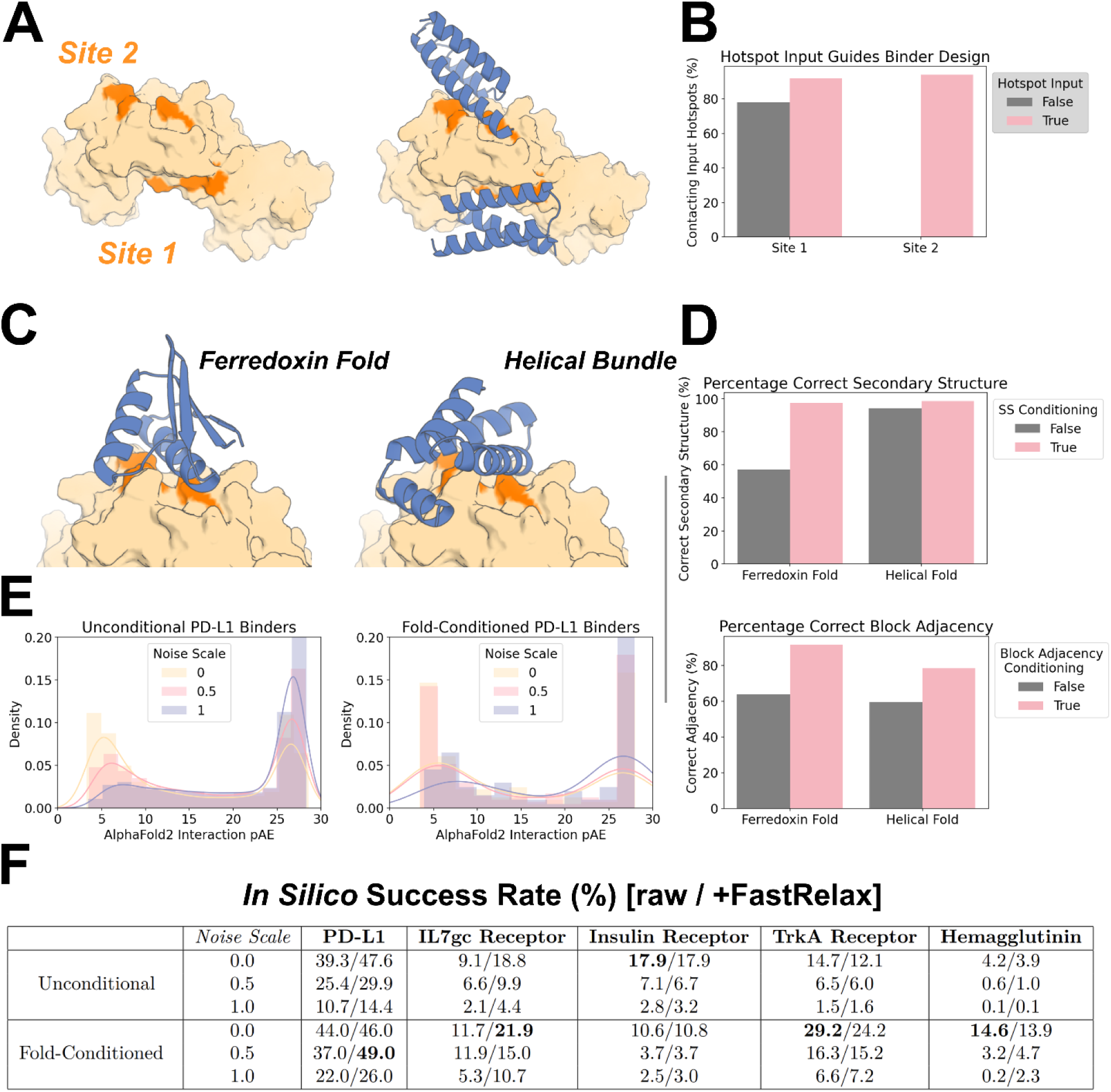
Targeted unconditional and fold-conditioned protein binder design. **A-B)** The ability to specify where on a target a designed binder should bind is crucial. Specific “hotspot” residues can be input to a fine-tuned RF*diffusion* model, and with these inputs, binders almost universally target the correct site. **A)** IL-7Ra (PDB: 3DI3) has two patches that are optimal for binding, denoted Site 1 and Site 2 here. For each site, 100 designs were generated (without fold-specification). **B)** Without guidance, designs typically target Site 1 (left bar, gray), with contact defined as C_ɑ_-C_ɑ_ distance between binder and hotspot reside < 10Å. Specifying Site 1 hotspot residues increases further the efficiency with which Site 1 is targeted (left bar, pink). In contrast, specifying the Site 2 hotspot residues can completely redirect RF*diffusion*, allowing it to efficiently target this site (right bar, pink). **C-D)** As well as conditioning on hotspot residue information, a fine-tuned RF*diffusion* model can also condition on input fold information (secondary structure and block-adjacency information - see Methods 4.5). This effectively allows the specification of a (for instance, particularly compatible) fold that the binder should adopt. **C)** Two examples showing binders can be specified to adopt either a ferredoxin fold (left) or a particular helical bundle fold (right). **D)** Quantification of the efficiency of fold-conditioning. Secondary structure inputs were accurately respected (top, pink). Note that in this design target and target site, RF*diffusion* without fold-specification made generally helical designs (right, gray bar). Block-adjacency inputs were also respected for both input folds (bottom, pink). **E)** Reducing the noise added at each step of inference improves the quality of binders designed with RF*diffusion*, both with and without fold-conditioning. As an example, the distribution of AF2 interaction pAEs (known to indicate binding when pAE < 10) is shown for binders designed to PD-L1. In both cases, the proportion of designs with interaction pAE < 10 is high (blue curve), and improved when the noise is scaled by a factor 0.5 (pink curve) or 0 (yellow curve). **F)** Full *in silico* success rates for the protein binders designed to five targets. In each case, the best fold-conditioned results are shown (i.e. from the most target-compatible input fold), and the success rates at each noise scale are separated. In line with current best practice^26^, we tested using Rosetta FastRelax^52^ before designing the sequence with ProteinMPNN, but found that this did not systematically improve designs. Success is defined in line with current best practice^26^: AF2 pLDDT of the monomer > 80, AF2 interaction pAE < 10, AF2 RMSD monomer vs design < 1Å.

**Figure S11:**
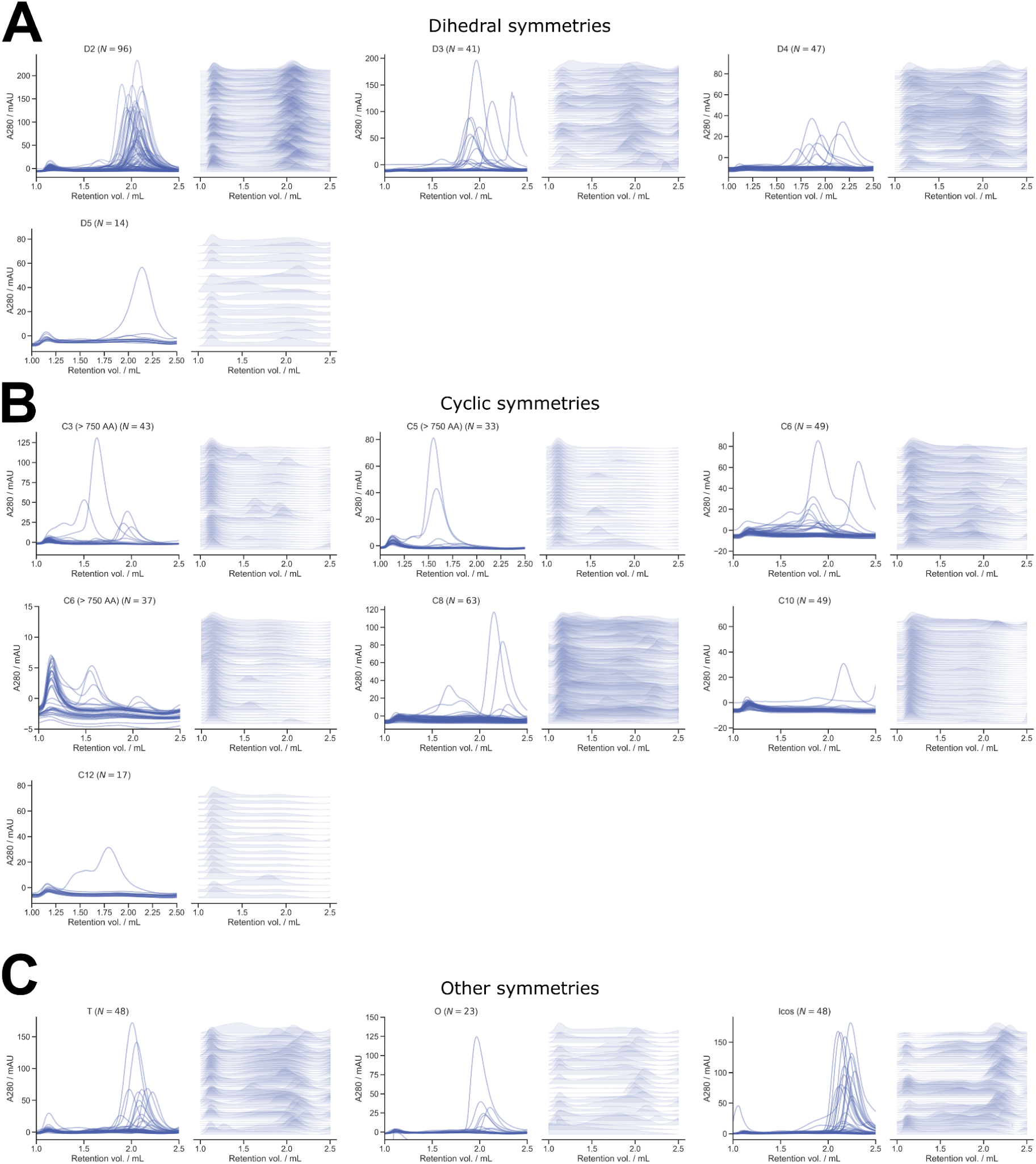
Size exclusion chromatography of symmetric oligomers. **A-C)** Size exclusion chromatography (SEC) was used as a primary screening method for all RF*diffusion*-generated oligomers. Here, SEC traces from 608 oligomers are shown for each of the experimentally tested symmetry groups, excluding the void volume. Panel **A)** shows dihedral symmetries, **B)** shows cyclic symmetries, and **C)** shows all others. For each set of traces, on the left, data are overlaid for all designs, and on the right, traces are normalized and stacked. As designs increase in complexity (higher number of individual subunits), the amount of soluble protein shown by SEC visibly decreases. For tetrahedral, octahedral, and icosahedral designs, many have soluble protein peaks that are possibly dimer and trimer subunits (unassembled cages).

**Figure S12:**
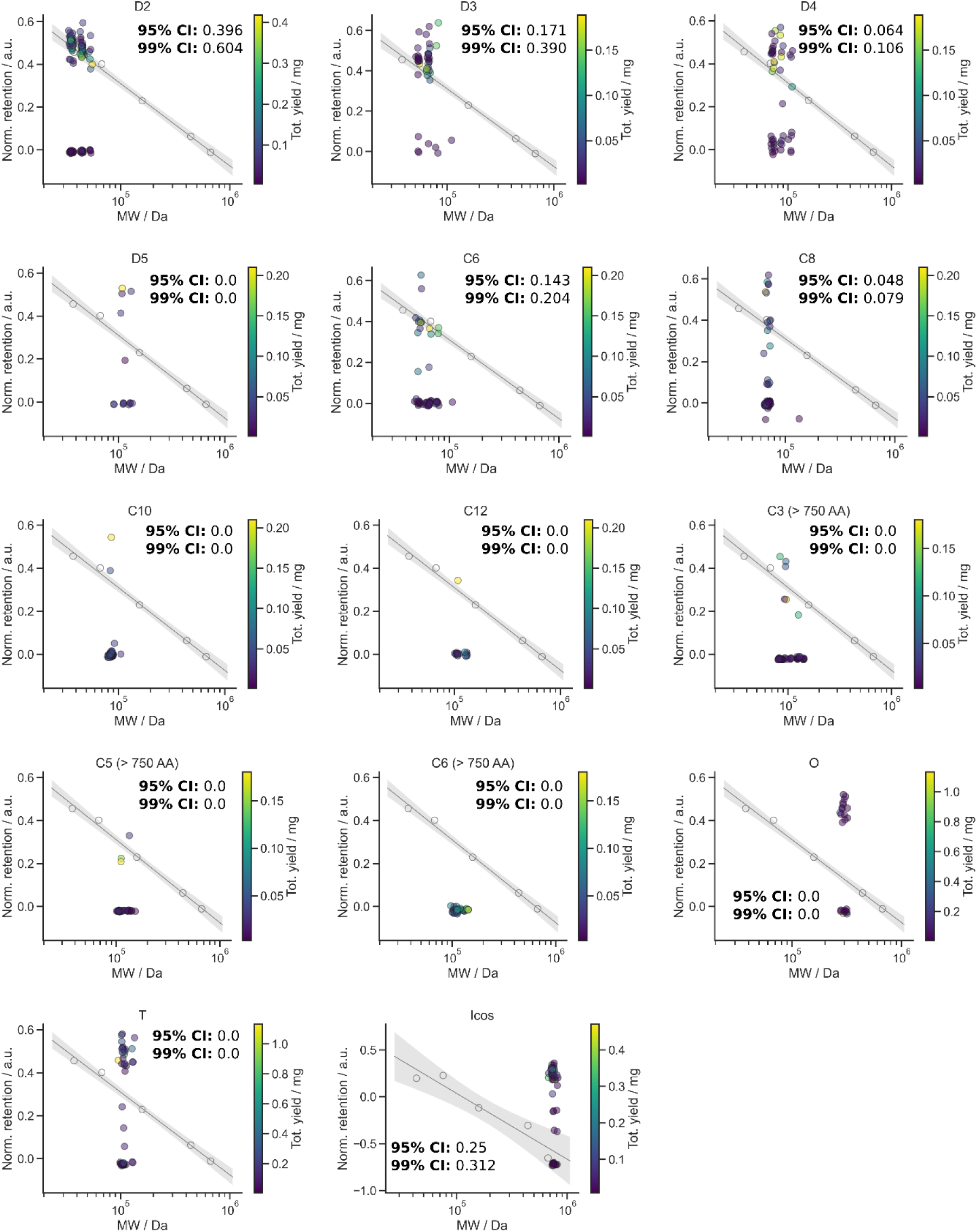
SEC elution peaks of symmetric oligomers vs. calibration curves. Retention volume for the major SEC peak versus molecular weight for each design are plotted in comparison to a known calibration curve. The calibration curve is shown in gray, with shading representing the 95% confidence interval. Total yield of each design is indicated by the scale bar on the right of the graphs, and success rates for the 95% CI and 99% CI are denoted on each graph per each symmetry. Given that MW is being used as a proxy for hydrodynamic radius, we expect that some designs (e.g. cycles with large pores) may be true to their design model, but deviate from the standard curve. These calibration curves provide a rough estimate of the success rate of each symmetry group, and help guide the selection process for downstream analysis of any design. In some cases, even though no designs are within the 99% CI, we still selected designs to screen by nsEM. For example, we are able to confirm HE0822 (C3) by nsEM despite misalignment between the theoretical and actual elution profiles (Fig. 3B). Because of their size, the icosahedra were run on an S6 column with lower resolution; thus, the calibration curve fit results in bigger confidence intervals compared to an S200 column, which was used to screen all other oligomers (See Methods 6.2). We expect that for oligomers run on the S200, reported success rates are fairly conservative, whereas for designs run on the S6, experimental success rates are likely lower.

**Figure S13:**
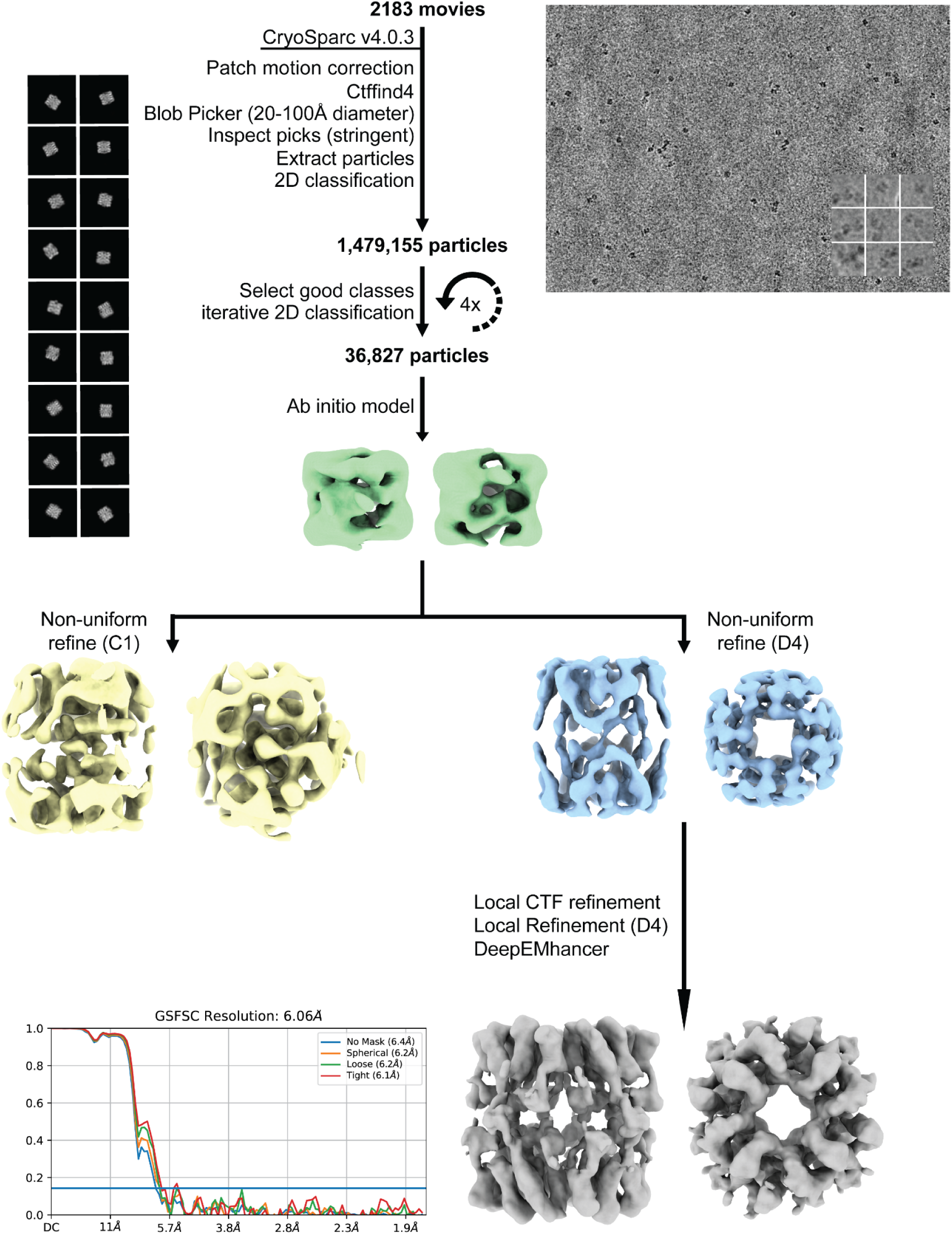
Details of HE0537 cryo-EM data processing pipeline. 2D class averages showing exclusively side-views of HE0537, and an *ab initio* reconstruction followed by a C1 non-uniform refinement yielding identifiable D4 features corresponding to the size and rough secondary structure of the design model. Further data processing was attempted with D4 symmetry imposed, but the strong preferred orientation precluded generation of a reliable 3D map for detailed structural analysis. At this time, only the predicted 2D projection images of the design model are analyzed/compared alongside the corresponding experimental cryo-EM 2D class average side views in Fig. 3D, which display strikingly high agreement to the design. A representative raw cryo-EM micrograph is shown on the right along with nine example extracted particles and characteristic 2D class averages used in the processing pipeline. An FSC validation curve for the final reconstruction is shown along with the density map.

**Figure S14:**
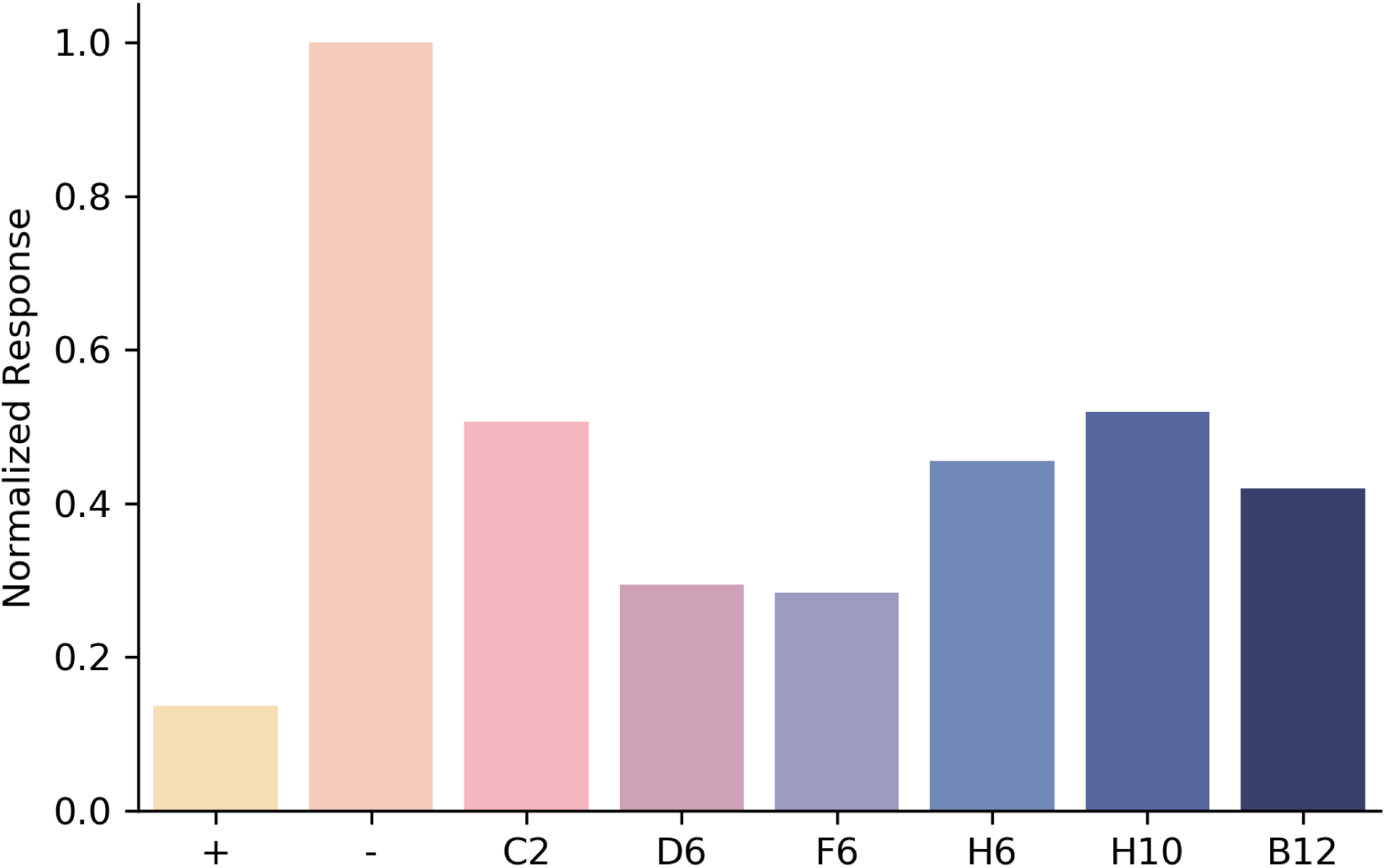
IL-7Ra Competition Assay. Positive control (known IL-7Ra binder from ref [^12^]) was amine conjugated to ar2g biosensor tips. 100nM IL-7Ra with 1μM of each design then was used as analyte. Positive control was also included as an analyte as there should be no binding. Response is normalized to binding of IL-7Ra on its own. All six diffusion-generated binders compete with the positive control, indicating they bind to the intended site.

**Figure S15:**
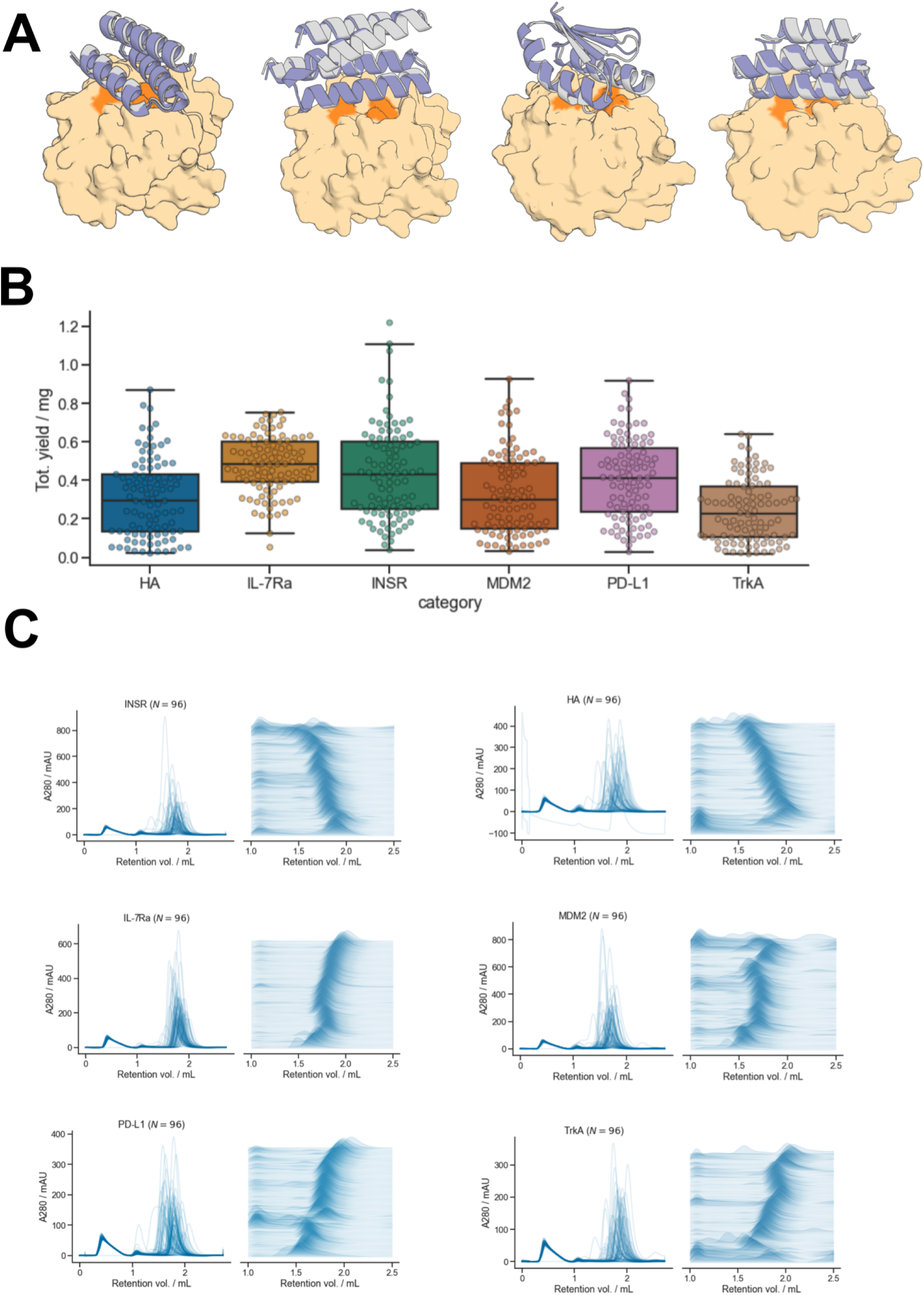
Analysis of Insulin Receptor Binding Campaign. **A)** Insulin Receptor binders are well-predicted by AF2. Yellow/orange: target/hotspot residues; gray: design model; purple: AF2 prediction. **B)** Insulin Receptor binders are expressed with high yield, in line with the rest of the binder campaigns. **C)** SEC elution profiles indicate most Insulin Receptor binders elute as monomers, in line with the rest of the binder campaigns.

