## Supplementary Methods for "Broadly applicable and accurate protein design by integrating structure prediction networks and diffusion generative models"

December 2022

### 1 RoseTTAFold: updated architecture and training details

In this section we provide an overview of relevant details of the three-track architecture of RoseTTAFold (RF) and its training. This architecture includes significant modifications as compared to the original RF [1]; these modifications will be described in full detail in a future publication, and are not a contribution of this work. We provide this section to assist in the understanding of the architecture of RF*diffusion*.

#### 1.1 Backbone structure representation

RF adopts a rigid-frame representation of the residues that comprise protein backbones. The structure of an $L$ residue backbone is described as a collection of residue frames $x = [x_1, \ldots, x_L]$, where each $x_l = (r_l, z_l)$ describes the translation $z_l \in \mathbb{R}^3$ and rigid rotation $r_l$ of the $l^{th}$ residue. In particular, each $z_l$ represents the coordinates of the $l^{th}$ $C_\alpha$ carbon, and each $r_l$ is a $3 \times 3$ rotation matrix that maps an axis-aligned residue with idealized geometry (i.e. bond lengths and angle) and its $C_\alpha$ at the origin to the positions of these atoms relative to the $C_\alpha$. For any backbone atom coordinates ($z_{C_\alpha}, z_C$ and $z_N$) for a given residue we may apply a Gram-Schmidt process to compute a $3 \times 3$ rotation matrix $r$ with rows

$$\begin{aligned} r_1 &= (z_C - z_{C_\alpha})/\|z_{C_\alpha} - z_C\|_2, \\ r_2 &= \left((z_N - z_{C_\alpha}) - ((z_N - z_{C_\alpha}) \cdot r_1)r_1\right)/\|(z_N - z_{C_\alpha}) - ((z_N - z_{C_\alpha}) \cdot r_1)r_1\|, \text{ and} \\ r_3 &= r_1 \times r_2, \end{aligned} \tag{1}$$

where $\cdot$ and $\times$ are the dot- and cross-products, respectively. 3D backbone coordinates can then be reconstructed by multiplication of idealized coordinates (with $z^*_{C_\alpha}$ at the origin,

$z_C^* - z_{C_\alpha}^*$ along the x-axis, and $z_N^* - z_{C_\alpha}^*$ in the xy-plane) by $r$ as

$$[z_C, z_N, z_{C_\alpha}] = r[z_C^*, z_N^*, z_{C_\alpha}^*] + z_{C_\alpha}\vec{1}_3,$$

where $\vec{1}_3 = [1, 1, 1]$. Accordingly, modeling the coordinates of a triplet of backbone atoms is equivalent to modeling the $C_\alpha$ coordinate $z$ and the rotation matrix $r$.

##### 1.2 RoseTTAFold architecture

RF includes multiple architectural improvements from the original RoseTTAFold network [1]: 1) the 3D structure track now extends throughout the entire network, with coordinates initialized from a template structure; 2) structure-biased axial attention is used to update 2D pair features, which considers geometric constraints between residues inferred from the current 3D structure; 3) a similar biased axial attention is used to update the 1D track, where the 2D and 3D tracks are used to bias the attention in the 1D sequence updates; and 4) RF uses of recycling, in which the network is executed multiple times with the updated input embeddings based on outputs from the previous cycle. RF contains two major types of architecture blocks: main three-track blocks and the final structure refinement blocks. The 3-track blocks consist of layers of biased row and column attention over the 1D and 2D features, SE(3)-equivariant layers [2, 3] to update 3D coordinates, and layers to communicate between 1D, 2D, and 3D features. The structure refinement block is based on SE(3)-equivariant network which gives refined 3D coordinates based on given 1D and 2D features. In recycle iterations, coordinates entering the 3D track are initialized from the predictions output in the previous pass through the network, and as a result of the SE(3)-equivariance of the architecture, the predictions in recycling passes update these initial coordinates with SE(3) equivariance. Following AlphaFold2 [4], in addition to predicting backbone structure updates, at each layer (3-track and fine-tuning) we predict up to 4 chi angles that define all protein sidechain atoms.

**Inputs and outputs of RF:** Before presenting the tensor input and outputs of RoseTTAFold, we introduce the relevant dimensions in these objects:

- L: The length of the query sequence
- I: The number of times the model will be executed (1 plus the number of recycles)
- T: The number of homologous structure provided to the model as templates
- N_short: The number of sequences in the truncated MSA
- N_long: The total number of sequences in the full MSA (capped at 1024)

**Multimer prediction with multiple chains:** As in the original RoseTTAFold [1] for hetero-complex structure prediction we indicate chain breaks in the positional encoding. Residue indices enter the network through the pair representation; for each pair of residues we include a sequence distance feature clipped between -32 and 32 (with sign indicating direction). To indicate breaks between chains of subunits, we increment residue indices by 100 at the start of each chain.

#### 1.3 RoseTTAFold training

RF was trained based on a mixture of datasets including 1) monomer/homo-oligomer structures in the PDB, 2) hetero-oligomer structures in the PDB (date cutoff August 2nd, 2021), 3) AlphaFold2 structural models having pLDDT $> 0.758$, and 4) negative protein-protein interaction examples generated by random pairing. The training examples were sampled from each database with a ratio of 2:1:4:1. The model was trained using the masked language model ($\mathcal{L}_{\text{MLM}}$) loss, distogram prediction loss $\mathcal{L}_{\text{dist}}$, FAPE loss $\mathcal{L}_{\text{FAPE}}$, accuracy estimation loss $\mathcal{L}_{\text{accuracy}}$, bond geometry loss $\mathcal{L}_{\text{bond}}$ and van der Waals (vdW) energy loss

| Input name (Shape) | Description |
|---|---|
| msa_masked (I, N_short, L, 48) | The truncated MSA with some portions of sequence masked (20aa, 1 unknown, 1 mask, MSA profiles (22), insertion/deletions (2), N-term/C-term (2)) |
| msa_full (I, N_long, L, 25) | The full length MSA (20aa, 1 unknown, 1 mask, insertion (1), N-term/C-term (2)) |
| seq (I, L, 22) | The sequence whose structure is being predicted (20aa, 1 unknown, 1 mask) |
| xyz_prev (L, 27, 3) | The structure recycling information. In recycle steps, this contains the model's previous prediction. In the first iteration (before recycling), this feature is populated with template structure coordinates if available. Otherwise, this feature set to all zeros. $N$-$C_\alpha$-$C$-$O$ backbone (4), (up to) 10 sidechain atoms, (up to) 13 hydrogen atoms) |
| idx_pdb (L) | The integer index of each residue. Used to assign each residue its neighboring residue. |
| t1d (T, L, 22) | The one-dimensional features associated with each template structure. (20 amino acids, missing template token (1), template-match confidence (1)) |
| t2d (T, L, L, 44) | The two-dimensional features associated with each template structure. (36 distance bins (2-20Å, 0.5Å bins) + 1 final distance bin ($>$ 20Å), angle maps (sine and cosine of omega, theta and phi angle) (6), missing residue mask (1)) |
| xyz_t (T, L, 27, 3) | The structure of the template structures. This feature is immediately converted to a distogram and anglegram representation by the model. (N, Ca, C backbone atoms) |
| alpha_t (T,L,10*3) | The backbone and sidechain torsion angles of each residue of the template structures. Initially T, L, 10, 2, with sine and cosine of (omega, phi, psi angles (3), (up to) 4 torsion angles, $C_\beta$ bend (1), $C_\beta$ twist (1), $C_\gamma$ bend (1)). This is concatenated with a mask (T, L, 10, 1) indicating which torsion angles are present for a given amino acid, and reshaped to T, L, 30. |
| msa_prev (N_short, L, Cm) | The MSA embedding recycling information. This is the model's previous embedding at each position in the truncated MSA. Cm = 256 |
| pair_prev (L, L, Cp) | The 2-D embedding recycling information. This is the model's previous embedding at each edge between each node. Cp = 128 |
| state_prev (L, Cs) | The 1-D embedding recycling information. This is the model's previous embedding at each position in the query sequence. Cs = 16 |

Table 1: Description of features input to RoseTTAFold.

| **Output name** (shape) | **Description** |
|---|---|
| `msa` (N_short, L, Cm) | The model's final embedding at each position in the truncated MSA. Cm = 256 |
| `pair` (L, L, Cp) | The model's final embedding at each edge between each node. Cp = 128 |
| `state` (L, L, Cs) | The model's previous embedding at each position in the query sequence. Cs = 16 |
| `xyz` (L, 27, 3) | The model's prediction of the structure. (N-Ca-C-O backbone (4), (up to) 10 sidechain atoms, (up to) 13 hydrogen atoms) |
| `alpha` (L, 10 * 3) | The model's prediction of sidechain torsions. Initially T, L, 10, 2, with sine and cosine of (omega, phi, psi angles (3), (up to) 4 torsion angles, $C_\beta$ bend (1), $C_\beta$ twist (1), $C_\gamma$ bend (1)). This is concatenated with a mask (T, L, 10, 1) indicating which torsion angles are present for a given amino acid, and reshaped to T, L, 30. |
| `logits_aa` (N_short , L , 21) | The model's prediction of the unmasked, truncated MSA. |
| `pred_lddt` (L) | The model's prediction of the LDDT error of each residue. |
| `logits_dist` (Cdist,L,L) | The model's prediction of the binned distances $d_{l,l'}$ between residue pairs, where $d_{l,l'}$ is the Euclidean distance $C_{\beta,l}$ and $C_{\beta,l'}$. Cdist = 37. |
| `logits_omega` (Cdist,L,L) | The model's prediction of the binned $\omega$ angles between residue pairs, where $\omega_{l,l'}$ is the $C_{\alpha,l}$-$C_{\beta,l}$-$C_{\alpha,l'}$-$C_{\beta,l'}$ dihedral angle. |
| `logits_theta` (Cdist,L,L) | The model's prediction of the binned $\theta$ angles between residue pairs, where $\theta_{l,l'}$ is the $N_l$-$C_{\alpha,l}$-$C_{\beta,l}$-$C_{\beta,l'}$ dihedral angle. |
| `logits_phi` (Cphi,L,L) | The model's prediction of the binned $\phi$ angles between residue pairs, where $\phi_{l,l'}$ is the pseudo-bond angle dictating the direction of $C_{\beta,l'}$ from residue $l$'s frame of reference. Cphi = 19. |

Table 2: Description of outputs returned by RoseTTAFold.

| | **Initial training** | **Fine-tuning** |
|---|---|---|
| Crop size | 256 | 384 |
| Batch size | 64 | 64 |
| Loss function | $3.0*\mathcal{L}_{\text{MLM}} + 1.0*\mathcal{L}_{\text{dist}} + 10.0*\mathcal{L}_{\text{FAPE}} + 0.1*\mathcal{L}_{\text{accuracy}}$ | $3.0*\mathcal{L}_{\text{MLM}} + 1.0*\mathcal{L}_{\text{dist}} + 10.0*\mathcal{L}_{\text{FAPE}} + 0.1*\mathcal{L}_{\text{accuracy}} + 0.1*\mathcal{L}_{\text{bond}} + 0.1*\mathcal{L}_{\text{vdW}}$ |
| Learning rate, & scheduling | 0.001, Linear warm-up for first 1000 optimization steps, then decay learning rate by 0.95 after every 15000 optimization steps | 0.0005, No warm-up. Decay learning rate by 0.95 after every 15000 optimization steps |
| Examples per epoch | 25600 | 25600 |
| Number of epochs | 200 | 100 |

Table 3: RoseTTAFold training hyperparameters.

$\mathcal{L}_{\text{vdW}}$. For the initial round of training, only the first four loss terms were used with crop size 256. After 200 epochs of initial round training, we performed fine-tuning with all the loss terms with crop size 384 for 100 epochs. RoseTTAFold was trained for 4 weeks on 64 V100 GPUs on Microsoft Azure. The training details are summarized in Table 3.

### 2 RF*diffusion*: principles and formulation as a generative model of structure

This section details how we have repurposed RoseTTAFold (RF) as the neural network in a diffusion model of protein backbones. When it is clear from context, we sometimes drop the residue subscript, and write a residue frame as $(r, z)$. Section 2.1 reviews denoising diffusion probabilistic models (DDPMs) to establish notation and terminology. RF*diffusion* adapts DDPMs to the rigid-frame representation of residues used by RF (as described in Section 1.1), and Sections 2.2 and 2.3 describe the forward and reverse processes for translation and rotation components of this representation. Section 2.4 introduces self-conditioning in RF*diffusion*. Section 2.5 presents the mean-squared-error denoising loss used in training, and discusses how minimizing this objective relates to learning the reverse process. Finally, Section 2.6 discusses rotational invariance in RF*diffusion*.

#### 2.1 Diffusion probabilistic modeling and rotational equivariance

DDPMs [5, 6] are a class of generative models that intend to reverse a discrete-time diffusion process. The forward process starts with a sample $x^{(0)} \sim q(x^{(0)})$ from an unknown data distribution $q$, to which we have access only though samples. The data are corrupted at each of $T$ steps, to obtain a sequence of increasingly noisy samples; for each $t = 1, \ldots, T$, we sample $x^{(t)} \sim q(x^{(t)} \mid x^{(t-1)})$ such that the final step $x^{(T)} \sim q(x^{(T)})$ is indistinguishable from a reference distribution that has no dependence on the data. DDPMs approximate $q(x^{(0)})$ with a second distribution $p(x^{(0)})$ parameterized by a backward transition kernel $p(x^{(t-1)} \mid x^{(t)})$ at each $t$. We train a neural network parameterizing each $p(x^{(t-1)} \mid x^{(t)})$ such that $p(x^{(t-1)}|x^{(t)})$ approximates $q(x^{(t-1)}|x^{(t)})$. One then draws from $p(x^{(0)})$ by first sampling from the reference distribution $x^{(T)} \sim p(x^{(T)}) = q(x^{(T)})$, and then for each $t < T$

repeatedly denoising by sampling $x^{(t-1)} \sim p(x^{(t-1)} \mid x^{(t)})$ until $x^{(0)} \sim p(x^{(0)})$ is obtained. In the limit that the approximation $p(x^{(t-1)}|x^{(t)})$ of $q(x^{(t-1)}|x^{(t)})$ is exact, $p(x^{(0)}) = q(x^{(0)})$.

In our case, $q(x^{(0)})$ is a distribution over a native protein backbones parameterized by residue frames. We define the forward noising process independently over the rotational and translational components of this representation. We similarly model the reverse process transitions as conditionally independent across these components given $x^{(t)}$ as

$$p(x^{(t-1)} \mid x^{(t)}) = p(r^{(t-1)}|x^{(t)})p(z^{(t-1)}|x^{(t)}).$$

We describe the details of the forward and reverse process conditionals for the translations in Section 2.2 and for the rotations in Section 2.3.

#### 2.2 Residue translations, forward and reverse transitions

Our forward process for translations follows closely from previous work by Trippe et al. [7] on $C_\alpha$ backbone generation, that treats $C_\alpha$ backbone coordinates as a 3D point cloud and corrupts them with 3D Gaussian noise. We let $\beta^{(1)}, \beta^{(2)}, \ldots, \beta^{(T)}$ be scalars between 0 and 1 that define a variance schedule such that for each $t = 1, 2, \ldots, T$ the transition density of the forward process is $q(z^{(t)} \mid z^{(t-1)}) = \mathcal{N}(z^{(t)}; \sqrt{1-\beta^{(t)}}z^{(t-1)}, \beta^{(t)} I_3)$. To sample $z^{(t)}$ during training, rather than ancestral sampling $z^{(s)}|z^{(s-1)}$ from $s = 1$ all the way up to $s = t$, we draw $z^{(t)}$ directly from the marginal distribution, $q(z^{(t)}|z^{(0)}) = \mathcal{N}\left(z^{(t)}; \sqrt{\bar{\alpha}^{(t)}}z^{(0)}, (1-\bar{\alpha}^{(t)})I_3\right)$. Where we define $\bar{\alpha}^{(t)} = \prod_{s=1}^{t} \alpha^{(t)}$ with $\alpha^{(t)} = 1 - \beta^{(t)}$.

For the reverse process, we desire to use a prediction of denoised coordinates from RoseTTAFold. Given that $q(z^{(t-1)}|z^{(t)}, z^{(0)}) = \mathcal{N}(z^{(t-1)}; \tilde{\mu}(z^{(t)}, z^{(0)}), \tilde{\beta}^{(t)} I_3)$ for $\tilde{\mu}(z^{(t)}, z^{(0)}) = \frac{\sqrt{\bar{\alpha}^{(t-1)}}\beta^{(t)}}{1-\bar{\alpha}^{(t)}}z^{(0)} + \frac{\sqrt{\alpha^{(t)}}(1-\bar{\alpha}^{(t)})}{1-\bar{\alpha}^{(t)}}z^{(t)}$, and $\tilde{\beta}^{(t)} = \frac{1-\bar{\alpha}^{(t-1)}}{1-\bar{\alpha}^{(t)}}\beta^{(t)} \approx \beta^{(t)}$, we define the reverse transitions by

$$
\begin{aligned}
& p(z^{(t-1)} \mid x^{(t)}) = \mathcal{N}(z^{(t)}; \hat{\mu}(x^{(t)}), \beta^{(t)} I_3), \\
& \text{with} \;\; \hat{\mu}(x^{(t)}) = \frac{\sqrt{\bar{\alpha}^{(t-1)}}\beta^{(t)}}{1-\bar{\alpha}^{(t)}} \hat{z}^{(0)}(x^{(t)}) + \frac{\sqrt{\alpha^{(t)}}(1-\bar{\alpha}^{(t-1)})}{1-\bar{\alpha}^{(t)}} z^{(t)},
\end{aligned}
\tag{2}
$$

where $\hat{z}^{(0)}(x^{(t)})$ denotes the predicted $C_\alpha$ coordinates obtained from RF*diffusion*, $\hat{x}^{(0)}(x^{(t)})$. We choose $\beta^{(t)}$ according to a linear variance schedule [6]. We chose these parameters such that signal remaining in $z^{(0)}$ (as quantified by $\bar{\alpha}^{(t)}$) decayed slowly toward zero as $t$ approaches $T$ (Table 6).

#### 2.3 Residue rotations, forward and reverse transitions

For the forward and reverse transitions on rotations, we adapt a generalization developed by De Bortoli et al. [8] of diffusion models to Riemannian manifolds. In particular, the space of $3 \times 3$ rotation matrices (known as the special orthogonal group of dimension 3, or $SO(3)$) is a compact Riemannian manifold where the techniques of Ho et al. [6] do not apply readily. In brief, De Bortoli et al. [8] build on the continuous-time score-based generative modeling framework of Song et al. [9] and define the forward process as Langevin dynamics on the manifold — and in particular as a Brownian motion when the manifold is compact. The time-reversal of this process is then characterized through the Stein score of the noised data distribution at each $t$.

**Forward process defined by Brownian motion on $SO(3)$:** The form of the Brownian motion on a manifold is well-defined only with the choice of a metric on and a basis vectors of the associated tangent space. In the case of $SO(3)$, the tangent space may be represented through the Lie algebra of $SO(3)$ (the set of $3 \times 3$ skew symmetric matrices); we choose a metric defined through the Frobenius inner product, on adopt the following basis vectors:

$$\begin{bmatrix} 0 & -\frac{1}{\sqrt{2}} & 0 \\ \frac{1}{\sqrt{2}} & 0 & 0 \\ 0 & 0 & 0 \end{bmatrix}, \quad \begin{bmatrix} 0 & 0 & \frac{1}{\sqrt{2}} \\ 0 & 0 & 0 \\ -\frac{1}{\sqrt{2}} & 0 & 0 \end{bmatrix}, \quad \text{and} \quad \begin{bmatrix} 0 & 0 & 0 \\ 0 & 0 & -\frac{1}{\sqrt{2}} \\ 0 & \frac{1}{\sqrt{2}} & 0 \end{bmatrix}. \tag{3}$$

Notably, these vectors are orthonormal with respect to the Frobenius inner product. With this choice of metric and basis, the marginal distribution of a rotation matrix $r^{(t)}$ evolving according to Brownian motion for time $t$ from an initial rotation $r^{(0)}$ is given by the $\mathcal{IG}_{SO(3)}$ distribution [10, 11], which we write as $r^{(t)} \sim \mathcal{IG}_{SO(3)}(\mu = r^{(0)}, \sigma^2 = t)$. The density of the $\mathcal{IG}_{SO(3)}$ distribution with respect to the uniform distribution on $SO(3)$ is given by

$$\mathcal{IG}_{SO(3)}(r^{(t)}; \mu, \sigma^2) = f(\omega(\mu^\top r^{(t)}); \sigma^2), \text{ for } f(\omega; \sigma^2) = \sum_{l=0}^{\infty} (2l+1) e^{-l(l+1)\sigma^2} \frac{\sin((l+\frac{1}{2})\omega)}{\sin(\omega/2)}, \tag{4}$$

where $\mu$ is $3 \times 3$ mean rotation matrix and $\omega(r)$ denotes the angle of rotation in radians associated with a rotation $r$; in particular, we may compute $\omega(r)$ by expressing $r$ in an axis-angle parameterization, that is as a 3D vector pointing in the direction of a rotation axis, whose length reflects an angle of rotation about that axis. We approximate the power series in Equation (4) by its truncation after 2000 terms. We formulate a discrete-time forward noising by discretizing the Brownian motion, which provides: $q(r^{(t)} | r^{(t-1)}) = \mathcal{IG}_{SO(3)}(r^{(t)}; r^{(t-1)}, \sigma_t^2 - \sigma_{t-1}^2)$ and marginally $q(r^{(t)} \mid r^{(0)}) = \mathcal{IG}_{SO(3)}(r^{(t)}; r^{(0)}, \sigma_t^2)$, where $\sigma_t^2$ is a variance schedule for rotations. We choose $\sigma_t$ so that the rotations are corrupted at a rate similar to the forward process on translations (Table 6). In contrast to the translations, which converge to a Gaussian distribution as $t$ increases, the rotations converge to the uniform distribution on $SO(3)$.

**Backward transition kernel:** To approximate the reverse transitions for the rotations we take inspiration from De Bortoli et al. [8, Theorem 1] and approximate the reversal of this discretized reverse process by

$$p(r^{(t-1)} \mid x^{(t)}) = \mathcal{IG}_{SO(3)}(r^{(t-1)}; r^{(t)}\mathrm{exp}\left\{(\sigma_t^2 - \sigma_{t-1}^2) r^{(t)\top} \nabla_{r^{(t)}} \log q(x^{(t)})\right\}, \sigma_t^2 - \sigma_{t-1}^2), \quad (5)$$

where $\nabla_{r^{(t)}} \log q(x^{(t)})$ denotes the Stein score of the forward process at time $t$, and exp denotes the exponential map to $SO(3)$ from the Lie algebra of $SO(3)$; in the case $SO(3)$, exp is the matrix exponential (see Solà et al. [12] for an introduction and background on $SO(3)$, its Lie algebra, and their properties). Notably, $\nabla_{r^{(t)}} \log q(x^{(t)})$ is in the tangent space of $SO(3)$ at $r^{(t)}$, $r^{(t)\top} \nabla_{r^{(t)}} \log q(x^{(t)})$ is in the Lie algebra, and $r^{(t)}\mathrm{exp}\left\{(\sigma_t^2 - \sigma_{t-1}^2) r^{(t)\top} \nabla_{r^{(t)}} \log q(x^{(t)})\right\}$ is in $SO(3)$.

**Approximating the score with a denoising prediction:** Equation (5) describes how one could sample from the reverse process using the $\mathcal{IG}_{SO(3)}$ distribution based on the score of the forward process. One could in principle learn this score function directly by score matching training [8]. However, we instead rely on an approximation that directly leverages RoseTTAFold's ability to produce denoised structures. For a given $t$ and $r^{(t)}$ we may write

$$\begin{aligned}
\nabla_{r^{(t)}} \log q(x^{(t)}) &= \mathbb{E}_q \left[\nabla_{r^{(t)}} \log q(x^{(t)} \mid x^{(0)}) \mid x^{(t)}\right] \\
&= \mathbb{E}_q \left[\nabla_{r^{(t)}} \log q(r^{(t)} \mid r^{(0)}) \mid x^{(t)}\right] \\
&\approx \nabla_{r^{(t)}} \log q(r^{(t)} \mid r^{(0)} = \hat{r}^{(0)}) \\
&= \nabla_{r^{(t)}} \log \mathcal{IG}_{SO(3)}(r^{(t)}; \hat{r}^{(0)}, \sigma_t^2),
\end{aligned} \quad (6)$$

where the first line is known as the denoising score matching identity [13], the second line is obtained from the conditional independence structure of the forward process, the third line is an approximation that can be thought of as replacing $q(r^{(0)} \mid r^{(t)})$ with a point mass on the noiseless rotation $\hat{r}^{(0)}$ predicted by RF*diffusion*, and the final line recognizes the approximation as the gradient of the tractable $\mathcal{IG}_{SO(3)}$ log density. In the expressions above, we use the notation $\mathbb{E}_q[g(x^{(0)}, x^{(t)}) \mid x^{(t)}] = \int g(x^{(0)}, x^{(t)}) q(x^{(0)} \mid x^{(t)}) dx^{(0)}$ to describe the conditional expectation according to $q$ of $g(x^{(0)}, x^{(t)})$ given $x^{(t)}$. We compute the score approximation in the final line of Equation (6) by applying the chain rule to obtain

$$\begin{aligned} \nabla_r \log \mathcal{IG}_{SO(3)}(r; \hat{r}, \sigma_t^2) &= \nabla_r \omega(\hat{r}^\top r) \frac{d}{d\omega} \log f(\omega; \sigma_t^2) \mid_{\omega=\omega(\hat{r}^\top r)} \\ &= r \frac{\log(\hat{r}^\top r)}{\sqrt{2}\omega(\hat{r}^\top r)} \frac{d}{d\omega} \log f(\omega, \sigma_t^2)|_{\omega=\omega(\hat{r}^\top r)} \end{aligned} \tag{7}$$

where $\log(\hat{r}^\top r)$ is the logarithmic map from $SO(3)$ to the Lie algebra of $SO(3)$ (i.e. the matrix logarithm), and $\omega(r\hat{r}^\top)$ and $f$ are as defined in Equation (4). $r\frac{\log(\hat{r}^\top r)}{\sqrt{2}\omega(\hat{r}^\top r)}$ is a unit length perturbation in the direction of $\log(\hat{r}^\top r)$ applied in the tangent space of $SO(3)$ at $r$. And $\frac{d}{d\omega} \log f(\omega, \sigma_t^2) \mid_{\omega=\omega(\hat{r}^\top r)}$ is a scaling of this direction. In the above, the factor of $\frac{1}{\sqrt{2}}$ arises because we adopted the Frobenius scalar product as the inner product on the Lie algebra of $SO(3)$, and $\|r\frac{\log(\hat{r}^\top r)}{\sqrt{2}\omega(\hat{r}^\top r)}\|_F = 1$.

We reasoned that the approximation in Equation (6) may be reasonably accurate for two reasons. First, in the case of diffusion probabilistic models with Gaussian noise, where optimizing to convergence would provide $\hat{z}^{(0)}(z^{(t)}) = \mathbb{E}_q\left[z^{(0)} \mid x^{(t)}\right]$, this approximation holds exactly in the sense that $\mathbb{E}_q[\nabla_{z^{(t)}} \log q(z^{(t)} \mid z^{(0)}) \mid x^{(t)}] = \nabla_{z^{(t)}} q(z^{(t)} \mid z^{(0)} = z^{(0)}(x^{(t)}))$ (see Proposition 1 below). Though this does not hold with equality with $\mathcal{IG}_{SO(3)}$, because $SO(3)$ is a Riemannian manifold and is therefore locally Euclidean, $\mathcal{IG}_{SO(3)}$ closely resembles a Gaussian for small $t$. Second, again when $t$ is near 1, $x^{(t)}$ will be close to a plausible

**Algorithm 1** RF*diffusion* rotation score approximation

**function** $\mathrm{F}(\omega, \sigma^2, L = 2000)$ ▷ $\mathcal{IG}_{SO(3)}$ density factor, truncated to $L$ terms
**return** $\sum_{l=0}^{L}(2l+1)e^{-l(l+1)\sigma^2}\frac{\sin((l+\frac{1}{2})\omega)}{\sin(\omega/2)}$
**end function**

**function** ROTATIONSCOREAPPROXIMATION$(r_t, \hat{r}_0, \sigma_t^2)$
  $\vec{r}_{0t} = \log(\hat{r}_0^\top r_t)$ ▷ $\vec{r}_{0t} \in \mathbb{R}^{3,3}, \vec{r}_{0t} = -\vec{r}_{0t}^\top$
  $\omega_{0t} = \omega(\hat{r}_0^\top r_t)$ ▷ angle of rotation $\omega_{0t} \in [0, \pi]$

  ▷ Compute score approximation
  $s = \frac{r_t \vec{r}_{0t}}{\sqrt{2}\omega_{0t}} \cdot \frac{d}{d\omega} \log \mathrm{F}(\omega; \sigma_t^2)|_{\omega=\omega_{0t}}$ ▷ $s \in \mathbb{R}^{3,3}$
**return** $s$
**end function**

structure and, if the model is trained well, $q(r^{(0)} \mid x^{(t)})$ will be concentrated near $r^{(0)}$. Although approximation error may be non-trivial for larger $t$, we find this approximation to be empirically useful nonetheless.

In summary, we approximate reverse transitions by

$$p(r^{(t-1)} \mid x^{(t)}) = \mathcal{IG}_{SO(3)}(r^{(t-1)}; r^{(t)}\exp\{(\sigma_t^2 - \sigma_{t-1}^2)r^{(t)\top}\nabla_{r^{(t)}} \log \mathcal{IG}_{SO(3)}(r^{(t)}; \hat{r}^{(0)}, \sigma_t^2)\}, \sigma_t^2 - \sigma_{t-1}^2),$$

with $\nabla_{r^{(t)}}\mathcal{IG}_{SO(3)}(r^{(t)}; r^{(0)}, \sigma_t^2)$ computed as in Equation (5). We present computation of this approximation to the score in Algorithm 1.

**Proposition 1.** *Suppose $z^{(0)}, z^{(t)} \sim q(z^{(0)}, z^{(t)})$, and that $z^{(t)} \mid z^{(0)} \sim \mathcal{N}(z^{(t)}; \alpha z^{(0)}, \sigma^2)$ according to $q$ for some $\alpha$ and $\sigma^2$. If $\hat{z}^{(0)}(z^{(t)}) = \mathbb{E}_q[z^{(0)} \mid z^{(t)}]$ for every $z^{(t)}$, then*

$$\mathbb{E}_q[\nabla_{z^{(t)}} \log q(z^{(t)} \mid z^{(0)}) \mid z^{(0)}] = \nabla_{z^{(t)}} \log q(z^{(t)} \mid z^{(0)} = \hat{z}^{(0)}(z^{(t)})). \quad (8)$$

*Proof.* To prove the result, we re-write the left hand side of Equation (8) to obtain

$$\begin{aligned}
\mathbb{E}_q[\nabla_{z^{(t)}} \log q(z^{(t)} \mid z^{(0)})|z^{(t)}] &= \mathbb{E}_q[-(2\sigma^2)^{-1}(z^{(t)} - z^{(0)}) \mid z^{(t)}] \\
&= -(2\sigma^2)^{-1}(z^{(t)} - \mathbb{E}_q[z^{(0)} \mid z^{(t)}]) \\
&= -(2\sigma^2)^{-1}(z^{(t)} - \hat{z}^{(0)}) \\
&= \nabla_{z^{(t)}} \log q(z^{(t)} \mid z^{(0)} = \hat{z}^{(0)}(z^{(t)})),
\end{aligned} \tag{9}$$

as desired. □

#### 2.4 Self-conditioning in reverse process sampling

Self-conditioning was introduced by Chen et al. [14], who showed the technique dramatically improves text diffusion. We implement self-conditioning in the manner similar to how it is described by Chen et al. [14].

For sampling in diffusion generative models without self-conditioning, at each denoising step once $x^{(t)}$ has been sampled, the prediction of the denoised data from the previous step ($\hat{x}^{(0)}_{\text{prev.}} = \hat{x}^{(0)}(x^{(t+1)})$) is discarded. However, since each denoising step is typically small, the predictions $\hat{x}^{(0)}(x^{(t)})$ can be similar, so much of the denoising computation must be repeated. By contrast, with self-conditioning one saves the denoising predictions at each step and provides them as an input to the denoising model at the next iteration, instead predicting $x^{(0)}$ as $\hat{x}^{(0)}(x^{(t)}, \hat{x}^{(0)}_{\text{prev.}})$. This process of providing previous network outputs as an input to subsequent iterations is reminiscent of 'recycling' in AlphaFold [4] and our updated variant of RoseTTAFold (Section 1.2). When training with self-conditioning, on 50% of examples one performs a usual denoising step, setting $\hat{x}^{(0)}_{\text{prev.}} = 0$ and computing a loss as $L(x^{(0)}, \hat{x}^{(0)}_{\text{prev.}} = 0)$. The other 50% of the time, one (i) simulates an additional forward noising step to obtain $x^{(t+1)} \sim q(x^{(t+1)} \mid x^{(t)})$, (ii) computes $\hat{x}^{(0)}_{\text{prev.}} = \hat{x}^{(0)}(x^{(t+1)}, \hat{x}^{(0)}_{\text{prev.}} = 0)$,

**Algorithm 2** RF*diffusion* generation

**function** SampleReference(L)
  ▷ Random initial structure for $L$ residues
  **for** $l = 1, \ldots, L$ **do**
    $r_l^{(T)} \sim \text{Uniform}(SO(3))$
    $z_l^{(T)} \sim \mathcal{N}(0, I_3)$
    $x_l^{(T)} = (r_l^{(T)}, x_l^{(T)})$
  **end for**
**return** $x^{(T)}$
**end function**

**function** ReverseStep($x^{(t)}, \hat{x}^{(0)}$)
  ▷ One step of reverse diffusion
  **for** $l = 1, \ldots, L$ **do**
    $(r_l^{(t)}, z_l^{(t)}) = x_l^{(t)}$
    $(\hat{r}_l^{(0)}, \hat{z}_l^{(0)}) = \hat{x}_l^{(0)}$
    ▷ Update translations
    $z_l^{(t-1)} \sim \mathcal{N}(\frac{\sqrt{\bar{\alpha}^{(t-1)}}\beta^{(t)}}{1-\bar{\alpha}^{(t)}}\hat{z}_l^{(0)} + \frac{\sqrt{\alpha^{(t)}}(1-\bar{\alpha}^{(t-1)})}{1-\bar{\alpha}^{(t)}} z_l^{(t)}, \beta^{(t)})$

    ▷ Update rotations
    $s_l =$RotationScoreApproximation$(r_l^{(t)}, \hat{r}_l^{(0)}, \sigma_t^2)$
    $\Delta r_l = \exp\{(\sigma_t^2 - \sigma_{t-1}^2) r_l^{(t)\top} s_l\}$ ▷ exponential map from Lie algebra to SO(3)
    $r_l^{(t-1)} \sim \mathcal{IG}_{SO(3)}(r_l^{(t)}\Delta r_l, \sigma_t^2 - \sigma_{t-1}^2)$
    $x_l^{(t-1)} = (r_l^{(t-1)}, z_l^{(t-1)})$
  **end for**
**return** $x^{(t-1)}$
**end function**

**function** Sample(L)
  ▷ RF*diffusion* generation of $L$-residue backbone structure
  $x^{(T)} =$ SampleReference$(L)$
  $\hat{x}^{(0)}_{\text{prev.}} = \vec{0}$ ▷ Initialize self-conditioning
  **for** $t = T, \ldots, 1$ **do**
    $\hat{x}^{(0)} =$ RF*diffusion*$(x^{(t)})$
    $x^{(t-1)} =$ ReverseStep$(x^{(t)}, \hat{x}^{(0)})$
    $\hat{x}^{(0)}_{\text{prev.}} = \hat{x}^{(0)}$
  **end for**
**return** $\hat{x}^{(0)}$
**end function**

and (iii) computes a loss as $L(x^{(0)}, \hat{x}^{(0)}(x^{(t)}, \hat{x}^{(0)}_{\text{prev.}}))$, backpropagating gradients only through the second denoising step. Training and sampling with self-conditioning are described in Algorithms 2 and 3.

In RFdiffusion, we input $\hat{x}^{(0)}_{\text{prev.}}$ through the template structure feature (`xyz_t`, see Table 1) and we input $x^{(t)}$ as coordinates to the 3D track of RF (`xyz_prev`, see Table 1). Inputting $x^{(t)}$ as coordinates, as opposed to the distogram and anglegram used in the template structure feature, allows the network to keep the motif fixed in coordinate space.

#### 2.5 Mean squared error loss on residue frames

The primary objective used to train RF*diffusion* is mean squared error loss, averaged across training examples, time steps and the forward noising process,

$$\text{MSE}_{\text{Frame}} = \frac{1}{T}\sum_{t=1}^{T} \mathbb{E}_q[d_{\text{frame}}(x^{(0)}, \hat{x}^{(0)}(x^{(t)}))^2],$$

where

$$d_{\text{frame}}(x^{(0)}, \hat{x}^{(0)}) = \sqrt{\frac{1}{L}\sum_{l=1}^{L} \|z_l^{(0)} - \hat{z}_l^{(0)}\|_2^2 + \|I_3 - \hat{r}_l^{(0)\top} r_l^{(0)}\|_F^2},$$

is a metric on the sets of predicted frames consisting of Euclidean distance on the $C_\alpha$ coordinates ($\|z^{(0)} - \hat{z}^{(0)}\|_2$), and a metric on rotation matrices ($\|I_3 - \hat{r}^{0\top} r^{(0)}\|_F^2$, where $\|\cdot\|_F$ denotes the Frobenius norm [15]. In practice, we use slight modification of $\text{MSE}_{\text{Frame}}$ chosen to improve stability of training (see Section 4.1 for details).

In each training step, we compute an unbiased Monte Carlo estimate of this objective by sampling a time step $t \sim \mathcal{U}(1, \ldots, T)$, a single structure $x^{(0)}$ from our dataset, simulating the forward process to obtain $x^{(t)} \mid x^{(0)} \sim q(x^{(t)} \mid x^{(0)})$, and taking a gradient step on our slight modification of $\text{MSE}_{\text{Frame}}$. Algorithm 3 summarizes the training procedure.

Our choice of $\text{MSE}_{\text{Frame}}$ takes inspiration from [6]. In particular, Ho et al. [6, section

**Algorithm 3** RF*diffusion* Training

**function** FORWARDNOISE($x^{(0)}, t$)<br>
    **for** $l = 1, \ldots, L$ **do**<br>
        $(r_l^{(0)}, z_l^{(0)}) = x_l^{(0)}$<br>
        $z_l^{(t)} \sim \mathcal{N}\left(\sqrt{\bar{\alpha}^{(t)}} z_l^{(0)}, (1 - \bar{\alpha}^{(t)}) I_3\right)$<br>
        $r_l^{(t)} \sim \mathcal{IG}_{SO(3)}\left(r_l^{(0)}, \sigma_t^2\right)$<br>
        $x_l^{(t)} = (r_l^{(t)}, z_l^{(t)})$<br>
    **end for**<br>
**return** $x^{(t)}$<br>
**end function**<br>
<br>
**function** TRAIN<br>
    **while** not converged **do**<br>
        $x^{(0)} \sim$ TrainingSet<br>
        $t \sim$ Uniform($\{1, \ldots, T\}$)<br>
        **if** Uniform$(0, 1.0) < 0.5$ **or** $t = T$ **then**<br>
            ▷ Train step without self-conditioning<br>
            $x^{(t)} =$ ForwardNoise($x^{(0)}, t$)<br>
            $\hat{x}^{(0)}_{\text{prev.}} = \vec{0}$<br>
        **else**<br>
            ▷ Train step with self-conditioning<br>
            $x^{(t+1)} =$ ForwardNoise($x^{(0)}, t + 1$)     ▷ Sample $\left(x^{(t+1)}, x^{(t)}\right) \sim q(x^{(t:t+1)} \mid x^{(0)})$<br>
            $x^{(t)} =$ ReverseStep($x^{(t+1)}, x^{(0)}$)<br>
<br>
            ▷ Compute self-conditioning input<br>
            $\hat{x}^{(0)}_{\text{prev.}} =$ *RFdiffusion*($x^{(t+1)}, \vec{0}$)<br>
            $\hat{x}^{(0)}_{\text{prev.}} =$ StopGradient($\hat{x}^{(0)}_{\text{prev.}}$)<br>
        **end if**<br>
        $\hat{x}^{(0)} =$ *RFdiffusion*($x^{(t)}, \hat{x}^{(0)}_{\text{prev.}}$)<br>
        Take gradient step on $d_{\text{frame}}\left(x^{(0)}, \hat{x}^{(0)}\right)$<br>
    **end while**<br>
**end function**

3.2] comment that when the forward process consists of adding Gaussian noise, the training objective of minimizing the Kullback-Leibler divergence of $q(z^{(t-1)} \mid z^{(t)})$ to $p(z^{(t-1)} \mid z^{(t)})$ can be rewritten as a rescaling of the expected squared error of a prediction of $z^{(0)}$ from noisy observations $z^{(t)}$. In particular if we fix the variance of the backward transitions to $\beta_t$ as in Section 2.2, then for each $t$

$$\mathbb{E}_q[\mathrm{KL}(q(z^{(t-1)} \mid z^{(t)}) \| p(z^{(t-1)} \mid z^{(t)}))] = \mathbb{E}_q\left[\frac{\bar{\alpha}^{(t-1)}(1-\alpha_t)^2}{2\beta_t(1-\bar{\alpha}^{(t)})^2}\|z^{(0)} - \hat{z}^{(0)}(z^{(t)})\|_2^2\right] + c, \quad (10)$$

where $c$ is a constant that does not depend on $p$ (see [16, Equation 99] and [6]). Consequently when one minimizes the right-hand-side of Equation (10) for every $t$, they maximize a weighted variational lower bound on the likelihood of the data. Moreover, this bound is globally minimized only when each $p(z^{(t-1)} \mid z^{(t)})$ matches $q(z^{(t-1)} \mid z^{(t)})$, and $p(z^{(0)})$ therefore matches the data-distribution [6]. Although Ho et al. [6] found better performance in generative modeling of images when predicting the noise added in the forward process (rather than $x^{(0)}$), we reasoned that by predicting $x^{(0)}$ we could better leverage the inductive biases of RoseTTAFold pre-trained for structure prediction to produce realistic structures.

However, the equivalence of learning to optimally denoise according to average squared distance and matching the reverse process is only known to apply when the forward process consists of Gaussian noise, and likely does not hold for the $\mathcal{IG}_{SO(3)}$ noise used for rotations. However the squared Frobenius norm metric seemed to be a sensible choice because (1) our chosen forward noising process for rotations is approximately Gaussian in the tangent space of $SO(3)$ at $r^{(0)}$ for $t$ close to zero, and (2) this metric is approximately equal to a scaling of squared Euclidean distance in the tangent space of $SO(3)$ when $\hat{r}^{(0)}$ is close to $r^{(0)}$ [17].

#### 2.6 Geometric equivariance in RF*diffusion*

We build on previous work [18, 7] and seek to learn a distribution over protein structure that is invariant to rotation; that is, we require that any protein structure is modeled as equally likely upon a rigid rotation. More formally, this means that for any structure $x$ and rotation $R$ we desire to have that $p(x) = p(R*x)$, where $p$ denotes the density parameterized by the model, and $R*x = [R*x_1, \ldots, R*x_L] = [(Rr_1, Rz_1), \ldots, (Rr_N, Rz_L)]$ describes the structure obtained by rotating $x$ about the origin by $R$. Invariance to translation is also of interest in generative modeling of proteins but, because no normalized probability distribution can be invariant to translation, properly handling translational invariance requires contending with unnormalized measures and is beyond the scope of our discussion; in practice, we obviate this challenge by centering all training examples at or near the origin.

To enforce invariance of $p$ with respect to rotations in our DDPM, we follow prior work [19, 7] by (1) using a rotation invariant reference distribution (satisfying $p(x^{(T)}) = p(R * x^{(T)})$) and (2) constraining the reverse transitions to be equivariant to rotations, i.e. satisfying $p(x^{(t-1)}|x^{(t)}) = p(R * x^{(t-1)}|R * x^{(t)})$. Criterion (1) is readily satisfied by the choices of the the zero mean Gaussian with isotropic covariance as the reference distribution for translations, and the uniform distribution on $SO(3)$ for rotations. That criterion (2) is satisfied owes to the SE(3) equivariance of the denoising network inherited from RoseTTAFold. We may see this separately for translations and for rotations.

For translations, because RF's prediction $\hat{z}^{(0)}$ is rotationally equivariant with respect to $x^{(t)}$, and because $\hat{\mu}(x^{(t)})$ in Equation (2) is a linear combination of $\hat{z}^{(0)}$ and $z^{(t)}$, $\hat{\mu}(x^{(t)})$ and therefore also $p(z^{(t-1)} \mid z^{(t)})$ are equivariant with respect to $x^{(t)}$.

For rotations, when we compute $\nabla_{r^{(t)}} \log \mathcal{IG}_{SO(3)}(r^{(t)}; \hat{r}^{(0)}, \sigma_t^2)$ in Equation (5) from $r^{(t)}$ and the prediction $\hat{r}^{(0)}$ from RF*diffusion*, this score approximation is rotationally equivariant with respect to $x^{(t)}$. One may see that this is the case by observing that, because $\hat{r}^{(0)}$

is an equivariant prediction, $r\hat{r}^{(0)\top}$ is invariant to rotation. Consequently, $\log(r\hat{r}^{(0)\top})$ and $\omega(r\hat{r}^{(0)\top})$ are also invariant, and we can recognize the right hand size of Equation (5) as the product of $r^{(t)}$ and a second (invariant) matrix. Therefore $\nabla_{r^{(t)}} \log \mathcal{IG}_{SO(3)}(r^{(t)}; \hat{r}^{(0)}, \sigma_t^2)$ rotates equivariantly with $r^{(t)}$.

### 3 RF*diffusion*: methodology for controlled design

We next describe several techniques to *control* generation of backbones in RF*diffusion* to meet specific design criteria. Section 3.1 describes generation of symmetric oligomers. Section 3.2 describes our approach to training RF*diffusion* for generation via conditional training. Section 3.3 then describes how we modify the architecture of and fine-tune RF*diffusion* for targeted binder design, and design subject to topology constraints. Finally, Section 3.4 describes how we can guide generation with extrinsically defined "potentials".

#### 3.1 Generation of oligomers with point group symmetries

As discussed in the main text, generating oligomeric assemblies obeying desired point-group symmetry constraints is crucial in several design contexts. Point group symmetries may be represented by a finite collection of rotation matrices that form a mathematical group with respect to matrix multiplication as the group operation [20]. For example, we may represent the cyclic symmetry group of order $K$ by the set of rotation matrices that rotate increments of $(360/K)^o$ about the z-axis, $C_k = \{R_z^{(k/K)360^o}\}_{k=0}^{K-1}$. Analogous representations exist for all other point groups (including dihedral, octahedral, tetrahedral, and icosahedral). Without loss of generality we set the first rotation to be the identity $R_1 = I_3$. We represent an oligomer with $K$ monomer subunits each with $L$ residues by $X = [x^1, \ldots, x^K]$ where each subunit $k$ consists of the translations and rotations $x^k = ([z_1^k, \ldots, z_L^k], [r_1^k, \ldots, r_L^k])$. Then we say an oligomer obeys a point group symmetry $\mathcal{R} = \{R_1, \ldots, R_k\}$, if $X = [R_1 * x^1, \ldots, R_K * x^1]$ where $R * x_1 = ([R * z_1^k, \ldots, R * z_L^k], [R * r_1^k, \ldots, R * r_L^k])$ denotes the rotation of a monomer backbone structure by $R$.

Previous work has demonstrated some success generating designs with symmetry through Hallucination [21, 22] with the inclusion of penalty terms on the deviation of predicted structures from the desired symmetry, but this work suffered from large computational

**Algorithm 4** Generation of symmetric oligomers

**function** SampleSymmetric($M, \mathfrak{R} = \{R_k\}_{k=1}^{K}$)
  ▷ RFDiffusion generation of oligomer with symmetry $\mathfrak{R}$
  $x^{(T,1)} = \text{SampleReference}(M)$
  **for** $t = T, \ldots, 1$ **do**
    $X^{(t)} = [R_1 x^{(t,1)}, \ldots, R_K x^{(t,1)}]$ ▷ Symmetrize chains
    $\hat{X}^{(0)} = RF\,diffusion(X^{(t)})$
    $[x^{(t-1,1)}, \ldots, x^{(t-1,K)}] = \text{ReverseStep}(X^{(t)}, \hat{X}^{(0)})$
  **end for**
**return** $\hat{X}^{(0)}$
**end function**

cost (on the order of 1 GPU day per design) and low success rates, presumably due to the inability to precisely control the desired symmetry[22]. We hypothesized that RF*diffusion* by contrast could provide improved control over symmetries in design by maintaining symmetry in denoising predictions, and by allowing us to enforce hard symmetry constraints during the reverse process (Algorithm 4).

Although we do enforce exact symmetry through explicit symmetrization at each denoising step, we observed that RF*diffusion* provides predictions of the denoised oligomer structures that preserve the desired symmetry nearly exactly, even in the first denoising steps (Fig. S4A). This property of denoised predictions owes to the exact equivariance of RoseTTAFold with respect to global rotations and the approximate equivariance with respect to permutation (i.e. relabeling) of chains. In particular, Proposition 2 guarantees that rotation and permutation equivariance of a neural network are sufficient conditions for maintenance of point group symmetries in the neural network's output. In RF*diffusion*, exact rotation equivariance is inherited from the SE(3)-transformer architecture used in the structure module of RoseTTAFold [3]. Permutation equivariance by contrast arises if the intermediate representations and outputs for each residue are unaffected by the ordering of chains. This is nearly the case with RF*diffusion*, with the exception that the RoseTTAFold pair representation contains directional sequence distance feature inputs for each pair of residues, clipped between -32 and 32 residues away; since oligomers are presented to RoseTTAFold by incrementing the sequence position index at the start of each chain, the sign of these features breaks exact permutation equivariance. However, we find empirically that deviation from exact symmetry in RF*diffusion* predictions is minimal (Fig. S4A).

**Proposition on preservation of symmetry**: We next provide a proposition that more precisely illuminates the mechanism by which predictions of denoised structures maintain the desired symmetry at each step.

**Proposition 2.** *Consider any function* $F : [x_1, \dots, x_K] \to [y_1, \dots, y_K]$ *and point group symmetry* $\mathcal{R} = \{R_1, \dots, R_K\}$*. If* $F$ *is both*

1. *rotation equivariant, that is* $F([R * x_1, \dots, R * x_K]) = [R * y_1, \dots, R * y_K]$ *for every rotation matrix* $R$*, and*

2. *permutation equivariant, that is* $F([x_{\sigma(1)}, \dots, x_{\sigma(K)}]) = [y_{\sigma(1)}, \dots, y_{\sigma(K)}]$ *for every permutation* $\sigma$*,*

*then* $F$ *is symmetry preserving. In particular, for any* $x$*,* $F([R_1 * x, \dots, R_K * x]) = [R_1 * y, \dots, R_K * y]$ *for some* $y$*.*

Notably, Proposition 2 holds for any neural network satisfying assumptions on $F$ above. We now prove the proposition.

*Proof.* We first establish some basic properties about permutations of point groups. First note that every member $R_k \in \mathcal{R}$ defines a permutation of $\mathcal{R}$ since $\{R_k R_1, R_k R_2, \dots, R_k R_K\} = \mathcal{R}$. Let $\sigma_k$ denote the permutation associated with $R_1 R_k^T \in \mathcal{R}$. In particular, $\sigma_k$ is the permutation such that for each $m$, $R_{\sigma_k(m)} = (R_1 R_k^T) R_m$. Notably, $\sigma_k(k) = 1$ because $R_{\sigma_k(k)} = (R_1 R_k^T) R_k = R_1$. For any permutation $\sigma$, we let $\bar{\sigma}$ denote its inverse, the permutation such that $\bar{\sigma}(\sigma(k)) = k$ for every $k$. Lastly, note that for $R_{\bar{\sigma}_k(m)} = (R_k R_1^T) R_m$, and so $R_{\bar{\sigma}_k(1)} = (R_k R_1^T) R_1$.

Assume without loss of generality that $F([R_1 * x, \ldots, R_K * x])_1 = R_1 * y$. To prove the proposition, it suffices to show that for any $k$, $F([R_1 * x, \ldots, R_k * x])_k = R_k * y]) = R_k * y$. Consider $\sigma_k$ as defined above. We can write

$$\begin{aligned} F([R_1 * x, \ldots, R_K * x])_k &= F(R_{\bar{\sigma}_k(1)} x, \ldots, R_{\bar{\sigma}_k(K)} x)_{\sigma_k(k)} \\ &= F((R_k R_1^T) R_1 x, \ldots, (R_k R_1^T) R_K x)_1 \end{aligned}$$

where the first equality follows from permutation equivariance of $F$, and the second equality follows from the definitions of $\sigma_k$ and $\bar{\sigma}_k$. Finally, by the rotation equivariance of $F$,

$$\begin{aligned} F((R_k R_1^T) R_1 x, \ldots, (R_k R_1^T) R_K x)_1 &= (R_k R_1^T) F(R_1 x, \ldots, R_K x)_1 \\ &= (R_k R_1^T) R_1 y = R_k y. \end{aligned}$$

Therefore $F(R_1 x, \ldots, R_k x)_k = R_k y$ as desired. □

#### 3.2 Conditional training for functional-motif scaffolding

Our approach to scaffolding functional motifs with RF*diffusion* follows Trippe et al. [7], who treat motif-scaffolding as a conditional generative modeling problem. We partition the residues of a structure into the residues comprising the *motif* and those comprising the remainder of the backbone, which we refer to as the *scaffold* that supports it. For a structure with $L$ residues, we let $\mathcal{M}$ denote the (potentially discontiguous) set of indices corresponding to the motif and $\mathcal{S}$ be the remaining scaffold indices, such that the union of $\mathcal{M}$ and $\mathcal{S}$ is the set of indices up to $L$ (i.e. $M \cup S = \{1, \ldots, L\}$).

We write $x_{\mathcal{M}}$ to denote the structure of the motif residues and $x_{\mathcal{S}}$ to be the scaffold residue frames such that we may write the whole (un-noised) protein structure as $x^{(0)} =$

$[x_{\mathcal{M}}^{(0)}, x_{\mathcal{S}}^{(0)}]$. Our goal is to sample scaffold backbones from the conditional distribution $q(x_{\mathcal{S}}^{(0)} \mid x_{\mathcal{M}}^{(0)})$. To do this, we aim to learn the reversal of the forward noising process applied only to scaffold residues, with the motif held fixed, $p(x_{\mathcal{S}}^{(t-1)} \mid x_{\mathcal{S}}^{(t)}, x_{\mathcal{M}}^{(0)}) \approx q(x_{\mathcal{S}}^{(t-1)} \mid x_{\mathcal{S}}^{(t)}, x_{\mathcal{M}}^{(0)})$, where $q(x_{\mathcal{S}}^{(t-1)} \mid x_{\mathcal{S}}^{(t)}, x_{\mathcal{M}}^{(0)})$ is the conditional forward noising process described in Sections 2.2 and 2.3.

In earlier work, Wang et al. [23] demonstrated that RoseTTAFold may be trained to respect motif constraints provided as inputs through the template structure input features through retraining. Because the division of residues into motif and scaffold is specific to each design problem, we desired to train RF*diffusion* such it may be used for any location of the motif within the sequence. To this end, we took an amortized training approach, wherein for each motif-scaffolding training example we 1) begin with a structure $x^{(0)}$, 2) choose a random division into motif and scaffold $x^{(0)} = [x_{\mathcal{M}}^{(0)}, x_{\mathcal{S}}^{(0)}]$ (see Section 4.1 for details), 3) apply noise to the scaffold to obtain $x_{\mathcal{S}}^{(t)} \sim q(x_{\mathcal{S}}^{(t)} \mid x_{\mathcal{S}}^{(0)})$, and 4) compute a loss on the RF*diffusion* prediction $\hat{x}^{(0)}([x_{\mathcal{M}}^{(0)}, x_{\mathcal{S}}^{(t)}])$ of $x^{(0)} = [x_{\mathcal{M}}^{(0)}, x_{\mathcal{S}}^{(0)}]$. In order to encourage RF*diffusion* to not move the motif, we set the time-step input for motif residues to $t = 0$, we include both the motif and the scaffold residues when we compute the loss on the prediction.

Because motif sidechain geometry is crucial for most motif-scaffolding problems, we additionally provide the amino acid sequence and side chain torsion angles for motif residues as inputs to RF*diffusion* (provided through RoseTTAFold's template feature inputs). Overall this strategy is akin to the diffusion model inpainting training and generation described by Saharia et al. [24], who use randomly generated image masks.

In summary, generation of scaffolds conditional on a motif with RF*diffusion* differs from unconditional generation only in (1) the inclusion of noise-free motif backbone and sidechain structure in the template inputs and (2) replacement of the motif backbone coordinates in

$x^{(t)}$ with un-noised motif coordinates at each step and (3) setting of the timestep for motif residues to 0.

#### 3.3 Fine-tuning and architecture modifications

We wish to train a diffusion model which can condition on arbitrary features; to accomplish this, additional features must be input to RF*diffusion* beyond the features already taken by RoseTTAFold. Given the vast difference in performance between models trained from scratch versus those initialized from RF weights (Fig. S2F), we also wish to continue to initialize training from RF weights. To allow the addition of more features, we choose to expand the size of existing features in RF and to expand the corresponding size of the weights of the embedding layer which initially embeds the feature into the model. We initialize the weights associated with these newly added dimensions in these embedding layers to zero. With this initialization, the model gives exactly the same output as unmodified RF with the initial weights. Upon training, the model can then learn to use the newly expanded features.

In Section 4.3, we describe how we do this in detail for binder design, by controlling secondary structure adjacency and fold family of designs and directing generated binders to target hotspots. In addition to architectural modifications, RF*diffusion* can also be further fine-tuned on different conditional tasks; we describe how we have done this for improved scaffolding of functional motifs in Section 4.2.

#### 3.4 Guiding RF*diffusion* inference with external potentials

In addition to the network's ability to condition on structural motifs, the inference process can be guided by external potential functions to generate proteins which possess arbitrary desired properties, such as the existence of contacts with another protein or a desired surface concavity. Previous work has demonstrated that diffusion models can be made to sample from conditional distributions $p(x^{(0)} \mid y)$ without retraining if given a classifier able to operate on noisy samples, $p(y \mid x^{(t)})$ [9, Appendix I]. In particular, $p(y = 1 \mid x^{(t)})$ may be understood as a predicted probability that an example $x^{(0)}$ has a property of interest (or is in a given "class") given only the noised observation $x^{(t)}$. In contrast to unguided generation, wherein one noisily moves in the direction $\nabla_{x^{(t)}} \log p(x^{(t)})$ (which points toward $\hat{x}^{(0)}$), with guidance one instead follows $\nabla_{x^{(t)}} \log p(x^{(t)}) + \nabla_{x^{(t)}} \log p(y = 1 \mid x^{(t)})$ in the reverse step[9].

In the present work, we construct heuristic approximations of these classification log probabilities $P(x^{(t)}) \approx \log p(y = 1 \mid x^{(t)})$ for two protein conditional generation objectives, symmetric oligomer design (Fig. S6) and enzyme design with concave pockets (Fig. S7). We show how to incorporate them into the sampling procedure in Algorithm 5, and detail the functional forms of these potentials in Section 4.4. In this work, we consider potentials the are defined as a function of the $C_\alpha$ coordinates alone, and so (in each individual step) these potentials do not impact residue orientations.

**Algorithm 5** Generation with guidance

**function** SampleGuided(L, P, GuideScale)<br>
    ▷ Generation of $L$-residue backbone structure, guided by potential $P$<br>
    $x^{(T)} = \text{SampleReference}(M)$<br>
    **for** $t = T, \ldots, 1$ **do**<br>
        $\hat{x}^{(0)} = \text{RFDiffusion}(x^{(t)})$<br>
        $x^{(t-1)} = \text{ReverseStep}(x^{(t)})$<br>
        $x^{(t-1)} = x^{(t-1)} + \text{GuideScale}(t) \nabla_{x^{(t)}} P(x^{(t)})$ ▷ Apply guidance<br>
    **end for**<br>
**return** $\hat{x}^{(0)}$<br>
**end function**

#### 3.5 Tuning diversity by scaling noise at inference

For some problems, reported throughout the manuscript, we reduce the noise added at each step, by including a multiplicative factor to the variance of the noise. Typically, this improves design quality, at the expense of design diversity.

### 4 RF*diffusion*: training and fine-tuning

Sections 2 and 3 described key aspects of our formulation of RF*diffusion* and how we have approached using it to generate designs with desired properties. In this section, we provide precise details on RF*diffusion*. Section 4.1 details the inputs to and outputs of a "base" version of RF*diffusion*; these inputs and outputs are adapted from their uses in RoseTTAFold (Section 1.2). Section 4.1 also describes the precise losses used and other training information. Section 4.2 provides details of the variant of RF*diffusion* fine-tuned on an enzyme design task. Section 4.3 describes modifications to and funetuning of RF*diffusion* for design of protein-protein interactions. Finally, Section 4.4 describes two specific instances of guiding potentials. We subsequently present final details of how we applied RF*diffusion* and these fine-tuned variants to specific design tasks in Sections 5 and 6.

#### 4.1 RF*diffusion* base model

RF*diffusion* was trained on monomer structures in the PDB used for RoseTTAFold training. Training examples consist of the unconditional task 20% of the time and the motif-conditional task 80% of the time. For the motif-conditional task, a contiguous set of residues is selected as the motif, and the true sequence and structure are provided to the model. RF*diffusion* is trained starting from the final RF weights. RF*diffusion* does not use recycling.

**Losses:** RF*diffusion* was trained with a loss comprising two terms,

$$\mathcal{L}_{\text{Diffusion}} = \mathcal{L}_{\text{Frame}} + w_{\text{2D}}\mathcal{L}_{\text{2D}},$$

| Input name (Shape) | Description |
|---|---|
| msa_masked<br>(1,1, L, 48) | The truncated MSA now contains only the masked sequence (20aa, zeros (1), mask (1), repeat aa (20), zeros (1), repeat mask (1), zeros (2), N-term/C-term (2)) |
| msa_full<br>(1, 1, L, 25) | The full MSA now contains only the masked sequence (20aa, zeros (1), mask (1), zeros (1), N-term/C-term (2)) |
| seq<br>(1, L, 22) | The masked sequence (20aa, zeros (1), mask (1)) |
| xyz_prev<br>(L, 27, 3) | The coordinates of all atoms (N-Ca-C-O backbone (4), (up to) 10 sidechain atoms, (up to) 13 hydrogen atoms) |
| idx_pdb<br>(L) | The integer index of each residue. Used to assign each residue its neighboring residue. This feature has the same definition as in RF |
| t1d<br>(1, L, 22) | The one-dimensional features associated with $x^{(t)}$ (20 amino acids, mask (1), timestep (1)). The timestep is set to 1 for all fixed motif residues, and to $1 - \frac{t}{T}$ in all other positions. |
| t2d<br>(1, L, L, 44) | The two-dimensional features associated with $x^{(t)}$ structure. These features are computed from $x^{(t)}$, not from $\hat{x}^{(0)}_{\text{prev.}}$. (36 distance bins (2-20Å, 0.5Å bins) + 1 final distance bin (> 20Å), angle maps (sine and cosine of omega, theta and phi angle) (6), missing residue mask (1)) |
| xyz_t<br>(1, L, 27, 3) | The self-conditioning feature. As described in Section 2.4, this is $\hat{x}^{(0)}_{\text{prev.}}$. This feature is immediately converted to a distogram and anglegram representation by the model. (N, $C_\alpha$, C backbone atoms) |
| alpha_t<br>(1, L, 30) | The sidechain torsions of the motif region of $x^{(t)}$. For positions with a masked sequence, zeros are provided. Initially T, L, 10, 2, with sine and cosine of (omega, phi, psi angles (3), (up to) 4 torsion angles, $C_\beta$ bend (1), $C_\beta$ twist (1), $C_\gamma$ bend (1)). This is concatenated with a mask (T, L, 10, 1) indicating which torsion angles are present for a given amino acid, and reshaped to T, L, 30. |
| msa_prev<br>(1, L, Cm) | The MSA embedding recycling information. This is the model's previous embedding at each position in the masked sequence. Cm = 256 |
| pair_prev<br>(L, L, Cp) | The 2-D embedding recycling information. This is the model's previous embedding at each edge between each node. Cp = 128 |
| state_prev<br>(L, Cs) | The 1-D embedding recycling information. This is the model's previous embedding at each position in the query sequence. Cs = 16 |

Table 4: Description of features input to RF*diffusion*.

| Output name (Shape) | Description |
|---|---|
| msa<br>(1, L, Cm) | The model's final embedding at each position in the masked sequence. Cm = 256 |
| pair<br>(L, L, Cp) | The model's final embedding at each edge between each node. Cp = 128 |
| state<br>(L, Cs) | The model's final embedding at each position in the query sequence. Cs = 16 |
| xyz<br>(L, 27, 3) | The model's full-atom prediction of structure (N-Ca-C-O backbone (4), (up to) 10 sidechain atoms, (up to) 13 hydrogen atoms) |

Table 5: Outputs returned by RF*diffusion*.

where $\mathcal{L}_{\text{Frame}}$ is a modified variant of squared distance loss in $\text{MSE}_{\text{Frame}}$ (Section 2.5), $\mathcal{L}_{2\text{D}}$ is a distogram and anglegram loss, and $w_{2\text{D}}$ is a weighting factor. We now describe each loss.

$\mathcal{L}_{\text{Frame}}$ includes two modifications from $\text{MSE}_{\text{Frame}}$ intended to improve the stability of optimization. First, whereas $\text{MSE}_{\text{Frame}}$ relies on a distance computed simply as sum of squared distances defined on the translation and rotation components of residue frames, $\mathcal{L}_{\text{Frame}}$ relies on a weighted sum of these components that includes clamping on translation distance,

$$d_{\text{Frame}}(x^{(0)}, \hat{x}^{(0)}) = \sqrt{\frac{1}{L}\sum_{l=1}^{L}\left(w_{\text{trans}}\min(\|z_l^{(0)} - \hat{z}_l^{(0)}\|_2, d_{\text{clamp}})^2 + w_{\text{rot}}\|I_3 - \hat{r}_l^{(0)\top} r_l^{(0)}\|_F^2\right)},$$

where $w_{\text{trans}}$ and $w_{\text{rot}}$ are weights on the rotation and translation distances, and $d_{\text{clamp}}$ is a maximum distance above which translation distances are clamped. Second, $\mathcal{L}_{\text{Frame}}$ includes contributions from $d_{\text{Frame}}(x^{(0)}, \hat{x}^{(0)})$ computed at each intermediate structure module iteration with an exponential weighting, $\gamma$ that places greater importance on later outputs. In particular, we have

$$\mathcal{L}_{\text{Frame}} = \frac{1}{\sum_{i=0}^{I-1}\gamma^i}\sum_{i=1}^{I}\gamma^{I-i} d_{\text{Frame}}(x^{(0)}, \hat{x}^{(0),i})^2$$

where $\hat{x}^{(0),i}$ is the $i^{\text{th}}$ structure block output.

The second term in the loss, $\mathcal{L}_{\text{2D}}$, is inspired by trRosetta [25]. In contrast to the definitionally unimodal structure track outputs, the model outputs multimodal distributions of expected distances, dihedral angles, and planar angles between all pairs of contacting residues. $D_{:,l,l'}$, $\Omega_{:,l',l'}$, $\Phi_{:,l,l'}$, $\Theta_{:,l,l'}$, together describe the orientation of residue $l$ relative to residue $l'$. The following loss consists of the cross entropy between the one-hot histogram of the known inter-residue distances and orientations and the corresponding distributions predicted by the model.

$$\begin{aligned}
\mathcal{L}_{\text{2D}}(\text{logits}_d, \text{logits}_\omega, \text{logits}_\theta, \text{logits}_\phi, z_0) =& \text{CrossEntropy}(\text{logits}_{\text{dist}}, D) + \\
& \text{CrossEntropy}(\text{logits}_\omega, \Omega) + \\
& \text{CrossEntropy}(\text{logits}_\theta, \Theta) + \\
& \text{CrossEntropy}(\text{logits}_\phi, \Phi)
\end{aligned}$$

where:

$$\begin{aligned}
& D \in \mathbf{R}^{[\mathbf{C_{dist}xLxL}]}; D_{b,l,l'} = \mathbb{1}[\text{bin}_{D,b}^{low} \leq \max(\|C_{\beta,l'} - C_{\beta,l'}\|_2, 18.5) < \text{bin}_{D,b}^{high}] \\
& \Omega \in \mathbf{R}^{[\mathbf{C_{dist}xLxL}]}; \Omega_{b,l,l'} = \mathbb{1}[\text{bin}_{\Omega,b}^{low} \leq \text{Dihedral}(C_{\alpha,l}, C_{\beta,l}, C_{\alpha,l'}, C_{\beta,l'}) < \text{bin}_{\Omega,b}^{high}] \\
& \Theta \in \mathbf{R}^{[\mathbf{C_{dist}xLxL}]}; \Omega_{b,l,l'} = \mathbb{1}[\text{bin}_{\Theta,b}^{low} \leq \text{Dihedral}(N_{\alpha,l}, C_{\alpha,l}, C_{\beta,l}, C_{\beta,l'}) < \text{bin}_{\Theta,b}^{high}] \\
& \Phi \in \mathbf{R}^{[\mathbf{C_{phi}xLxL}]}; \Omega_{b,l,l'} = \mathbb{1}[\text{bin}_{\Phi,b}^{low} \leq \text{Planar}(C_{\alpha,l}, C_{\beta,l}, C_{\beta,l'}) < \text{bin}_{\Phi,b}^{high}]
\end{aligned}$$

and the bin edges for converting these angles and distances into a one-hot distribution are given by:

$$
\begin{aligned}
\text{bin}_{D,i} &= [\frac{i}{2}, \frac{i+1}{2}] \\
\text{bin}_{\Omega,i} = \text{bin}_{\Theta,i} &= [-\pi + \frac{2\pi i}{37}, -\pi + \frac{2\pi(i+1)}{37}] \\
\text{bin}_{\phi,i} &= [\frac{\pi i}{19}, \frac{\pi(i+1)}{19}].
\end{aligned}
$$

And the formulae for computation of dihedral and planar angles are given by

$$
\begin{aligned}
&\text{Dihedral}(a, b, c, d) = \text{atan2}( \\
&\qquad [c-b] \cdot (([b-a] \times [c-b]) \times ([c-b] \times [d-c])), \\
&\qquad \|c-b\|([b-a] \times [c-b]) \cdot ([c-b] \times [d-c])) \\
&\text{Planar}(a, b, c) = \arccos(\frac{(a-b) \cdot (c-b)}{\|a-b\|\|c-b\|}).
\end{aligned}
$$

**Motif-centering during training:** For both training and scaffold generation, we center the motif at the origin. In preliminary computational experiments, we found that a model trained with non-centered motifs extracted from structures that were globally centered exhibited biased motif placement. In particular, for motifs not placed at the origin we found that the scaffolds sampled from this model typically placed most residues towards the side of the motif oriented towards the origin. We interpreted this as subtle instance of undesirable label–leakage, and so trained RF*diffusion* with motifs centered at the origin to correct this.

Once the distance loss is computed on the motif, motif coordinates are detached from the computation graph. As such, predicted motif coordinates are treated as constants for the purpose of all other loss comptutations. While this does not affect the values of any losses, this choice ensures that the components of loss on which gradients are computed with respect to the predicted motif coordinates are minimized when by predictions which doe not move from its initial coordinates. We made this choice to prevent the possibility that other computed losses would drive the motif from its (desired) initialization, but have not thoroughly explored this choice empirically.

**Hyperparameters and coordinate scaling:** We train RF*diffusion* using the hyperparameters in Table 6. Although the coordinate inputs and outputs of RoseTTAFold are in units of Angstroms, we define the diffusion process in a downscaled space by dividing all coordinate values of $x^{(t)}$ and $x^{(0)}$ before performing each diffusion step by a factor of 4 (chosen empirically), and then scaling back up to Angstroms.

**Training time:** RF*diffusion* trained to convergence when initialized from RF weights in 5 epochs. This took 3 days on 8 NVIDIA A100 GPUs.

#### 4.2 Enzyme active site scaffolding by fine-tuning on minimal motifs

The version of RF*diffusion* fine-tuned for enzyme active site scaffolding is trained starting from the base version of RF*diffusion*. During fine-tuning 30% of tasks are from the base model task set (Table 6) and the other 70% are a "triple-contact" task, in which a random set of 3 residues all $>$ 10 residues apart in sequence space but with pairwise $C_\beta$–$C_\beta$ distances $<$ 6Å is selected to form a model "active site". These three residues are included in the motif, and for each, there is a 50% chance of including one flanking residue. If no such triad is found in the monomer (as is the case for approximately 23% of training PDBs), the task would fall back to the base model training task. In addition, the motif-specific

| **Parameter name** | **Value** |
|---|---|
| Crop size | 384 |
| Pseudo-batch size | 64 |
| $w_{\text{trans}}$ | 0.5 |
| $w_{\text{rot}}$ | 1.0 |
| $w_{\text{2D}}$ | 1.0 |
| $d_{\text{clamp}}$ | 10 |
| Structure block iteration decay rate $\gamma$ | 0.99 |
| Learning rate | 0.0005, No warm-up. Decay learning rate by 0.95 after every 10000 optimization steps. |
| Examples per epoch | 25600 |
| Number of diffusion timesteps (T) | 200 |
| Variance schedule for translations | $\beta^{(t)} = \beta^{(0)} + (\frac{t}{T})(\beta^{(T)} - \beta^{(0)})$ with $\beta^{(0)} = 0.01$ and $\beta^{(T)} = 0.07$. |
| Variance schedule for rotations | $\sigma_t = t * \beta_{min} + \frac{1}{2}(\frac{t}{T})^2(\beta_{max} - \beta_{min})$, with $\beta_{min} = 1.06$ and $\beta_{max} = 1.77$ |
| Fraction of protein residues masked (when motif is provided) | Randomly picked from a uniform distribution between 20% and 100%, inclusive. |
| Probability of motif being contiguous or discontiguous | 0.5 |
| Probability of providing self-conditioning information | 0.5 |
| Coordinate scaling | 0.25 |

Table 6: RF*diffusion* training hyperparameters.

displacement loss is upweighted by a factor of 10 to encourage the network to keep the motif fixed, in order to compensate for the fact that otherwise motif recapitulation would comprise a significantly lower portion of the overall loss due to the much shorter motif length in this task. The network was fine-tuned for 5 epochs in this manner.

#### 4.3 Architectural modifications for protein-protein interaction design

In this subsection, we describe RF*diffusion* architecture modifications to incorporate target "hot-spots" (Section 4.3.1) and desired binder topology (Section 4.3.2), and details fine-tuning training (Section 4.3.3) for protein-protein interaction (PPI) design.

##### 4.3.1 Protein–protein interface hotspots

When designing protein–protein interfaces, it is critical to be able to control the area of the target (fixed) protein to which the designed binding (diffused) protein should associate. To allow this control, we train the model to perform complex design conditioned on "interface hotspot" residues. We define an interface hotspot residue as any residue on the target (fixed) chain of the example that is within 10Å $C_\beta$–$C_\beta$ distance of the binder (diffused) chain.

The 1-D one-hot tensor of interface hotspot residues is concatenated to RF's 1-D template feature (t1d, Table 1). We train two models for complex design, one which includes just hotspot information and another that includes fold–conditioning information and hotspot information. The updated feature shapes and definitions of the entries in each dimension for each model are provided in Table 7, respectively.

| Input name (shape) | Description |
|---|---|
| `t1d`<br>(1, L, 24) | The one-dimensional features associated with $x^{(t)}$. A concatenation of one hot amino acids (20 features), mask (1 feature), timestep (1 feature), repeat mask (1 feature), hotspot (1 feature). |

Table 7: RF*diffusion* input feature modified for fine-tuning on complexes.

| Input name (shape) | Description |
|---|---|
| `t1d`<br>(1, L, 28) | The one-dimensional features associated with $x^{(t)}$ (20 amino acids, mask (1), timestep (1), repeat mask (1)[1], hotspot (1), secondary structure {'helix', 'sheet', 'loop', 'mask'} (4) ) |
| `t2d`<br>(1, L, L, 44) | The two-dimensional features associated with $x^{(t)}$ structure. These features are computed from $x^{(t)}$, not from $\hat{x}^{(0)}_{\text{prev.}}$. (36 distance bins (2–20Å, 0.5Å bins) + 1 final distance bin (> 20Å), angle maps (sine and cosine of omega, theta and phi angle) (6), missing residue mask (1), secondary structure {'adjacent', 'non-adjacent', 'mask'} (3)) |

Table 8: Modified features used in fine-tuning on complexes and fold-conditioning.

###### 4.3.2 Secondary structure and block adjacency

The idea to use secondary structure and block-adjacency information was first introduced by Anand and Achim [26]. We review the idea here, discuss the motivation behind the idea, and describe our implementation in depth.

Often, a protein designer will desire a protein with a specific fold (for example: a transmembrane pore made of a beta barrel or a protein binder made of a three-helix bundle). A protein fold is defined by (1) the secondary structure blocks (contiguous regions of alpha helix, beta strand, or loop) it contains and (2) the exact orientation (translation and rotation) of these blocks with respect to one another. A protein designer will often desire to generate diversity within a specific type of fold as well. In such cases, it is critical to allow the under-specification of the exact fold to allow for diversity within the generated structures; we call these broad collections of folds of the same type a "fold family".

We wish to train a model which can be conditioned on a fold family. To allow the under-specification of a fold we choose to provide the model coarse information on which secondary structure blocks are within a distance cutoff of one another. Specifically, we provide the following features to the model: (1) an [L,4] one-hot tensor where each position is assigned to a secondary structure type {helix, sheet, loop, mask} and (2) an [L,L] one-hot tensor (called the block adjacency matrix) where entries indicate membership in blocks that are within a distance cutoff of one another (as in Fig. S5B).

Secondary structure annotations of every residue in the training set were calculated using DSSP [27]. DSSP is a structure-based algorithm that assigns a per-residue classification of secondary structure type. The block-adjacency matrix of every structure in the training set was calculated from the secondary structure string returned by DSSP. Blocks are marked as "adjacent" in the block-adjacency matrix if (1) neither block is of loop type and (2) the minimum $C_\alpha$–$C_\alpha$ distance of any pair of inter-block residues is within 8Å.

The 1-D secondary structure tensor is concatenated to RF's 1-D template feature (`t1d`, see Table 1). The 2-D block-adjacency matrix is concatenated to RF's 2-D template feature (`t2d`, see Table 1). The updated feature shapes and the definitions of the entries in each dimension are provided in Tables 7 and 8.

###### 4.3.3 Fine-tuning for protein–protein interaction design

The version of RF*diffusion* fine-tuned on protein complexes, is trained starting from the base version of RF*diffusion* trained for 5 epochs. The training task consists of monomer examples (50%) and complex examples (50%). When the model is shown a complex example, only one side of the complex is noised, the other side is kept fixed (this is in keeping with established PPI design methods [28] where the target protein is kept fixed). When the model is shown a complex example the model is provided with the residue indices of 0–20% of the residues ("hotspot residues") in the interface on the fixed chain side (the interface is defined as all residues within 10Å $C_\beta$–$C_\beta$ distance of another chain), to permit targeting of the designed binder at inference time. In a separate model, also trained on protein complexes, during both complex and monomer training the model is provided with secondary structure 50% of the time and (independently) block-adjacency information 50% of the time for the noised region. The junctions between blocks of secondary structure and their corresponding entries in the block-adjacency matrix are masked during training, such that at inference time, one does not need to specify exact, per residue secondary structure and block-adjacency matrices. Specifically, 0–75% of secondary structure (and corresponding adjacency, when provided) is masked, with this masking occurring over junctions in secondary structure (mask length 1–8 residues).

#### 4.4 Potentials details for symmetric oligomers and pocket design

Section 3.4 describes our approach to guiding the reverse diffusion process with potentials. We now describe the details of our choices of $P(x^{(t)})$ in applications to symmetric oligomer design and design of enzymes with concave pockets. When designing symmetric oligomers, we employ an inter-chain and intra-chain contact potential to promote the formation of contacts between subunits. Letting $Z = [z^1, \ldots, z^K]$ denote the $C_\alpha$ coordinates in oligomer with $K$ subunits and $L$ residues in each subunit (so for each $k$, $z_k = [z_{k,1}, \ldots, z_{k,L}]$ with each $z_{k,l} \in \mathbb{R}^3$) we set

$$P_{\mathrm{sym}}(Z) = \sum_{1 \le k, k', \le L} \sum_{1 \le l, l' \le L} (\mathbb{1}[k \neq k'] w_{\mathrm{inter}} + \mathbb{1}[k = k'] w_{\mathrm{intra}}) \mathrm{Switch}(\|z_{k,l} - z_{k',l'}\|_2^2),$$

where $w_{\mathrm{inter}}$ and $w_{\mathrm{intra}}$ weight the inter-chain and intra-chain potentials, respectively. We set $w_{\mathrm{inter}} = 2$ and $w_{\mathrm{intra}} = 0.2$ to prioritize the formation of inter-subunit contacts while encouraging individual subunits to be well–packed.

$\text{Switch}(r) = \frac{1-(\frac{r-d_0}{rc})^n}{1-(\frac{r-d_0}{r_c})^m}$, is a switching function which smoothly transitions from 1 if two atoms are within contact range to 0 when they are out of range. We set the hyperparameters that control its functional dependence on distance as $n = 6$, $m = 12$, $d_0 = 8$, and $r_c = 4$, to reflect the contact distances we would expect between interacting sidechains. It is sufficient to bias the in the early sampling steps ($t \approx T$) to promote contacts in the higher order structure, and unnecessary to continue to do so at towards the end of design trajectories, by which point the quaternary structure is sufficiently determined. As such, we scale the potential by a "guide-scale", $g(t)$, as

$$P_{\text{sym}'}(Z, t) = g(t)P_{\text{sym}}(Z),$$

for $g(t) = (\frac{t}{T})^2$.

When designing enzymes, in addition to recapitulating the sidechain geometry of the active site, a pocket must be formed which has shape complementarity to the substrate. This condition can be captured effectively by a simple attractive-repulsive potential parameterized by the minimum distance between enzyme $C_\alpha$ carbons and substrate atoms. Denoting the coordinates of a substrate with $K$ atoms by $s = \{s_k\}_{k=1}^K$ and the $C_\alpha$ coordinates by $z = [z_1, \ldots, z_L]$, we set: $P_{\text{enzyme}}(z, s) = w_{\text{attr}}[\sum_{1 \leq l \leq L} \text{Switch}(\min_{1 \leq k \leq K} \|z_l - s_k\|_2^2) - w_{\text{rep}}[\sum_{1 \leq l \leq L} \text{Rep}(\min_{1 \leq k \leq K} \|z_l - s_k\|_2^2)]$, where $\text{Rep}(r) = \max(0, \frac{|r-r0|^p}{pr_0^{p-1}})$, and we set $w_{\text{attr}} = 1, w_{\text{rep}} = 4, r_c = 2$.

The gradient of $\text{Rep}(r)$ decays smoothly from $-1$ at $r = 0$ to 0 at $r = r_0$, penalizing clashes between the protein backbone and the substrate. We do not use a guide scale with $P_{\text{enzyme}}$, as the potential relates to fine-grained details of the structure which are not fully determined until late in the reverse diffusion process. Empirically, we find the model is sufficiently receptive and robust to bespoke potentials with hyperparameters chosen based on physical intuition. We find that were able to achieve our objectives of interface production in the case of symmetric oligomer design, and implicit substrate modeling in the case of enzyme design without exhaustive hyperparameter tuning.

### 5 In silico experimental methods

#### 5.1 Unconditional benchmarking

To test RF*diffusion* on unconditional generation of monomers (Fig. 1C-F), we generated 100 designs for lengths 70, 100, 200, 300, 400, 600, 800 and 1000 amino acids. For each backbone, we generated 8 sequences with ProteinMPNN and subsequently predicted their structures with AF2 (or ESMFold - Fig. S1I). The best sequence (by alignment of the predicted structure to the design model) was taken for each backbone. We benchmarked against the recently-published RoseTTAFold Hallucination [21]. As some knowledge of how best to use RoseTTAFold for Hallucination is required, these samples were generated by the respective expert. ProteinMPNN was used to design sequences for all benchmarking designs. For ProteinMPNN, a sampling temperature of 0.1 was used, and cysteines were omitted from the designs (as these are often problematic for protein purification). Fourteen 300 amino acid proteins and four 200 amino proteins were ordered and tested experimentally for expression and CD profiles.

#### 5.2 Conditional benchmarking

The full conditional benchmark is described in Supplementary Table 1, and encompasses 25 design challenges from six recent publications [23, 7, 29, 30, 31, 32]. RF*diffusion* was compared to RoseTTAFold Hallucination and $RF_{joint}$, Inpainting. While both Hallucination and Inpainting are able to generate sequences directly, for the fairest comparison, we also redesigned the sequence with ProteinMPNN, and took the best of 8 sequences per backbone. Both $RF_{joint}$ Inpainting and RF Hallucination are able to scaffold structure without sequence, so in cases where functional-site residues were not required for function, these methods were permitted to redesign the sequence of the non-functional residues, which is generally beneficial for design. Finally, as Hallucination requires some expert knowledge and empirical hyperparameter tuning, some exploration of the benchmark set was permitted, and these designs were generated by the respective expert.

For a number of comparisons made in the paper (Fig. S1, S2C-D), a smaller benchmark encompassing a subset of unconditional and conditional benchmark problems described above was used.

| Name [Ref.] | Description | Input | Total Length | Sequence to be redesigned* |
|---|---|---|---|---|
| 1PRW[23] | Double EF-hand motif | 5-20,A16-35,10-25,A52-71,5-20 | 60-105 | A16-19,A21,A23,A25, A27-30,A32-35,A52-55, A57,A59,A61,A63-66,A68-71 |
| 1BCF[23] | Di-iron binding motif | 8-15,A92-99,16-30,A123-130,16-30,A47-54,16-30,A18-25,8-15 | 96-152 | A19-25,A47-50,A52-53,A92-93,A95-99,A123-126,A128-129 |
| 5TPN[23] | RSV F-protein Site V | 10-40,A163-181,10-40 | 50-75 | A163-168,A170-171,A179,A189 |
| 5IUS[23] | PD-L1 binding interface on PD-1 | 0-30,A119-140,15-40,A63-82, 0-30 | 57-142 | A63,A65,A67,A69,A71,A72,A76, A79,A80,A82,A119,A120,A121, A122,A123,A125,A127,A129, A130,A131,A133,A135,A137, A138,A140 |
| 3IXT[31] | RSV F-protein Site II | 10-40,P254-277,10-40 | 50-75 | P255,P258-259,P262-263, P268,P271-272,P275-276 |
| 5YUI[23] | Carbonic anhydrase active site | 5-30,A93-97,5-20,A118-120,10-35,A198-200,10-30 | 50-100 | A93,A95,A97,A118,A120 |
| 1QJG[23] | Delta5-3-ketosteroid isomerase active site | 10-20,A38,15-30,A14,15-30,A99,10-20 | 53-103 | n/a |
| 1YCR[23] | P53 helix that binds to Mdm2 | 10-40,B19-27,10-40 | 40-100 | B17-18,B20-22,B24-25 |
| 2KL8[23, 29] | De novo designed protein | A1-7,20,A28-79 | 79 | n/a |
| 7MRX_60[29] | Barnase ribonuclease inhibitor | 0-38,B25-46,0-38 | 60 | n/a |
| 7MRX_85[29] | Barnase ribonuclease inhibitor | 0-63,B25-46,0-63 | 85 | n/a |
| 7MRX_128[29] | Barnase ribonuclease inhibitor | 0-122,B25-46,0-122 | 128 | n/a |
| 4JHW[30] | RSV F-protein Site 0 | 10-25,F196-212,15-30,F63-69, 10-25 | 60-90 | F196,F198,F203,F211-212,F63,F69 |
| 4ZYP[30] | RSV F-protein Site 4 | 10-40,A422-436,10-40 | 30-50 | A422-427,A430-431,A433-436 |
| 5WN9[31] | RSV G-protein 2D10 site | 10-40,A170-189,10-40 | 35-50 | A170-175,A188-189 |
| 6VW1[23, 32] | ACE2 interface binding SARS-CoV-2 | E400-510/20-30,A24-42,4-10, A64-82,0-5† | 62-83 | A25-26,A29-30,A32-34,A36-42,A64-82 |
| 5TRV_short[7] | De novo designed protein | 0-35,A45-65,0-35 | 56 | n/a |
| 5TRV_med[7] | De novo designed protein | 0-65,A45-65,0-65 | 86 | n/a |
| 5TRV_long[7] | De novo designed protein | 0-95,A45-65,0-95 | 116 | n/a |
| 6E6R_short[7] | Ferridoxin Protein | 0-35,A23-35,0-35 | 48 | n/a |
| 6E6R_med[7] | Ferridoxin Protein | 0-65,A23-35,0-65 | 78 | n/a |
| 6E6R_long[7] | Ferridoxin Protein | 0-95,A23-35,0-95 | 108 | n/a |
| 6EXZ_short[7] | RNA export factor | 0-35,A28-42,0-35 | 50 | n/a |
| 6EXZ_med[7] | RNA export factor | 0-65,A28-42,0-65 | 80 | n/a |
| 6EXZ_long[7] | RNA export factor | 0-95,A28-42,0-95 | 110 | n/a |

Table 9: **A benchmarking set of recently published functional-motif scaffolding problems.** To benchmark RF*diffusion* at functional-site scaffolding, against existing methods, we generated a benchmark set encompassing problems described in six recent publications [23, 7, 29, 30, 31, 32], which utilize a range of design methodologies to address these problems. For each problem, named by PDB accession (and, where applicable, the length of the designs to be generated, left column), we recapitulated the inputs as closely as possible with respect to details available in each publication. So that others can test methods on this benchmark, the exact input is specified in the third column. In bold, prefixed by a letter, are the inputs (chain, residues) from the PDB structure provided to the model (the "functional-site"). In non-bold text are the lengths that the different methods randomly sampled to generate good designs. The final lengths of the proteins were either specified by the input to the model, or were provided as constraints (for example, for '6EXZ_Long', the model could sample any N- and C-terminal length between 0 and 95 residues, but the total length of the output had to equal 110 amino acids). For each design challenge, 100 designs were generated, and, where ProteinMPNN was used, 8 sequences were designed, with the best sequence chosen for each backbone. *Both the $RF_{joint}$ and RoseTTAFold constrained hallucination approaches can simultaneously redesign sequences during generation, which can, in some cases, be helpful (if extracting the motif exposes hydrophobic residues which may subsequently end up as surface residues in the output designs, for example). Therefore, in this benchmark, these methods were allowed to redesign non-functional residues, listed in the right-most column. † This example is multi-chain generation (scaffolding a functional-site in the presence of a second chain). All methods benchmarked here can represent chain breaks (with large residue index jumps). Full results are shown in Fig. 3A, and tabulated in Table 10.

#### 5.3 Assessing diversity of designs

Designs were assessed for their diversity both to each other, and to the PDB (PDB100 April 19, 2022), using the TM score [33]. In Fig. S1F, designs were clustered at a 0.6 pairwise TM score cutoff.

#### 5.4 Assessing choice of losses

Previous work on using DDPMs for protein design has used Frame Aligned Point Error (FAPE) as the loss function [26]. FAPE was introduced in AF2 and was also used to train RoseTTAFold. FAPE is SE(3) invariant but not invariant to reflections, this makes it an ideal loss for protein structure prediction where the exact global orientation of the

| **Problem Name** | **RFdiffusion (noise=0)** | **RFdiffusion (noise=1)** | **RFjoint** | **RFjoint + Protein-MPNN** | **RF Hallucination** | **RF Hallucination + Protein-MPNN** |
|---|---|---|---|---|---|---|
| 1BCF | **100** | 98 | 65 | 94 | 0 | 0 |
| 6E6R_med | **89** | 67 | 0 | 14 | 2 | 9 |
| 2KL8 | 88 | **96** | 71 | 62 | 20 | 34 |
| 6E6R_long | **86** | 63 | 0 | 1 | 0 | 1 |
| 6EXZ_long | **76** | 51 | 0 | 0 | 1 | 4 |
| 1YCR | **74** | 58 | 12 | 20 | 11 | 61 |
| 6VW1 | **69** | 66 | 0 | 2 | 2 | 32 |
| 5TPN | **61** | 59 | 0 | 1 | 0 | 1 |
| 6EXZ_med | **49** | 33 | 0 | 0 | 5 | 15 |
| 4ZYP | **40** | 31 | 1 | 7 | 1 | 18 |
| 6E6R_short | **39** | 29 | 0 | 15 | 3 | 7 |
| 5TRV_long | **37** | 30 | 0 | 0 | 0 | 2 |
| 3IXT | 25 | 16 | 21 | 24 | 2 | **34** |
| 5TRV_med | **24** | 20 | 0 | 0 | 0 | 3 |
| 7MRX_85 | **11** | 6 | 0 | 0 | 0 | 0 |
| 7MRX_128 | **9** | 4 | 0 | 0 | 0 | 0 |
| 1PRW | 8 | **9** | 0 | 5 | 0 | 0 |
| 5TRV_short | 4 | **7** | 0 | 0 | 0 | 1 |
| 7MRX_60 | **2** | 0 | 0 | 0 | 0 | 0 |
| 6EXZ_short | 2 | 4 | 1 | 10 | 4 | **15** |
| 5IUS | **2** | 0 | 0 | 0 | 0 | 0 |
| 5YUI | 0 | 0 | 0 | 0 | 0 | 0 |
| 5WN9 | 0 | **1** | 0 | 0 | 0 | 0 |
| 4JHW | 0 | 0 | 0 | 0 | 0 | 0 |
| 1QJG | 0 | **1** | 0 | 0 | 0 | 0 |

Table 10: **Functional-site scaffolding benchmark results.** Full results for the benchmark test described in Fig. 3A and Table 1. In each case, values represent the success rate (%) in a set of 100 designs generated with each method.

predicted structure is arbitrary, but chirality within the structure is important. With a DDPM, however, $x^{(0)}$ must be in the same global frame as $x^{(t)}$ since $x^{(0)}$ and $x^{(t)}$ are interpolated between to generate $x^{(t-1)}$. We reasoned that, as FAPE is SE(3) invariant, a model trained with FAPE would not learn to make predictions in the same global frame as the inputs. We tested this by comparing a model trained with FAPE to a model trained with the $C_\alpha$ and rotation squared distance losses described in Section 1.3. By contrast these losses are not SE(3) invariant.

We found that the model trained with FAPE was unable to perform unconditional generation; the model was unable to preserve a global frame and this caused the denoising process to diverge after repeatedly interpolating between coordinates in different frames. In the motif scaffolding task, $x^{(0)}$ and $x^{(t)}$ can be aligned to one another using the fixed motif, this effectively eliminates the global frame problem as any arbitrary SE(3) action applied by the model can be reversed by this motif-alignment step. In the motif-scaffolding task we found that the model trained with displacement losses dramatically outperformed the model trained with FAPE (Fig S1A). We attribute the performance difference to better agreement of the squared distance losses with the objective of matching the reversal of the forward process, which thereby yields a model that better matches the data distribution (Section 2.2, 2.3, 2.5).

#### 5.5 Design of fold-conditioned proteins

To design TIM barrels, we constructed secondary structure and block adjacency inputs from a previously-designed TIM barrel (PDB: 6WVS). Any regions of loop secondary structure were masked, and to generate larger TIM barrels than the original, we randomly sampled additional length (1-15 residues inserted as "mask" tokens into the loops). We generated a total of 2400 designs, generating designs with three different noise scales during inference

(0, 0.5, 1). No external potentials were used. Designs were classified as TIM barrels if the TM score against 6WVS was greater than 0.5, and AF2 repredicted the designs (pAE $< 5$, RMSD to design $< 2$Å). 11 designs passing stringent filters (AF2 pAE ¡ 3.5, RMSD AF2 vs design $< 0.75$Å) were ordered.

To design NTF2 folds, we constructed secondary structure and block adjacency inputs from a preexisting set of 1000 NTF2 proteins. These were randomly selected and used as input to RFdiffusion. 900 designs were generated in total, at three different noise scales (0, 0.5, 1). Designs were classified as NTF2 folds if the TM score to PDB: 1GYS.

#### 5.6 Design of symmetric oligomers

To better understand RF*diffusion*'s capacity for designing symmetric oligomers, we generated backbones for the following groups: dihedral (D2, D3, D4, D5), cyclic (C3, C5, C6, C8, C10, C12), tetrahedral, octahedral, and icosahedral. We tested RF*diffusion*'s ability to design symmetry for these groups both with and without a guiding potential function for inter- and intra- chain contacts, weighting in all cases the intrachain contacts over the interchain. For dihedral, cyclic, and tetrahedral symmetries, protomers had 60-110 AA per chain, and for a subset of the cyclic symmetries (C3, C5, C6), additional models were designed with large protomers (150-400 AA per chain) to test RF*diffusion*'s ability to design unconditional yet large oligomers. The octahedral and icosahedral models were designed by modeling the minimal number of subunits (100-200 AA per protomer) required to capture all axes of symmetry (O: 4-, 3-, and 2-fold; I: 5-, 3-, and 2-fold).

Original backbones were filtered by sufficient oligomeric interfaces (determined by $C_\alpha$-$C_\alpha$ backbone distances between chains) to enrich for backbones with a higher likelihood for assembly following design. Cyclic and D2 symmetries were filtered for backbones consisting of protomers forming at least two distinct 10 residue interfaces, whereas all other symmetries were required to form at least three distinct 10 residue interfaces. Following filtering, all backbones were redesigned with ProteinMPNN, and then sequences were validated by AF2 (for the cyclic and dihedral symmetries). Given the complexity and challenge these symmetries present, we provided AF2 with an initial guess, as done in Bennett et al. [28], and increased the number of recycles the model could use in the predictions. Tetrahedra were predicted using RoseTTAFold, and octahedron and icosahedron were predicted with AF2 along their C3 axes of symmetry only. Designs were considered successful (success rates for cyclic and dihedral shown in Fig. S4) if the structure predictions had a mean pLDDT $> 80$ and an RMSD between prediction and design model of $< 2$Å. This same filtering regime was also used for the cage symmetries, but applied to the C3 predictions (for octahedra and icosahedra), and the monomer predictions (for tetrahedra).

#### 5.7 Design of p53 helix scaffolds

To design scaffolds able to hold the Mdm2-binding helix of p53, we used the version of RF*diffusion* fine-tuned for protein-protein interaction design (Section 4.2), and provided the network with both the p53 helix and the whole Mdm2 protein structure from PDB: 1YCR. To encourage extra contacts with the target protein, we used an external potential to encourage inter-chain contacts (Section 4.4). No fold information was provided to the network in this case.

#### 5.8 Design of theoretical C3-symmetric spike SARS-CoV-2 spike protein binding oligomers

To design the theoretical C3-symmetric oligomers to scaffold the ACE2 mimic AHB2, we started with the C3-symmetric cryo-EM structure of the minibinders against the spike protein from [34]. The first 55 residues of the minibinder were used as the asymmetric unit in a C3-symmetric motif input to the model.

The model weights from both the 5th epoch of RF*diffusion* training as well as the 8th epoch were used with T=200 length trajectories. All combinations of inter- and intrachain contact potentials with weights (0.1,0.3,0.5,1) and (0.5,1), respectively, were applied to the trajectories, with 25 designs being computed per combination. 32 ProteinMPNN sequences per design were computed, and AF2 was used to predict the structure of the oligomers via the inference method described by Wicky et al. [22].

#### 5.9 Design of symmetric $Ni^{2+}$ binding oligomers

To design the C4-symmetric $Ni^{2+}$-binding proteins, we started from three sets of imidazole groups positioned in square planar coordination geometry bearing C4 rotational symmetry with the associated symmetry axis being aligned to the Z-axis (Fig 4B,S8A). The imidazoles were placed with the NE2 atoms at a distance of 2.2 Å from the metal center (a common bond length for His–$Ni^{2+}$ in the MetalPDB [35]) and the different sets of imidazole groups were positioned such that they formed dihedral angles of 0°, 22°, and 45° between the Z-axis and the plane of the heterocyclic system (Figure S8, A). We note that larger dihedral angles resulted in clashing imidazole moieties and were therefore not considered in our designs.

Next, three sets of backbone dependent, non-clashing inverse rotamers [36] from the Dunbrack rotamer library were sampled for pieces of ideal alpha-helix ($\phi = -40°, \psi = -60°$) containing the histidine rotamers in the middle, and an alanine residue on either side of the histidine (three residues total per asymmetric unit going into the model). For set 1, rotamers chosen were of probability 0.3502, 0.1207, 0.0647, 0.0474, 0.0469 (Figure S8, B), for set 2 probability 0.0365, and for set 3 probabilities 0.3502, 0.0648, 0.0305, and 0.0131 (Figure S8, E). Note that the differences between the sets is that set 1 is associated with scaffolding the imidazole groups with no shear (0° dihedral) while sets 2 and 3 are associated with scaffolding the imidazole groups with shear (22°, 45°).

After construction of the inverse rotamers, the imidazole groups from their histidines were aligned to the aforementioned square-planar imidazole groups, resulting in various C4-symmetric motifs that could then be input to the model. 100 reverse diffusion trajectories were run for the full T=200 steps for all symmetric motifs, with 50 residues designed on either side of the inverse rotamer helix chunks in each chain (total complex length 412 residues, coordinating histidine always at position 52 in each chain). As in Section 4.4, an intra-chain guiding potential was used during the trajectory with a weight of 1, an inter-chain guiding potential with a weight of .06, and a global multiplicative factor of 2. For set 1, half (50) of the designs per motif were designed such that the effect of the external potential decayed quadratically during the trajectory, while the other half having potentials decay cubicly, while for sets 2 and 3 only a quadratically decaying external potential was utilized. Importantly, multiple models from the training session which produced RF*diffusion* were tested to see which checkpoint could scaffold the sites most accurately, and pilot experiments suggested the set of weights after the 8th epoch, rather than the 5th epoch (standard used for this paper) should be used.

Before sequence design with ProteinMPNN, RF*diffusion* outputs were filtered to only allow designs for which the backbone RMSD from the model < 1Å RMSD from the true motif. This yielded 199 backbones for set 1 and 201 backbones for sets 2 and 3, and ProteinMPNN was then used to perform symmetric sequence design on all residues except the histidines (the original alanines in the motif were also re-designed), with 16 sequences per backbone. AF2 was then used to predict the structure of all designed sequences.

To assemble a final set of 24 designs to order for testing from set 1, designs from the set were filtered with the following criteria: (1) full-atom RMSD on all 4 histidines between the AF2 prediction and the motif $\leq 0.6$Å, (2) AF2 pLDDT $\geq 90$ (3) AF2 PAE $\leq 6$. From here, set 1 designs were clustered at a TM-score cutoff of 0.85, yielding 20 representative backbones. 4 additional designs passing the RMSD, pLDDT and PAE metrics (presumably with TM-score $> 0.85$ vs some of the aforementioned 20) were hand-picked to create a final 24 designs from set 1.

To assemble a final set of 24 designs to order for testing from sets 2 and 3, designs from these sets were filtered with the following criteria: (1) Full atom RMSD on all 4 histidines between the AF2 prediction and the motif $\leq 0.66$Å, (2) AF2 pLDDT $\geq 90$, (3) AF2 PAE $\leq 6$. This yielded exactly 24 designs without clustering by TM-score.

Mutant sequences for all 48 ordered designs were created by simply replacing the histidine at position 52 with alanine. 44 of the designs were successfully transformed into *E. coli.*, which is what is reported on in the main text and the experimental methods.

#### 5.10 Design of protein binders to rigid targets

To test the ability of RF*diffusion* to design de novo binders to rigid targets, we designed binders to five targets: PD-L1 (PDB: 5O45), IL7 Receptor $\alpha$ (PDB: 3DI3), Insulin Receptor (PDB: 4ZXB), TrkA Receptor (PDB: 1WW7) and Flu Hemagglutinin (PDB: 5VLI). We generated designs both with and without fold conditioning, with the folds used derived from scaffold sets typically used for Rosetta-based protein binder design [37]. In all cases, we targeted binders, using input "hotspot" residues, to a specific site on the target protein. In line with current best practice [28], we tried using Rosetta FastRelax[38] before running a single ProteinMPNN, although we found that this was not systematically helpful for design success rates. For the five design cases, we generated several thousand designs. We classed a design as successful if it had AF2 pAE of interaction between binder and target $< 10$ (this has been shown to be highly indicative of design success), as well as RMSD between the designed binder and the AF2 prediction < 1Å, and AF2 pLDDT > 80. Success rates are reported in Fig. 4B, and were several orders of magnitude higher than with traditional Rosetta binder design.

#### 5.11 Figures and statistics shown in the paper

Protein structures depicted in this paper were rendered in PyMOL [39], and graphs were plotted with Matplotlib [40] and Seaborn [41]. Note that for all boxplots displayed in the paper, for aesthetic reasons, outliers are not displayed. Appropriate statistical tests were performed using SciPy [42], as indicated in figure legends.

### 6 In vitro experimental methods

#### 6.1 Plasmid construction

Symmetric oligomer, unconditional proteins, TIM barrels, COVID binder trimer scaffolds, and protein binder designs were ordered as synthetic genes (eBlocks, Integrated DNA Technologies) with compatible BsaI overhangs to the target cloning vector, LM0627 (see Wicky et al. [22]) for Golden Gate assembly. LM0627 is a modified expression vector containing a Kanamycin resistance gene and a ccdb lethal gene between BsaI cut sites. Subcloning into LM0627 results in the following product: MSG-[**protein**]-GSGSHHWGSTHHHHHH, with the C-terminal SNAC [43] cleavage tag and 6XHis affinity tag respectively underlined. Helical peptide binders were ordered in a similar format, except for the addition of adaptors (GGGSGGGGSASHMRS, SSEISFCSEPPPSRRS) permitting cloning into the pETcon3 vector (as well as LM0627), to permit both purification in E. coli and yeast surface display.

#### 6.2 Protein expression and purification

For the oligomeric, unconditional proteins, TIM barrels, COVID binder trimer scaffolds, and protein binder expression screens, a previously reported protocol was followed[22], with some modifications as denoted. In short, Golden Gate subcloning reactions of designs were carried out in 96-well PCR plates in 1$\mu$L volume. Reaction mixtures were then transformed into a chemically competent expression strain (BL21(DE3)), and 1-hour outgrowths were split directly into four 96-deep well plates containing 0.9-1.0mL of auto-induction media (autoclaved TBII media supplemented with Kanamycin, 2mM MgSO4, 1X 5052) for a final total volume of approximately 4mL. The following day (20-24 hrs later), cells were harvested and lysed, and clarified lysates were applied directly to a 50$\mu$L bed of Ni-NTA agarose resin in a 96-well fritted plate equilibrated with a Tris wash buffer. After sample application and flow through, the resin was thoroughly washed, and samples were eluted in 200$\mu$L of a Tris elution buffer containing 300mM imidazole. For oligomers, 0.5 M EDTA was spiked into the eluates (10 mM final) to reduce self-association due to the 6XHis tag. All eluates were sterile filtered with a 96-well 0.22$\mu$m filter plate (Agilent 203940-100) prior to size exclusion chromatography.

Protein designs were then screened via SEC using an AKTA FPLC outfitted with an autosampler capable of running samples from a 96-well source plate. The symmetric oligomers, unconditional proteins, TIM barrels, and COVID binder trimer scaffolds were run on a Superdex200 Increase 5/150 GL column (Cytiva 28990945). The protein binders were run on a Superdex75 Increase 5/150 GL column (Cytiva 29148722). The icosahedral designs were run on a Superose6 5/150 GL column (Cytiva 29091597). For the cyclic and dihedral symmetric oligomers, and the unconditional proteins, TIM barrels, and COVID binder trimer scaffolds, either a running buffer of 20 mM NaPhos pH 7.4, 100 mM NaCl or 20 mM Tris pH 8, 50 mM NaCl was used. For the tetrahedral, octahedral, and icosahedral oligomers, samples were run in 20 mM Tris pH 8, 50 mM NaCl, 100 mM Glycine. To improve peak resolution, the SEC column was connected directly in line from the autosampler to the UV detector. 0.25 mL fractions were collected from each run, and selected fractions were pooled for further analysis (LC-MS, native mass spectrometry, negative stain EM, SDS-page).

Genes encoding the designs for Ni2+-binding were cloned into a C-terminal Strep-tag construct via the Golden Gate method. Resulting plasmids were transformed into BL21(DE3) cells and protein expression was performed at 50 mL scale via autoinduction for approximately 24 hours, in which the first 4 hours cultures were grown at 37°C and the remaining time at 18°C. Cultures were harvested at 4000xg for 10 minutes in a tabletop centrifuge, supernatant discarded, and resuspended in approximately 30 mL lysis buffer (50 mM Tris-

HCl, 150 mM NaCl, 1 mM EDTA, 0.1 mg/mL lysozyme, 0.01 mg/mL DNAse, $\frac{1}{2}$ tablet of pierce protease inhibitor tablet/50 mL culture, pH 8.0). Sonication was performed with a 4-prong head for 5 minutes total, 10s pulse on-off at 80% amplitude. The resulting lysate was clarified by centrifugation at 14000xg for 30 minutes. Resulting supernatant was applied to 1 mL of Streptactin resin equilibrated with wash buffer (50 mM Tris-HCl, 150 mM NaCl, 1 mM EDTA, pH 8.0) and incubated for approximately 15 minutes with mild agitation. Resin was subsequently washed with at least 10 CVs of wash buffer and 0.4 CVs of elution buffer (50 mM Tris-HCl, 150 mM NaCl, 1 mM EDTA, 2.5 mM desthiobiotin, pH 8.0). Another 1.3 CV of elution buffer was applied to the resin and eluate was collected for purification by size-exclusion chromatography. Samples were applied to an S200 column equilibrated once with 20 mM HEPES, 50 mM NaCl, 1 mM EDTA, pH 7.4 to ensure removal of any trace metals, then again with the same buffer without EDTA.

#### 6.3 Negative-Stain EM sample preparation

De novo designed oligomeric proteins were diluted to $\sim$ 0.1mg/mL for negative stain. 3$\mu$L of the diluted complexes were immediately negatively stained after diluting using Gilder Grids overlaid with a thin layer of carbon and 0.75% uranyl formate.

#### 6.4 Negative-Stain EM data collection, processing, and validation

Data were collected on an Talos L120C 120kV electron microscope equipped with a CETA camera. A total of $\sim$150-250 images were collected per sample by using a random defocus range of 1.3–2.3 $\mu$m, with a total exposure of between 30 and 50 e-/A2, with a pixel size of either 1.54 or 2.49 Å/pixel. All data were automatically acquired using EPU (ThermoFisher Scientific). All data processing was performed using CryoSPARC V4.0.3 [44].

The parameters of the contrast transfer function (CTF) were estimated using Patch CTF, with minimal and maximal fitting resolutions set to 40Å and 8Å, respectively. Particles were picked initially in a reference-free manner using blob picker, followed by template picking using well-defined 2D classes of intact oligomers. Particles were extracted after correcting for the effect of the CTF for each micrograph with a box size of 80 pixels, except for icosahedron HE0902, which was extracted with a box size of 180 pixels to account for its large relative size. Extracted particles were sorted by reference-free 2D classification over 20 iterations. Given the small size of these particles, 2D classification was performed both in the presence and absence of CTF correction, with the best resulting classes selected for 3D *ab initio* reconstruction. More often than not, turning off CTF correction dramatically improved 2D class average quality. Notably, only constructs that displayed a combination of both "top-down" and "side" views (to ensure complete angular coverage) were selected for nsEM 3D reconstruction attempts. 3D *ab initio* jobs for each RFdiffusion construct displaying good angular coverage were performed by sorting into 3-4 classes, in a single attempt, in the presence and absence of the appropriate symmetry operator and compared. Resulting *ab initio* jobs which immediately converged on a map that exhibited clearly discernible features bearing a striking degree of similarity to both the 2D class average projections and computational design model were subsequently rigid-body docked against the AlphaFold2 prediction model for validation. All *ab initio* 3D maps with near perfect agreement to the design model were next run through homogenous refinement in the absence of applied symmetry to further validate their authenticity. Any 3D maps where any level of ambiguity was observed were immediately discarded. Furthermore, any 3D reconstruction attempts requiring multiple rounds of *ab initio* generation to yield convergence on a map in agreement with the design model were deemed as "low confidence" and were also discarded. For both cases, only the 2D classification results were reported in the supplementary material.

#### 6.5 CryoEM Sample Preparation and Data Collection

CryoEM grids were prepared by diluting protein samples with TBS 1 to 10 times immediately before applying 3.5 $\mu$L to glow-discharged 400 mesh, C-flat, 2 micron holes, 2 micron spacing, CF-2/2-4C (CF-224C-100) (Electron Microscopy Sciences) cryoEM grids. For D4 samples, 6 consecutive blots were applied in order to obtain the highest particle density [45]. Grids were blotted using a blot force of 0 and 5.5 second blot time at 100% humidity and 4°C and plunge-frozen in liquid ethane using a Vitrobot Mark IV (FEI Thermo Scientific). cryoEM grids were screened on a Glacios transmission electron microscope (FEI Thermo Scientific) operated at 200 kV and equipped with a K3 Summit direct detector. Automated Glacios data collection was carried out using SerialEM software at a nominal magnification of 36,000x (0.883 Å/pixel). Movies were acquired in counting mode fractionated in 50 frames of 200 ms at 8.5 e-/pixel/sec for a total dose of $\sim$ 65e-/Å$^2$. Details of data processing for each design are illustrated in Figure S13.

#### 6.6 CryoEM data processing

Multiple datasets were collected for each design and combined early on during processing. See Fig. S13 and processing flowcharts for details. Briefly, images were manually curated to remove poor quality acquisitions such as bad ice or large regions of carbon. Dose-weighting and image alignment of all 50 frames was carried out using MotionCor2 [46] with 5X5 patch or with cryosparc v4 patch alignment tool with default parameters. Super-resolution data was binned 2X during alignment. Initial CTF parameters were estimated using CTFfind4 [47]. Particle picking was done with a Gaussian blob picker and in some cases followed by a template picker. Particles were extensively classified in 2D to remove ice and noisy particles, but unfortunately yielded practically no "top" or "tilted" view particles. However, multiple orthogonal side views down the 2-fold axis were observed displaying high agreement to the design model. Starting models for all designs were always obtained *ab initio*. FSC curves were generated using cryoSPARC. All EM maps will be deposited in the EMDB and can be found in the supplementary data.

#### 6.7 Circular dichroism experiments

For circular dichroism (CD) experiments, designs (TIM barrels or unconditional designs) were diluted to 0.2mg/ml in 20mM Tris (pH 8.0) and 50mM NaCl. Spectra were acquired on a JASCO J-1500 CD Spectrophotometer. Thermal melt analyses were performed between 25° and 95°C, measuring CD at 222 nm. All reported measurements were acquired within the linear range of the instrument.

#### 6.8 Bio-layer Inferometry (BLI) Binding Experiments

BLI experiments were performed on an Octet Red96 (ForteBio) instrument, with streptavidin coated tips (Sartorius Item no. 18-5019). Buffer comprised 1X HBS-EP+ buffer (Cytiva BR100669) supplemented with 0.1% w/v bovine serum albumin. Prior to target loading, each design was tested for binding against unloaded tips via a 120 s baseline, 120 s association and 120 s dissociation cycle. For IL7-Ra, PD-L1, Mdm2, hemagglutinin and TrkA, 40nM of biotinylated target protein was loaded on the tips for 300 s followed by a 60 s baseline measurement. After loading, all designs underwent a 120 s baseline, 120 s association and 120 s dissociation. For each design, a previously validated minibinder[37] was included alongside 95 new designs. Three out of four positive controls were the same sequence as the previously reported designs with the addition of an MSG on the N-terminus with SNAC and His tags on the C-terminus, as per the cloning protocol described. The flu positive control was a reengineered version of the previously reported HA binder (PDB: 7RDH) where non-interface residues were redesigned using ProteinMPNN to improve expression. Baseline measurements of unloaded tips were subtracted from their matched measurement of the loaded tip. The response was taken as the average reading from 105 - 115 s during association. Binders were classified as those whose response was $>$ 50% of the control, indicating an approximate Kd of $<$ 10$\mu$L given comparable molecular weights. The positive control for InsR showed a saturating signal of only 0.08 nm. As such, we could not confidently separate the signal of 50% Rmax for our designs from baseline drift. Alternative approaches to experimental validation for this binder set is underway and will be reported on shortly. Up to 20 of the hits were taken forward for further titration experiments where concentration, association and dissociation times were chosen based on apparent affinity from the single point screen. Global kinetic fitting was used to determine Kds across the dilution series [48].

#### 6.9 Comparison of experimental success rates between design campaigns

All 5 targets chosen for *de novo* binder design were previously targeted for binder design with earlier Rosetta-based methods. The success rates in Fig 5C under "Rosetta" come from the following publications: IL-7R$\alpha$, and TrkA are from Cao et al. [37] in Extended Data Table 1; PD-L1 is from an unpublished binder campaign using the method of Cao et al. [37]. For PD-L1 a binder is called a success if the upper bound KD estimate calculated from yeast surface display enrichment is less than 50 $\mu$L; Flu HA is from Fleishman et al. [49], where a success rate of 2 / 88 is reported.

#### 6.10 Isothermal Titration Calorimetry

Protein and NiSO4 samples were prepared in buffer containing 20 mM HEPES, 50 mM NaCl, pH 7.4. Protein sample concentrations ranged from 30 $\mu$M to 100 $\mu$M and NiSO4 samples were prepared at 10 times the effective concentration of Ni2+ coordination sites (e.g. 100 $\mu$M of a designed tetramer yields 25 $\mu$M Ni2+ coordination sites in the sample cell, and thus a NiSO4 concentration of 250 $\mu$M was used in the syringe). Isothermal titration calorimetry experiments were performed on an automated Microcal PEAQ-ITC. Fits of the resulting data for the determination of dissociation constants (KD) was performed in the Microcal PEAQ-ITC Analysis Software.

### References

[1] Minkyung Baek, Frank DiMaio, Ivan Anishchenko, Justas Dauparas, Sergey Ovchinnikov, Gyu Rie Lee, Jue Wang, Qian Cong, Lisa N. Kinch, R. Dustin Schaeffer, Claudia Millán, Hahnbeom Park, Carson Adams, Caleb R. Glassman, Andy DeGiovanni, Jose H. Pereira, Andria V. Rodrigues, Alberdina A. van Dijk, Ana C. Ebrecht, Diederik J. Opperman, Theo Sagmeister, Christoph Buhlheller, Tea Pavkov-Keller, Manoj K. Rathinaswamy, Udit Dalwadi, Calvin K. Yip, John E. Burke, K. Christopher Garcia, Nick V. Grishin, Paul D. Adams, Randy J. Read, and David Baker. Accurate prediction of protein structures and interactions using a three-track neural network. *Science (New York, N.Y.)*, 373(6557):871–876, August 2021. ISSN 1095-9203 0036-8075. doi: 10.1126/science.abj8754.

[2] Nathaniel Thomas, Tess Smidt, Steven Kearnes, Lusann Yang, Li Li, Kai Kohlhoff, and Patrick Riley. Tensor field networks: Rotation- and translation-equivariant neural

networks for 3D point clouds, May 2018. URL http://arxiv.org/abs/1802.08219. arXiv:1802.08219 [cs].

[3] Fabian B. Fuchs, Daniel E. Worrall, Volker Fischer, and Max Welling. SE(3)-Transformers: 3D Roto-Translation Equivariant Attention Networks, November 2020. URL http://arxiv.org/abs/2006.10503. arXiv:2006.10503 [cs, stat].

[4] John Jumper, Richard Evans, Alexander Pritzel, Tim Green, Michael Figurnov, Olaf Ronneberger, Kathryn Tunyasuvunakool, Russ Bates, Augustin Žídek, Anna Potapenko, Alex Bridgland, Clemens Meyer, Simon A. A. Kohl, Andrew J. Ballard, Andrew Cowie, Bernardino Romera-Paredes, Stanislav Nikolov, Rishub Jain, Jonas Adler, Trevor Back, Stig Petersen, David Reiman, Ellen Clancy, Michal Zielinski, Martin Steinegger, Michalina Pacholska, Tamas Berghammer, Sebastian Bodenstein, David Silver, Oriol Vinyals, Andrew W. Senior, Koray Kavukcuoglu, Pushmeet Kohli, and Demis Hassabis. Highly accurate protein structure prediction with AlphaFold. *Nature*, 596(7873):583–589, August 2021. ISSN 1476-4687 0028-0836. doi: 10.1038/s41586-021-03819-2.

[5] Jascha Sohl-Dickstein, Eric A. Weiss, Niru Maheswaranathan, and Surya Ganguli. Deep Unsupervised Learning using Nonequilibrium Thermodynamics, November 2015. URL http://arxiv.org/abs/1503.03585. arXiv:1503.03585 [cond-mat, q-bio, stat].

[6] Jonathan Ho, Ajay Jain, and Pieter Abbeel. Denoising Diffusion Probabilistic Models, December 2020. URL http://arxiv.org/abs/2006.11239. arXiv:2006.11239 [cs, stat].

[7] Brian L Trippe, Jason Yim, Doug Tischer, Tamara Broderick, David Baker, Regina Barzilay, and Tommi Jaakkola. Diffusion probabilistic modeling of protein backbones in 3d for the motif-scaffolding problem. *arXiv preprint arXiv:2206.04119*, 2022.

[8] Valentin De Bortoli, Emile Mathieu, Michael Hutchinson, James Thornton, Yee Whye Teh, and Arnaud Doucet. Riemannian Score-Based Generative Modelling, November 2022. URL http://arxiv.org/abs/2202.02763. arXiv:2202.02763 [cs, math, stat].

[9] Yang Song, Jascha Sohl-Dickstein, Diederik P. Kingma, Abhishek Kumar, Stefano Ermon, and Ben Poole. Score-Based Generative Modeling through Stochastic Differential Equations, February 2021. URL http://arxiv.org/abs/2011.13456. arXiv:2011.13456 [cs, stat].

[10] Dmitry I Nikolayev and Tatjana I Savyolov. Normal distribution on the rotation group so (3). *Textures and Microstructures*, 29, 1970.

[11] Adam Leach, Sebastian M Schmon, Matteo T Degiacomi, and Chris G Willcocks. Denoising diffusion probabilistic models on so (3) for rotational alignment. In *ICLR 2022 Workshop on Geometrical and Topological Representation Learning*, 2022.

[12] Joan Solà, Jeremie Deray, and Dinesh Atchuthan. A micro Lie theory for state estimation in robotics, December 2021. URL http://arxiv.org/abs/1812.01537. arXiv:1812.01537 [cs].

[13] Pascal Vincent. A Connection Between Score Matching and Denoising Autoencoders. *Neural Computation*, 23(7):1661–1674, July 2011. ISSN 0899-7667, 1530-888X. doi: 10.1162/NECO_a_00142.

[14] Ting Chen, Ruixiang Zhang, and Geoffrey Hinton. Analog Bits: Generating Discrete Data using Diffusion Models with Self-Conditioning, August 2022. URL http://arxiv.org/abs/2208.04202. arXiv:2208.04202 [cs].

[15] Pierre M. Larochelle, Andrew P. Murray, and Jorge Angeles. A Distance Metric for Finite Sets of Rigid-Body Displacements via the Polar Decomposition.

*Journal of Mechanical Design*, 129(8):883–886, August 2007. ISSN 1050-0472, 1528-9001. doi: 10.1115/1.2735640. URL `https://asmedigitalcollection.asme.org/mechanicaldesign/article/129/8/883/471695/A-Distance-Metric-for-Finite-Sets-of-RigidBody`.

[16] Calvin Luo. Understanding diffusion models: A unified perspective. *arXiv preprint arXiv:2208.11970*, 2022.

[17] Du Q. Huynh. Metrics for 3D Rotations: Comparison and Analysis. *Journal of Mathematical Imaging and Vision*, 35(2):155–164, October 2009. ISSN 1573-7683. doi: 10.1007/s10851-009-0161-2. URL `https://doi.org/10.1007/s10851-009-0161-2`.

[18] Namrata Anand and Possu Huang. Generative modeling for protein structures. In *Advances in Neural Information Processing Systems*, volume 31. Curran Associates, Inc., 2018. URL `https://proceedings.neurips.cc/paper/2018/hash/afa299a4d1d8c52e75dd8a24c3ce534f-Abstract.html`.

[19] Minkai Xu, Lantao Yu, Yang Song, Chence Shi, Stefano Ermon, and Jian Tang. GeoDiff: a Geometric Diffusion Model for Molecular Conformation Generation, March 2022. URL `http://arxiv.org/abs/2203.02923`. arXiv:2203.02923 [cs, q-bio].

[20] D. S. Goodsell and A. J. Olson. Structural symmetry and protein function. *Annual Review of Biophysics and Biomolecular Structure*, 29:105–153, 2000. ISSN 1056-8700. doi: 10.1146/annurev.biophys.29.1.105.

[21] Ivan Anishchenko, Samuel J. Pellock, Tamuka M. Chidyausiku, Theresa A. Ramelot, Sergey Ovchinnikov, Jingzhou Hao, Khushboo Bafna, Christoffer Norn, Alex Kang, Asim K. Bera, Frank DiMaio, Lauren Carter, Cameron M. Chow, Gaetano T. Montelione, and David Baker. De novo protein design by deep network hallucination. *Nature*,

600(7889):547–552, December 2021. ISSN 1476-4687. doi: 10.1038/s41586-021-04184-w. URL https://www.nature.com/articles/s41586-021-04184-w. Number: 7889 Publisher: Nature Publishing Group.

[30] Fabian Sesterhenn, Che Yang, Jaume Bonet, Johannes T. Cramer, Xiaolin Wen, Yimeng Wang, Chi-I. Chiang, Luciano A. Abriata, Iga Kucharska, Giacomo Castoro, Sabrina S. Vollers, Marie Galloux, Elie Dheilly, Stéphane Rosset, Patricia Corthésy, Sandrine Georgeon, Mélanie Villard, Charles-Adrien Richard, Delphyne Descamps, Teresa Delgado, Elisa Oricchio, Marie-Anne Rameix-Welti, Vicente Más, Sean Ervin, Jean-François Eléouët, Sabine Riffault, John T. Bates, Jean-Philippe Julien, Yuxing Li, Theodore Jardetzky, Thomas Krey, and Bruno E. Correia. De novo protein de-

sign enables the precise induction of RSV-neutralizing antibodies. *Science (New York, N.Y.)*, 368(6492), May 2020. ISSN 1095-9203 0036-8075. doi: 10.1126/science.aay5051.

[31] Che Yang, Fabian Sesterhenn, Jaume Bonet, Eva A. van Aalen, Leo Scheller, Luciano A. Abriata, Johannes T. Cramer, Xiaolin Wen, Stéphane Rosset, Sandrine Georgeon, Theodore Jardetzky, Thomas Krey, Martin Fussenegger, Maarten Merkx, and Bruno E. Correia. Bottom-up de novo design of functional proteins with complex structural features. *Nature chemical biology*, 17(4):492–500, April 2021. ISSN 1552-4469 1552-4450. doi: 10.1038/s41589-020-00699-x. Place: United States.

[32] Anum Glasgow, Jeff Glasgow, Daniel Limonta, Paige Solomon, Irene Lui, Yang Zhang, Matthew A. Nix, Nicholas J. Rettko, Shoshana Zha, Rachel Yamin, Kevin Kao, Oren S. Rosenberg, Jeffrey V. Ravetch, Arun P. Wiita, Kevin K. Leung, Shion A. Lim, Xin X. Zhou, Tom C. Hobman, Tanja Kortemme, and James A. Wells. Engineered ACE2 receptor traps potently neutralize SARS-CoV-2. *Proceedings of the National Academy of Sciences of the United States of America*, 117(45):28046–28055, November 2020. ISSN 0027-8424. doi: 10.1073/pnas.2016093117. URL `https://www.ncbi.nlm.nih.gov/pmc/articles/PMC7668070/`.

[33] Yang Zhang and Jeffrey Skolnick. Scoring function for automated assessment of protein structure template quality. *Proteins*, 57(4):702–710, December 2004. ISSN 1097-0134 0887-3585. doi: 10.1002/prot.20264. Place: United States.

[34] Andrew C. Hunt, James Brett Case, Young-Jun Park, Longxing Cao, Kejia Wu, Alexandra C. Walls, Zhuoming Liu, John E. Bowen, Hsien-Wei Yeh, Shally Saini, Louisa Helms, Yan Ting Zhao, Tien-Ying Hsiang, Tyler N. Starr, Inna Goreshnik, Lisa Kozodoy, Lauren Carter, Rashmi Ravichandran, Lydia B. Green, Wadim L. Matochko, Christy A. Thomson, Bastian Vögeli, Antje Krüger, Laura A. VanBlargan,

Rita E. Chen, Baoling Ying, Adam L. Bailey, Natasha M. Kafai, Scott E. Boyken, Ajasja Ljubetič, Natasha Edman, George Ueda, Cameron M. Chow, Max Johnson, Amin Addetia, Mary Jane Navarro, Nuttada Panpradist, Michael Gale, Benjamin S. Freedman, Jesse D. Bloom, Hannele Ruohola-Baker, Sean P. J. Whelan, Lance Stewart, Michael S. Diamond, David Veesler, Michael C. Jewett, and David Baker. Multivalent designed proteins neutralize SARS-CoV-2 variants of concern and confer protection against infection in mice. *Science translational medicine*, 14(646): eabn1252, May 2022. ISSN 1946-6234. doi: 10.1126/scitranslmed.abn1252. URL `https://www.ncbi.nlm.nih.gov/pmc/articles/PMC9258422/`.

[35] Claudia Andreini, Gabriele Cavallaro, Serena Lorenzini, and Antonio Rosato. MetalPDB: a database of metal sites in biological macromolecular structures. *Nucleic Acids Research*, 41(D1):D312–D319, January 2013. ISSN 0305-1048. doi: 10.1093/nar/gks1063. URL `https://doi.org/10.1093/nar/gks1063`.

[36] Maxim V. Shapovalov and Roland L. Dunbrack. A Smoothed Backbone-Dependent Rotamer Library for Proteins Derived from Adaptive Kernel Density Estimates and Regressions. *Structure*, 19(6):844–858, June 2011. ISSN 0969-2126. doi: 10.1016/j.str.2011.03.019. URL `https://www.sciencedirect.com/science/article/pii/S0969212611001444`.

[37] Longxing Cao, Brian Coventry, Inna Goreshnik, Buwei Huang, William Sheffler, Joon Sung Park, Kevin M. Jude, Iva Marković, Rameshwar U. Kadam, Koen H. G. Verschueren, Kenneth Verstraete, Scott Thomas Russell Walsh, Nathaniel Bennett, Ashish Phal, Aerin Yang, Lisa Kozodoy, Michelle DeWitt, Lora Picton, Lauren Miller, Eva-Maria Strauch, Nicholas D. DeBouver, Allison Pires, Asim K. Bera, Samer Halabiya, Bradley Hammerson, Wei Yang, Steffen Bernard, Lance Stewart, Ian A.

Wilson, Hannele Ruohola-Baker, Joseph Schlessinger, Sangwon Lee, Savvas N. Savvides, K. Christopher Garcia, and David Baker. Design of protein-binding proteins from the target structure alone. *Nature*, 605(7910):551–560, May 2022. ISSN 1476-4687. doi: 10.1038/s41586-022-04654-9. URL `https://www.nature.com/articles/s41586-022-04654-9`. Number: 7910 Publisher: Nature Publishing Group.

[38] Andrew Leaver-Fay, Michael Tyka, Steven M. Lewis, Oliver F. Lange, James Thompson, Ron Jacak, Kristian Kaufman, P. Douglas Renfrew, Colin A. Smith, Will Sheffler, Ian W. Davis, Seth Cooper, Adrien Treuille, Daniel J. Mandell, Florian Richter, Yih-En Andrew Ban, Sarel J. Fleishman, Jacob E. Corn, David E. Kim, Sergey Lyskov, Monica Berrondo, Stuart Mentzer, Zoran Popović, James J. Havranek, John Karanicolas, Rhiju Das, Jens Meiler, Tanja Kortemme, Jeffrey J. Gray, Brian Kuhlman, David Baker, and Philip Bradley. ROSETTA3: an object-oriented software suite for the simulation and design of macromolecules. *Methods in Enzymology*, 487:545–574, 2011. ISSN 1557-7988. doi: 10.1016/B978-0-12-381270-4.00019-6.

[39] LLC Schrödinger and Warren DeLano. PyMOL, May 2020. URL `http://www.pymol.org/pymol`.

[40] J. D. Hunter. Matplotlib: A 2D graphics environment. *Computing in Science & Engineering*, 9(3):90–95, 2007. doi: 10.1109/MCSE.2007.55. Publisher: IEEE COMPUTER SOC.

[41] Michael Waskom, Olga Botvinnik, Drew O'Kane, Paul Hobson, Saulius Lukauskas, David C. Gemperline, Tom Augspurger, Yaroslav Halchenko, John B. Cole, Jordi Warmenhoven, Julian de Ruiter, Cameron Pye, Stephan Hoyer, Jake Vanderplas, Santi Villalba, Gero Kunter, Eric Quintero, Pete Bachant, Marcel Martin, Kyle Meyer, Alistair Miles, Yoav Ram, Tal Yarkoni, Mike Lee Williams, Constantine Evans, Clark Fitzger-

ald, Brian, Chris Fonnesbeck, Antony Lee, and Adel Qalieh. mwaskom/seaborn: v0.8.1 (September 2017), September 2017. URL `https://doi.org/10.5281/zenodo.883859`.

[42] Pauli Virtanen, Ralf Gommers, Travis E. Oliphant, Matt Haberland, Tyler Reddy, David Cournapeau, Evgeni Burovski, Pearu Peterson, Warren Weckesser, Jonathan Bright, Stéfan J. van der Walt, Matthew Brett, Joshua Wilson, K. Jarrod Millman, Nikolay Mayorov, Andrew R. J. Nelson, Eric Jones, Robert Kern, Eric Larson, C. J. Carey, İlhan Polat, Yu Feng, Eric W. Moore, Jake VanderPlas, Denis Laxalde, Josef Perktold, Robert Cimrman, Ian Henriksen, E. A. Quintero, Charles R. Harris, Anne M. Archibald, Antônio H. Ribeiro, Fabian Pedregosa, and Paul van Mulbregt. SciPy 1.0: fundamental algorithms for scientific computing in Python. *Nature Methods*, 17(3): 261–272, March 2020. ISSN 1548-7105. doi: 10.1038/s41592-019-0686-2. URL `https://www.nature.com/articles/s41592-019-0686-2`. Number: 3 Publisher: Nature Publishing Group.
